# Physically Interpretable Performance Metrics for Clustering

**DOI:** 10.1101/2024.11.16.623959

**Authors:** Kinjal Mondal, Jeffery B. Klauda

## Abstract

Clustering is a type of machine learning (ML) technique which is used to group huge amounts of data based on their similarity into separate groups or clusters. Clustering is a very important task which is nowadays used to analyze the huge and diverse amount of data coming out of molecular dynamics (MD) simulations. Typically, the data from the MD simulations in terms of their various frames in the trajectory are clustered into different groups and a representative element from each group is studied separately. Now a very important question coming in this process is what is the quality of the clusters that are obtained. There are several performance metrics that are available in literature like Silhouette index and Davies-Bouldin Index that are often used to analyze the quality of clustering. However, most of these metrics focus on the overlap or the similarity of the clusters in the reduced dimension that is used for clustering and do not focus on the physically important properties or the parameters of the system. To address this issue, we have developed two physically interpretable scoring metrics that focus on the physical parameters of the system that we are analyzing. We have used and tested our algorithm on three different systems (1) Ising model, (2) peptide folding and unfolding of WT HP35, (3) a protein-ligand trajectory of an enzyme and substrate and (4) a protein-ligand dissociated trajectory. We show that the scoring metrics provide us clusters that match with our physical intuition about the systems.

## 1. Introduction

Biomolecular data, especially data generated from molecular dynamics (MD) simulations can be huge and require complex coding to interpret. On top of that, MD simulations often must deal with slow time scales as traditional unbiased MD can reach only ms time scales for all-atom resolution of reasonably sized systems (100k atoms). Although initial ideas on Machine Learning (ML) started in the mid-1900s, it is only recently that computer chips are at the level to use ML to analyze and interpret the huge volume of data coming from MD simulations, as well as enhance MD to reach longer time scales^1–5^.

ML techniques that are used in analyzing and enhancing biomolecular simulations can be classified into two primary forms. i.e., supervised and unsupervised learning ^2–4^. Supervised ML is a form of ML in which data and labels are used to train the ML model. Through this training hidden correlations between the input data and the output labels are learned, which can be used to make further predictions on input data for which labels are not known ^6, 7^. On the other hand, unsupervised ML is a technique which can be used to learn hidden features from just the data itself and can help analyze the system better^1, 8^. In addition to unsupervised and supervised ML, there exists other forms of ML like Generative AI (Generative Artificial Intelligence) that uses techniques both from supervised and unsupervised ML that can be used to learn features about the system of interest at arbitrary temperatures for which simulations/experiments are hard to carry out by using data for which simulations were already carried out^9, 10^.

Unsupervised ML comes in two different forms, i.e., dimensionality reduction and clustering. Data from biomolecular simulations can be huge and multidimensional in nature. Dimensionality reduction is a technique that can be used to project huge multidimensional data into a smaller dimensional space that can be easier to analyze and comprehend^11–14^. On the other hand, clustering is a technique that can be used to group large amounts of data based on similarity into separate groups or clusters. Generally, members of the same cluster are very similar to each other and usually one member of each cluster is studied separately^15–17^. Dimensionality reduction and clustering often complement each other in the way that a real biomolecular system can be described in terms of its order parameters (OPs) which are generally multidimensional in nature. These multidimensional OPs are projected to lower dimensions and this low dimensional representation (generally 1D in dimensional) is further feed into clustering algorithms which further subdivides the data into separate groups or clusters^18–22^.

Dimensionality reduction and clustering algorithms methods have been used in recent past to analyze many biomolecular systems in the recent past. Some recent systems of interest that have been of focus in clustering method development have been peptides/protein systems that show folding and unfolding in relatively short time scales, as well as protein-ligand trajectories. Ghorbani et al. developed a method of using Gaussians in the latent space to cluster multiple protein folding trajectories^18^. Cai et al. developed a method of using coarse focus knob to identify conformations with low free energy and fine focus knob to distinguish the various conformations between these multiple free energy states and finally cluster the protein folding trajectories^21^. Wang et al. used their method of State Predictive Information Bottleneck for clustering protein folding trajectories into distinct states^23^. Chen et al. developed a clustering method to improve upon KMeans++ algorithm by initializing the best centroids for clustering^24^. Bray et al. developed a clustering method based on Principal Component Analysis and then training a XGBoost model on top of that to identify unbinding paths of ligand bound to a protein. They also clustered trajectories according to their pairwise mean Euclidean distance by employing a neighbor-net algorithm^25^. Ray et al. developed a dynamic-time wrapping algorithm to classify protein ligand dissociation trajectories^26^. Wolf et al. developed a method to cluster protein ligand dissociation trajectories based on a community detection algorithm that requires the definition of only a single parameter^27^.

An important question while clustering data is often what the quality of clustered data is. This is extremely important as this helps in determining the quality of clustered data and will help in determining the optimal number of clusters, the clustering algorithm to use, as well as the optimal order parameters necessary to describe the system^28^. There are different clustering metrics that people often have used in the past. Some popular clustering metrics include the V-Measure score^29^, Calinski-Harabasz index^30^, Silhouette index^31^, and Davies-Bouldin Index^32^ all of which uses the separation of the various clusters as well as the similarity of the members of each of cluster to determine the clustering quality^23, 24^ . In addition to this, metrics like Shanon Entropy can also be used to quantify the populations of the various clusters^23^. However, all these metrics lack clear physical interpretation as they do not focus on the microscopic/macroscopic physical properties or parameters of the system which must be well separated as well for the quality of the clustering to clear or well defined.

In this paper, we introduce two different physical property-based scoring scales to analyze or quantify the quality of clustering data. Our paper is organized in the following manner. In section 2.1, we discuss the various methods that we use for clustering our data along with dimensionality reduction, and in sections 2.2 and 2.3 we discuss the two different metrics that we use for scoring our clustering results. In section 3, we apply our clustering methods on three different simulated systems-(1) Ising model system^33^, (2) a 398-ms long simulation of WT-HP35 protein folding and unfolding obtained from DESRES (DE Shaw Research)^34^ and (3) a protein-ligand unbiased MD simulation trajectory from our own simulations^35^.

## 2. Methods

### 2.1 General overview of Clustering, Dimensionality Reduction and the different clustering algorithms used

Clustering is a method by which different members of an ensemble are grouped into distinct categories or groups. To perform clustering especially on biomolecular systems, a suitable 1D representation of the system must be created based on which we can perform clustering. Generally, it is very hard to find suitable order parameters that can accurately describe a system in 1-dimension. Because of this reason, biomolecular systems are generally represented in terms of multidimensional order parameters on which dimensionality reduction techniques are applied to accurately get a 1D representation of this multidimensional order parameter space. Once a 1D representation of this order parameter space is obtained, we can run our clustering algorithms. In our paper, we use a Variational Autoencoder (VAE) for our dimensionality reduction technique^36, 37^.

VAE is often used to compress multidimensional information into a reduced 1-dimension. A general VAE consists of three components: encoder, latent space and decoder. The input data to the VAE is a complex multidimensional representation of the order parameters that a researcher determines to best represents the input data e.g., the biomolecular conformation in our case^38–40^. This complex representation of the biomolecule is fed into the encoder and projected into a low dimensional latent space. Next, the decoder tries to reconstruct the complex dimensional representation of the original input and in turn learns a low dimensional representation that best resembles the original high dimensional input data. The only thing that differentiates a VAE from an autoencoder is that the autoencoder samples directly from the latent projections to feed into the decoder whereas the VAE learns to represent the latent dimension as a Gaussian with its mean and variance being the learnable parameters of the Gaussian and after that that it samples from this Gaussian to feed the decoder.

So, once we have found suitable order parameters that best describe the system, a VAE is trained to represent the multidimensional order parameter space into a latent 1D representation. Now this latent 1D representation of our simulation trajectory time frames are fed into the following clustering algorithms to cluster our simulation trajectory into distinct clusters or states^18, 22^.

i. K-Means^41–43^: Here data points are assigned to K clusters based on the distance between the points and their distances from the center of the clusters. The cluster centroids initially are randomly initialized. Once each data point is assigned a cluster based on their minimum distance to the centroids, new cluster centroids are found out and reinitialized. This process is run iteratively until a solution converges.
ii. Agglomerative^44–46^: In Agglomerative clustering, each observation starts in its own cluster and pairs of clusters are merged as we gradually form a tree of clusters. In this method, two clusters are merged based on the distances between them and a linkage criterion as well which specifies which clusters to merge.
iii. BIRCH^47–49^: BIRCH (*balanced iterative reducing and clustering using hierarchies*) is a two-step unsupervised clustering algorithm in which a summarization of the original data sets in smaller dense regions called Clustering Feature (CF) entries is attempted. After forming CF entries, a compact representation of the original dataset is formed by making a tree where each node contains a sub-cluster. This is called as a Tree Clustering Feature.
iv. Gaussian Mixture^50, 51^: In the Gaussian mixture clustering algorithm, each point is attributed to belong to one of the several K Gaussians. Each of these Gaussians are identified through three parameters, mean, variance, and mixing coefficient which denotes the probability of the occurrence of each of the Gaussians. The mean, variance and the mixing coefficient of each of the Gaussians are estimated through an entropy maximization algorithm for the parameters.
v. DBSCAN^52–55^ (Density based spatial clustering based on noise) is a density-based clustering algorithm which separates datapoints into clusters based on the density and their neighborhood to each other. Initially datapoints are assigned to three distinct categories (i) core points if each of these data points have at least a minimum number of samples (min-points) within a specified neighborhood (ε) of the chosen sample. (ii) border points if each of these data points are not core points but still, they lie within some ε neighborhood of a core points and (iii) outliers i.e. points that neither belong to a core point or a border point. Through an iterative process, nearby core points with their neighbors are gradually merged with each other to form a single cluster until they can be merged no more. In the next step, the next set of core points are merged with each other to form another cluster. This process is repeated until we are just left with outliers or noise.

We have chosen to study three different model systems for our work. The details of the order parameters that we have chosen to get an initial multidimensional representation of our systems are described below in detail. Once we have clustered our systems using the various algorithms, we proceed forward to analyze the clustering results using the two different metrics that are discussed next.

### 2.2 A Detailed Workflow for our Scoring Metric-1

The purpose of any basic clustering algorithm should be to cluster the biomolecular simulation trajectory of interest into clear and separate well defined groups. Now a very important question related to this is how we can classify clusters as well-defined groups. A simple way of thinking is that an object of interest is well defined only in terms of its physical properties. For example, a group of used cars can be defined in terms of their most important physical properties, e.g., cost, mileage, color, manufacturer, and year built. This way of thinking forms the basis for our two different scoring metrics.

For our first scoring metric, our idea is that even though we do not have ground truth labels in the case of clustering algorithms, we can use property-based ground truth labels to develop a clustering quality metric.

In detail, let us say we have clustered our objects of interest into k clusters. After doing this, we can assign cluster labels to each frame of our trajectory L = {L_1_, L_2_, L_3_, …., L_n_}, where N = {1, 2, 3, …., n} denotes the frames in the trajectory with a total of n frames. Now say that the biomolecular simulation trajectory can be defined in terms of an important set of properties P_imp_. Then for each element p that belongs to set P_imp_, we can find out individually the property values P = {P_1_, P_2,_ P_3_, …., P_n_} for each frame in the trajectory. After collecting these set of properties, we can then do a clustering on these set of properties themselves into k clusters. Once we have clustered these properties, we can assign cluster labels to these properties PL= {PL_1_, PL_2,_ PL_3_, …., PL_n_}. We set PL as our ground truth labels for our major property P. Now our scoring metric consists of two terms. The first term is the Silhouette score^31^ label (SC) for the property labels themselves which can give an insight into how clear/well separated the clusters have been formed for the property p as well as similarity for each member of the cluster.

Note that since we are using a property-based pseudo ground truth label for our clustering metric, this term acts as a penalty for using ground truth labels for building our metric. Ultimately, a lower score will reflect poor clustering on these pseudo ground truth labels themselves. Our second term consists of a term F that can give a score on how close our original cluster labels L are close to the ground truth labels PL. To do so, we try to form a one-one mapping between our ground truth labels PL and our original cluster labels L. To do a one-one mapping, we iterate through each of the clusters 1, 2, 3, …, k in the pseudo ground truth labels PL. For each cluster j in these pseudo ground truth labels, we gather the frame numbers in PL that are equal to j. A subset F_j_⊆ N are the frame numbers in PL that are equal to j. Then from our original cluster labels L, we find the subset L_fj_ ⊆ L for all elements (frame numbers) f in F_j_. Now, we find the most frequent cluster number m that occurs in L_fj._ Now, this number m from L_fj_ (original cluster labels) maps to j from PL (ground truth labels).

Based on this mapped number m from j, we can define two different fractions-f_pmj_that calculates the probability of occurrence of this number m in L_f_*_j_*, as well as the probability of occurrence of F_j_ itself in N equals f_p_*_j_*. So based on this j to m mapping, we can define a product f*_j_*= f*_pmj_* x f*_pj_* that gives a weighted score of how well aligned the mapped cluster number m is with respect to the pseudo ground truth cluster j by taking into the consideration the probability of occurrence of the ground truth cluster label j itself. In addition to this, there can be some cases where multiple j’s from the pseudo ground truth labels are mapped to the same m. Note that in these cases, we set only to retain the max f_j_ and set all others to 0. Now we can sum over all the j’s in the ground truth labels to get our final term F. Hence, our final scoring function is SCxF. For clearer and well aligned original cluster labels L with respect to the pseudo ground truth labels and for clear one-one mapping between m and j, F will go to 1. Also, for very clear and well-defined ground truth labels, the SC will go to 1 because of the definition of the Silhouette-Score itself. So, our overall score S_p_ would go to 1 in case of ideal and most efficient clustering. A tabular description of our algorithm is given below.

#### Algorithm-1

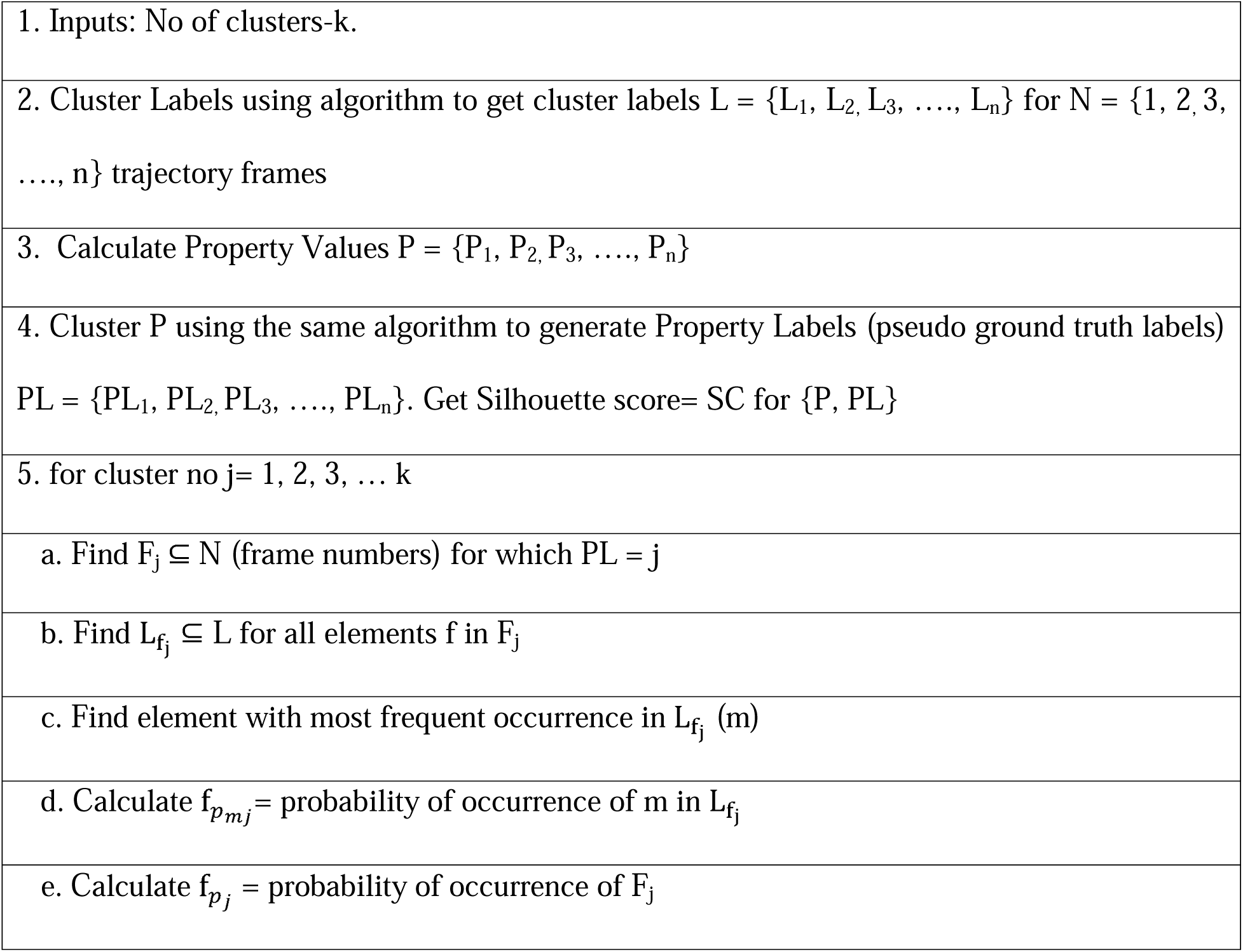
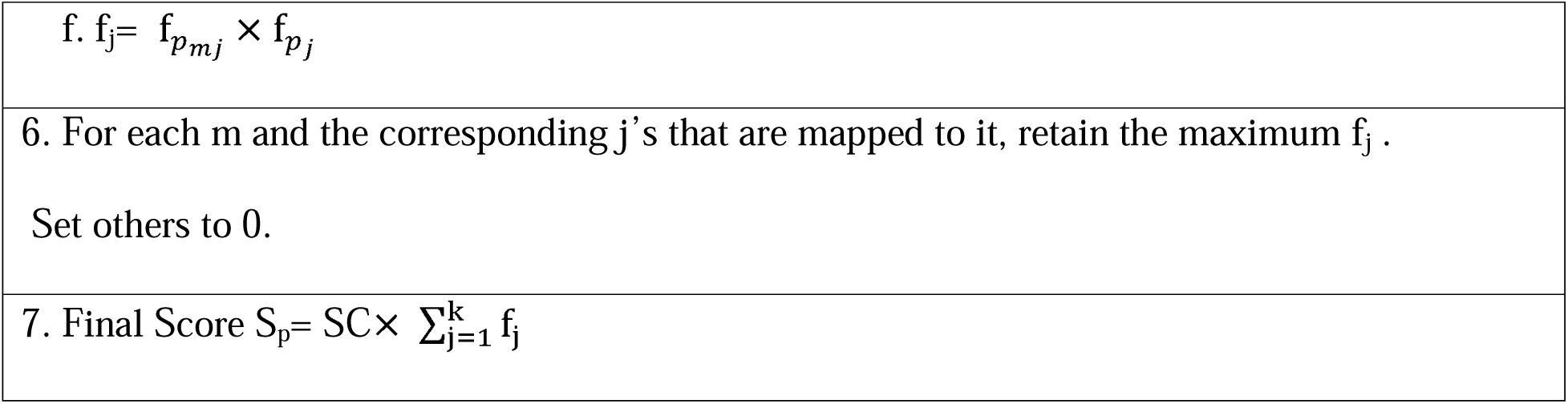

### 2.3 A Detailed Workflow for our Scoring Metric-2

In this scoring metric, we do not consider the ground truth labels associated to each property. Instead, we consider the properties belonging to the elements of each cluster from the cluster labels (we call them sub-properties) and verify if those sub-properties can form multiple clusters among them (we call them sub-clusters) or not. The smaller number of sub-clusters these sub-properties form, the more there is an increment to the score.

In detail, we describe our second physically interpretable scoring metric. Consider, we have again clustered our object of interest into k clusters. Then, we assign cluster labels to each frame of our trajectory L = {L_1_, L_2,_ L_3_, …., L_n_} where N = {1, 2, 3, …., n} denotes the frames in the trajectory with a total of n frames. Next, we choose an important set of properties P_imp_ and we study each property p amongst them. Now for each such property p, we again find out the property values P = {P_1_, P_2,_ P_3_, …., P_n_}. After this, we iterate through each of our clusters j=1, 2, 3, …, k and find frame numbers in L that are equal to j. Consider a subset F_j_⊆ N are the frame numbers in L that are equal to j. Now, we select properties P_fj_ from P, for each element/frame number f in F_j_ which we call sub-properties. Once we get these sub-properties P_f_*_j_*, we try to cluster these properties with n=2,3,4,5,..k clusters and use a Silhouette-score metric to see how many clear clusters we can form (sub-clustering) which can get us the probability of occurrence of the most dominant sub-property P_sub_*_f_* within these set of sub-properties P_f_*_j_* . The maximal of these Silhouette-scores (sc-max) will give us the optimal number of clusters for our sub-properties. However, we would also want to check if sub-clustering on these sub-properties would make any sense or not. In this case, we set a silhouette-cutoff (sc-cutoff) to set a bound on the maximal number of clusters that we can allow to be formed. If sc-max<sc-cutoff, we can say that there is no further sub-clustering possible on the sub-properties and P_sub_ equals the whole _j_ set of sub-properties P_f_*_j_*. Else, we look at the optimal number of clusters through sc-max and look at the most majorly occurring sub-cluster label and declare P_sub_ as having the most major _j_ dominant sub-property. Now, that we have done this, we can define two different fractions-f_p_*_j_* that denotes the probability of occurrence of F*_j_* in N, as well as another fraction f_psub*j*_ that denotes the probability of occurrence of the most dominant subproperty P_sub_ within this subset _j_ F_j_. We can now take their product f_j_ =(f_p_*_j_* x f_psub*j*_) to denote the weighted score of how majorly dominant each dominant sub-property P_sub_ is. After this, we can sum over the product f*_j_* to get _j_ our final scoring term S_p_.

#### Algorithm-2

|  |
| --- |
| 1. Inputs: No of clusters-k. Silhouette-cutoff (sc-cutoff) |
| 2. Cluster Labels using algorithm to get cluster labels $L = \{L_1, L_2, L_3, \dots, L_n\}$ for $N = \{1, 2, 3, \dots, n\}$ trajectory frames |
| 3. Calculate Property Values $P = \{P_1, P_2, P_3, \dots, P_n\}$ |
| 4. for cluster_no $j = 1, 2, 3, \dots k$ |
| a. Find $F_j \subseteq N$ (frame numbers) for which $L = j$ |
| b. Find $P_{f_j} \subseteq P$ (*sub – properties*) for all elements $f$ in $F_j$ |
| c. Sub-cluster $P_{f_j}$ into 2, 3, 4, 5, ... , $k$ clusters using the same algorithm. |
| d. Get silhouette-scores for 2, 3, 4, 5, ... , $k$ sub-clusters. |
| e. Find the maximal silhouette-score (sc-max). |
| f. If $sc-max < sc-cutoff = 0.30$ (chosen here), no further sub-clustering possible: $P_{sub_{f_j}} = P_{f_j}$<br>and goto h. |
| g. Look at optimal clusters through sc-max and the most dominant cluster = $P_{sub_{f_j}}$ |
| h. Calculate $f_{p_j}$ = probability of occurrence of $F_j$ |
| i. Calculate $f_{p_{subj}}$ = probability of occurrence of $P_{sub_{f_j}}$ |
| j. $f_j = f_{p_j} \times f_{p_{subj}}$ |
| 5. Final Score $S_p = \sum_{j=1}^k f_j$ |

Now that we have discussed our scoring functions, an important factor to decide is which properties are chosen for use in the scoring function. It is noteworthy to mention that we might decide to look at multiple properties before deciding on the best algorithm/number of clusters to select. To do so, we may decide to look at multiple properties and from physical intuition decide to give more importance to some properties than the others. From this, we can construct an overall scoring function, i.e.

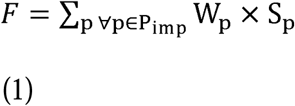

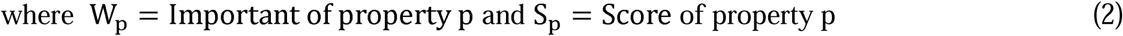

where we can set up importance of each property p as to our physical intuition. Trivially, summation over weights of all properties should be 1. This means

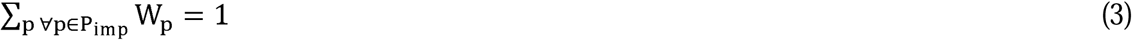

Now that we have described our algorithm in detail, we will extend this approach to clustering methods that do not include the number of clusters as an input parameters. Specifically the DBSCAN algorithm will be our focus and it takes in input the neighborhood radius (ε) as well as the minimal number of points (min-points). A modified version for our algorithm that can input ε as well as min-points is provided in the Supplementary Information (pages S2-S6). In fact, a major challenge in using the DBSCAN algorithm is to determine the values for min-points and ε^56^. Generally, min-points are taken to be the twice as that of the spatial dimensionality and ε is set as the knee point in a k-distance graph^57, 58^. However, in many cases the actual dimensionality of the space is not known or well defined. In these scenarios choosing min-points and ε especially becomes a cumbersome task. In our case, we show that using our physically interpretable methods the optimal min-points and ε matches with the physical intuition for our systems.

Now that we have described our algorithm in detail, we can proceed forward to apply our methods to our three chosen systems in detail: (A) Monte Carlo simulations of an Ising model system, (B) MD simulations of a HP-35 protein folding and unfolding, and (C) our past MD simulations of Smlt1473 enzyme bound to various substrates^35^.

## 3 Results and Discussions

### 3.1 Ising Model System

The Ising Model is a simple well studied 2D physical system consisting of interacting spins. The spins are allowed to take two possible values a-=1 or a-=-1. The spins interact with each other through an interaction strength J, and the Hamiltonian is given by H=J∑_<i,j>_ a-_i_ a-_j_ . It is one of the simplest systems that can show different energy states or phases as it shows a phase transition from a ferromagnetic to a paramagnetic phase as we gradually increase the temperature. At very low temperatures, the spins are mostly aligned to each other in this system as this tends to reduce the energy of the system and so the overall magnetization/spin is 1 and the overall energy of the lattice/site is -2 J. As we gradually increase the temperature, the spins gradually start misaligning with each other and after a particular temperature, the overall magnetization of the lattice per site goes to 0 and the overall energy per lattice site also increases to 0. Apart from this, there are also two other parameters of interest (a) heat capacity and (b) susceptibility which capture/s the fluctuations of the Ising spin lattices at a particular temperature. These two parameters show a divergence as a function of temperature as we gradually change the temperature from a low value to a high value.

We have studied the Ising model using two different approaches. In our first approach, a general clustering of different Ising model samples collected at various temperatures are calculated. Specifically, we considered a 30×30 Ising model and collected 9000 Ising model samples at a particular temperature for 20 different temperatures ranging from T=1 to T=5 (assuming the Boltzmann constat value to take a value of 1). This creates a 180000×30×30-dimensional matrix for clustering. To make a suitable representation of the 30×30 Ising model, we trained a VAE on this 180000×30×30 vectors to create a linear representation of this 30×30 spin lattice and converted it to a 180000×200-dimensional vector. Now using this reduced low dimensional system, we trained our clustering algorithms K-Means, Agglomerative, BIRCH, and Gaussian Mixture and we attempt to form 2, 3, 4, 5, and 6 clusters using each algorithm.

For our second clustering approach, we did a time-series based clustering for the Ising model trajectories. To do this, 200 equally spaced temperatures between 1 and 5 are used and for each temperature 900 samples are considered. Again, we trained a VAE on these 180000×30×30-dimensional matrix and based on the trained VAE, we generated 180000×200 dimensional 1D representations of the Ising model consisting of 900 representations for each temperature. Once, we generate 900 one-dimensional representations for each temperature, we perform an average-pooling operation to get an ensemble representation for each temperature. We can perform an average pooling operation as the Ising model is in equilibrium at each temperature and hence the time-series Ising model simulated data effectively turns into time-independent data due to ergodicity. For more advanced time series data, which is not in equilibrium, like protein ligand dissociation trajectories, a more advanced clustering approach may be required^26, 27, 59^.

In figures 1 and 2, we represent the properties of the different Ising model samples used for clustering. Energy and magnetizations are in the y axis respectively for figures 1 and S1 and the temperature is on the x axis and the different cluster samples are represented using different colors. In figures S2 and S3, we cluster the properties themselves and look at the ground truth labels which we use a lot for our metrics defined in our scoring metric-1. In figure 2, we look at different scores assigned by our two different scoring metrics. We see that for our scoring metric-1, we get a maximum score with two clusters for properties magnetization and energy which matches with the physical intuition that finding the order and disordered states should have been done by the different clustering models. Also, since our clustering property-based scoring metric-1 considers pseudo ground truth cluster labels (figures S2 and S3), this matches on physical observation that only for two clusters the cluster labels for figures 1 and S1 align the most with figures S2 and S3(figure 1 aligning with figure S2 and figure S1 aligning with figure S3). Now looking at scoring metric 2 from figure 2, we find that the optimal number of clusters are found for number of clusters=2 for the energy-based scoring metric-2 and number of clusters=3 for the magnetization-based scoring metric-2 which is a mixture between the ordered and the disordered state by agglomerative clustering. There are more clusters predicted by the magnetization-based scoring metric-2 because magnetizations for the disordered states have much more spread than the energy for the disordered states which is very well classified by the scoring metric-2. Now on comparing scoring metrics-1 and 2, we see that scoring metric-1 is able to provide us with large separation between the various clusters, whereas metric-2 provides less separation. This is probably because in metric-1 we are trying to do a one-one mapping between our pseudo ground truth labels and our cluster labels with a penalty of 0 score is when there lacks a one-one map between our cluster labels and pseudo ground truth labels (step 6 in algorithm-1), as well bringing in a penalty term for bad pseudo ground-truth labels. So, we must choose between scoring metric-1 vs metric-2 according to our needs for clustering our data. If we are interested in getting just the major states and are not interested in intermediate states, we might decide to look at scoring metric-1 while if we do want to get intermediate states, we should decide on scoring metric-2.

**Figure 1:**
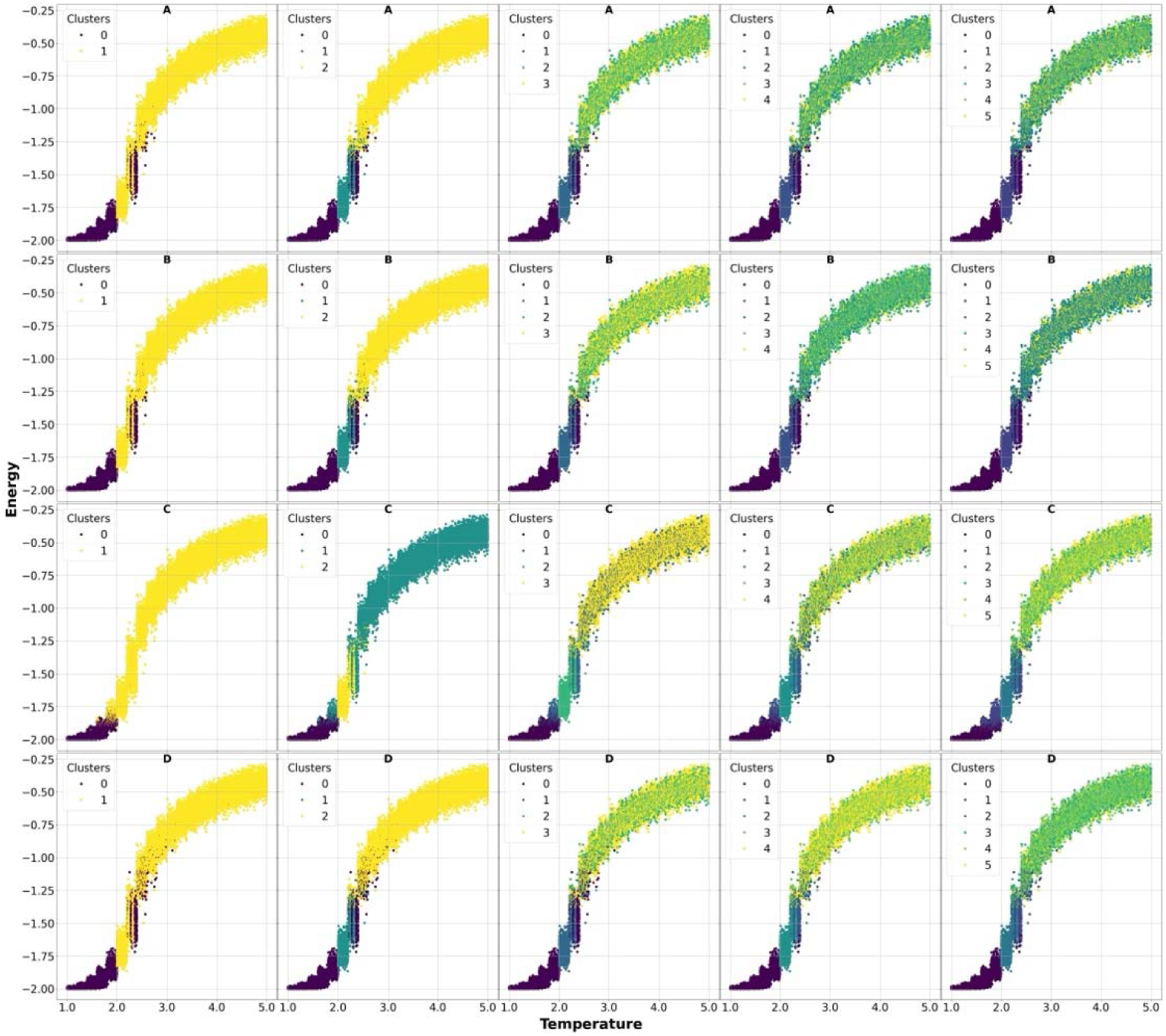
Energy properties of the different Ising model samples used for clustering using our first approach. Different clusters are marked using different colors. A, B, C, and D sets denotes our 4 clustering algorithms K-Means, Agglomerative, BIRCH, and Agglomerative. Each column respectively denotes 2,3,4,5, and 6 clusters.

**Figure 2:**
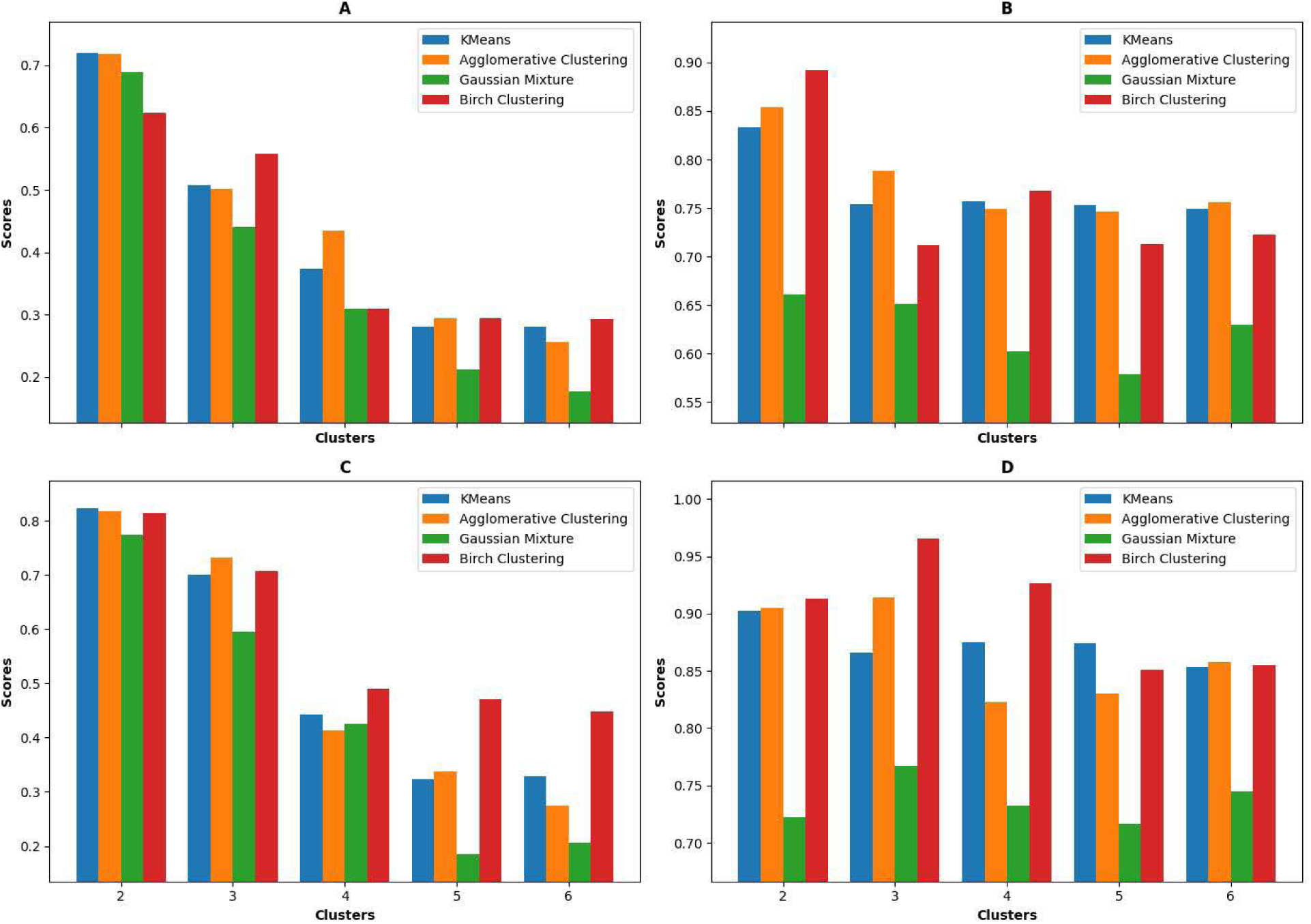
The different scores obtained by our scoring metrics (Scoring metric-1 and Scoring Metric-2 respectively) on energy (A) and (B) and scoring metrics (Scoring Metric-1 and Scoring Metric-2 respectively) on magnetization (C) and (D).

Finally, based on our overall scoring function defined in equations 1-3 in the simplest case we might decide to give equal importance to both the properties magnetization and energy. This would give us 2 clusters both in cases of scoring metrics-1 and -2 like we have shown in figure S4.

In addition, our second clustering approach based on the time series data of the Ising model is shown in figures 3 and S5. We see from those figures that the region below the transition temperature is majorly a particular cluster for number of clusters >3 whereas the region above the temperature is another cluster. Only for number of clusters=2, do we see that most of the regions falling in the same cluster except for the ones near the critical point, which is what we want looking at the perspective from clustering with respect to the heat capacity and the susceptibilities as heat capacities and susceptibilities of an Ising model are relatively the similar throughout the temperature scans except for the regions near the critical point, where we expect more clusters.

**Figure 3:**
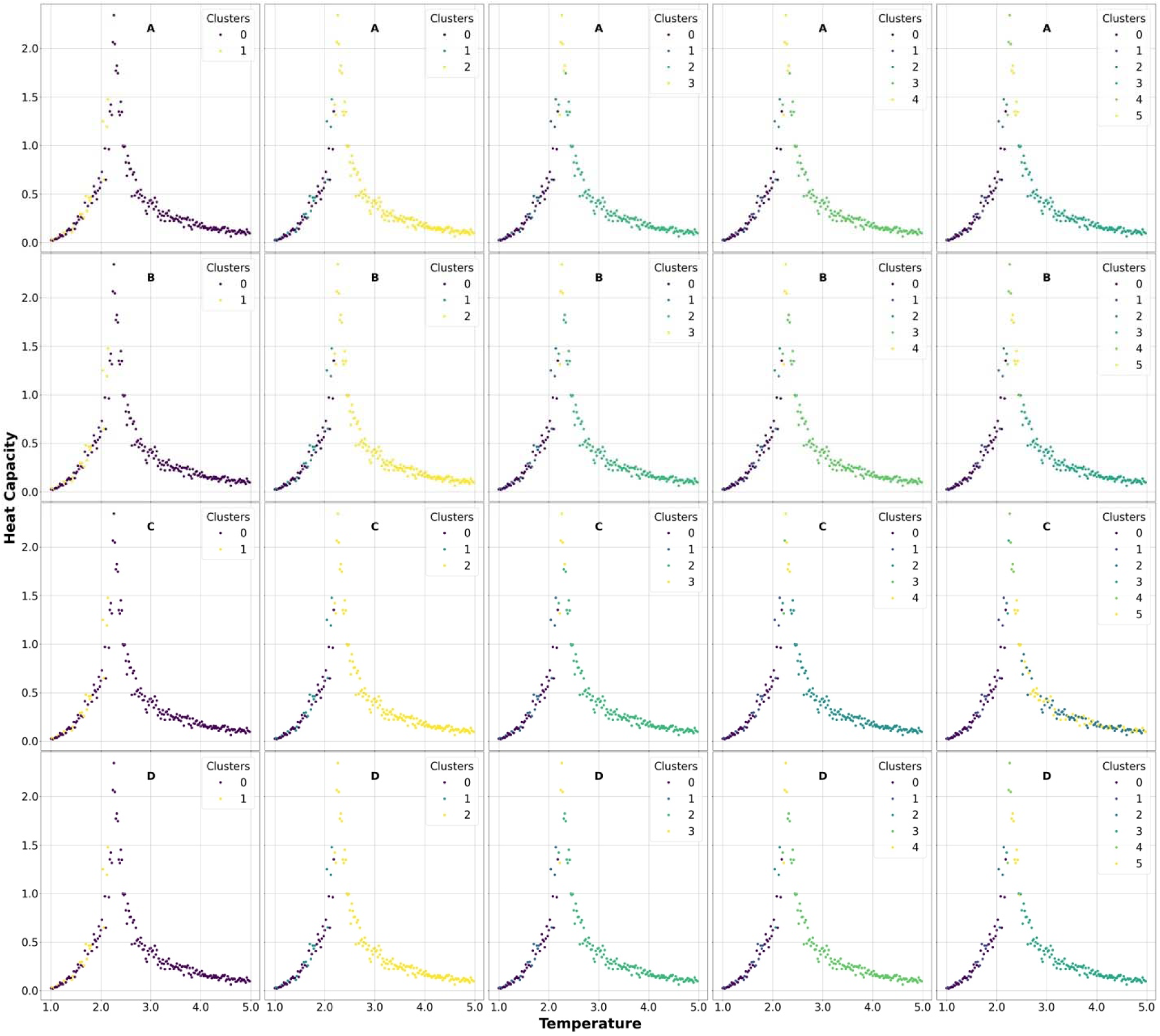
Heat Capacity properties of the different Ising model samples used for clustering using our first approach. Different clusters are marked using different colours. A, B, C, and D sets denotes our 4 clustering algorithms K-Means, Agglomerative, BIRCH, and Agglomerative. Each column respectively denotes 2,3,4,5, and 6 clusters.

This same fact is highlighted by our clustering scoring metrics majorly in figure 4. In figures S6 and S7, we present the ground truth labels used for clustering using heat capacity and susceptibility using scoring metric-1. For figure 4B and 4D, where we also observe less scoring difference the occurrence of 2, 3 or 4 clusters because the right side and the left side of the transition temperatures are mostly in different clusters and the scoring metric-2 does not penalize for this whereas the scoring metric-1 does penalize leading to the occurrence of a smaller number of clusters in figures 4A and 4C. In addition to this, from scoring metric-1, we are getting 2 major clusters as our optimal for both heat capacity and susceptibility (4A and 4C) whereas from metric-2 (4B and 4D), we are getting 2, 3 or 4 optimal clusters. This again highlights a major difference in the methods of design for our scoring metrics-1 and 2. Scoring metric-1 only highlights the most dominant and major states (i.e. the state away from the critical point in which the heat capacity/susceptibility remains close to 0 and the state near the critical point where the heat capacity/susceptibility diverges). But scoring metric-2 also highlights several intermediate states 3 or 4 from the susceptibility scores, which again some reside in the critical region (figure S5). Also, following up from our previous discussion, we might decide to put equal weights to both heat capacity and susceptibility for constructing our overall scoring function based on scoring functions-1 and 2, which would lead to mostly 2 clusters as shown in figure S8. We note here that these choices of weights are not ad-hoc but rather based on the physical intuition that magnetization might be as important a property as the energy of the ising systems.

**Figure 4:**
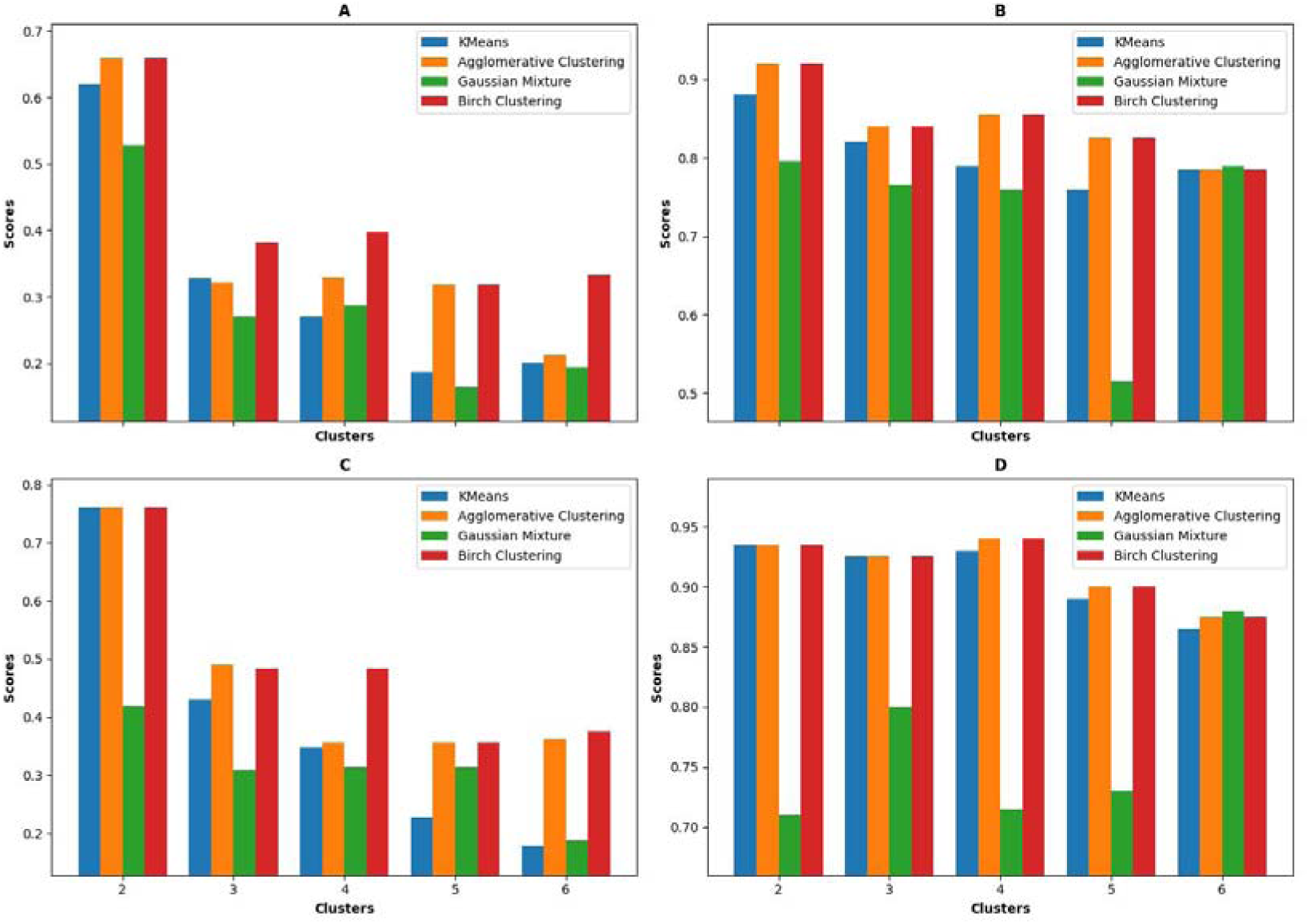
The different scores obtained by our scoring metrics (Scoring metric-1 and Scoring Metric-2 respectively) on heat capacity (A) and (B) and scoring metrics (Scoring Metric-1 and Scoring Metric-2 respectively) on susceptibility (C) and (D).

**Figure 5:**
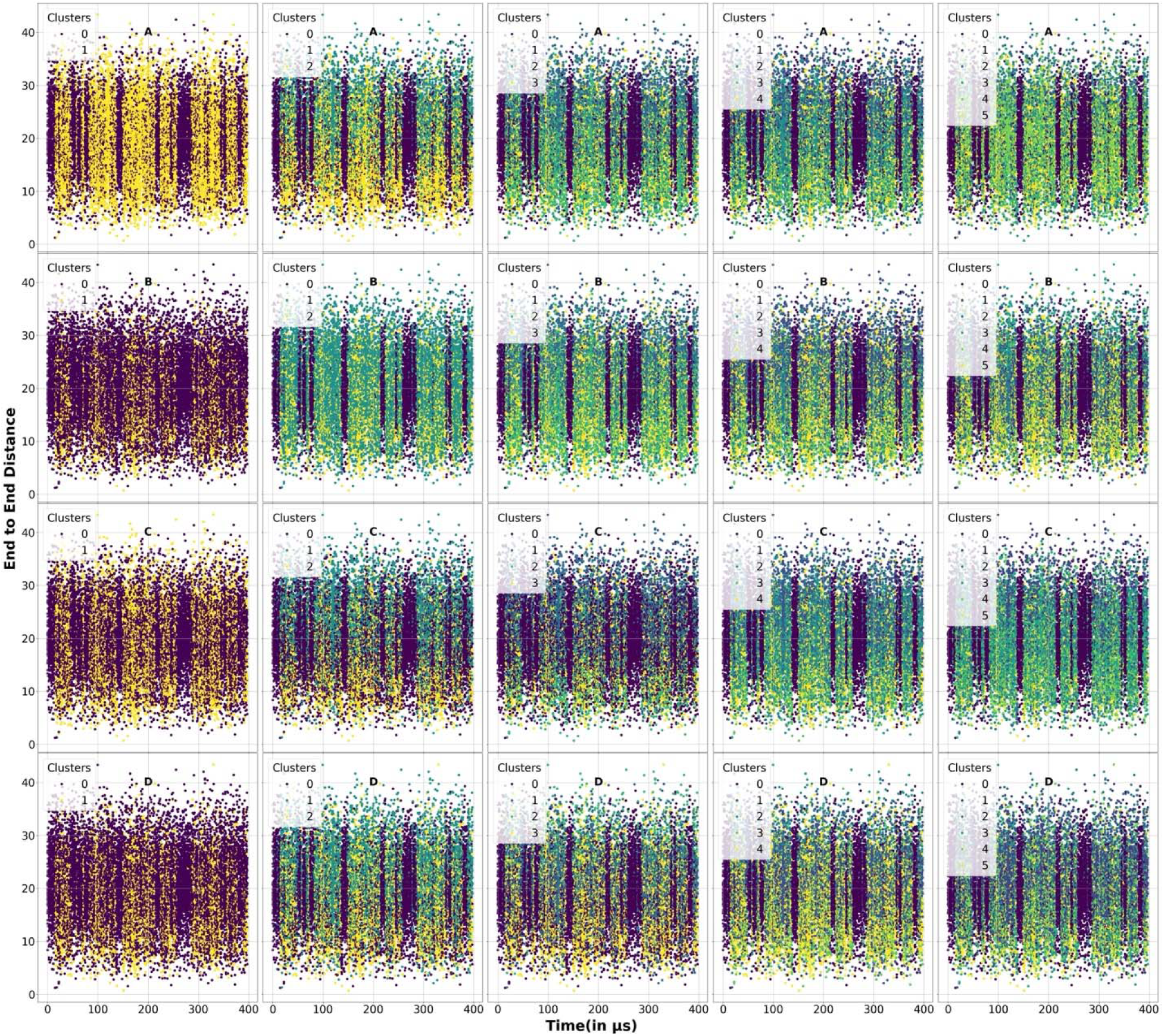
Distance properties of the protein folding simulation trajectories used for clustering. Different clusters are marked using different colors. A, B, C, and D sets denote our 4 clustering algorithms K-Means, Agglomerative, BIRCH, and Agglomerative. Each column respectively denotes 2,3,4,5, and 6 clusters.

As our final study involving the Ising model, we applied the DBSCAN algorithm to show that we can correctly identify the optimal min-points and ε which is given by our clusters using our methods. In figures S9 and S10, we show the clustering results obtained by our DBSCAN algorithm for energy and magnetization, respectively. From these plots identified that minpoints=2800, and 3800 with ε=0.6 have the most clear and well separated clusters along with minpoints=800 and ε=0.5. In figure S11, the scores predicted by our modified algorithms for DBSCAN using equal weights on energy and magnetization and show that our algorithms indeed identify min-points=2800 and 3800 along with ε values=0.6 as our optimal scores along with min-points=800 and ε=0.5. Our scoring metric-2 from figure S11B also identifies min-points=800 and ε=0.5 as our optimal clusters which means it also identified some intermediate states from min-points=800 and ε=0.5. Also, we further applied our DBSCAN based metrics on ranking the clusters obtained from heat capacity and susceptibility (figure S12 and S13) and show that we get the optimal scores of min-points=2,10, or 20 with ε=0.5 (figure S14A) or min-points=2,10, or 20 and ε=0.5 or 0.6 (figure S14B). Finally, we would comment that it would be unwise to compare the scores obtained by various clustering algorithms with the scores obtained by the DBSCAN clustering algorithm as DBSCAN takes in different inputs as compared to the other clustering algorithms and hence our scoring algorithms themselves are different.

### 3.2 System WT-HP35 protein

Our next system of interest is the wild type (WT) villin headpiece 35 (HP-35) protein folding and unfolding from the simulations done previously by the DE Shaw Research (DESRES)^34^. Further details of the simulation are given in their paper. For our interest, this protein shows multiple folding and unfolding events for a 398 ms long MD simulation along with some multiple intermediate states.

Now, to convert the raw trajectory into a usable ML, an inter-residue C_a_ distance matrix of the protein which is 35×35 dimensional is selected, we can iterate through all the trajectories of this simulation (choosing every 100^th^ frame) and have obtained a 19875×35×35-dimensional representation of the protein trajectory. Note that since our system is small, it is possible to treat our system exactly. In cases where the protein size becomes extremely large it would be beneficial to consider specialized Collective Variable selection methods^56^. Once we obtain this, we train a VAE on this trajectory to obtain a 19875×200-dimensional representation of the protein trajectory. After we have obtained this representation, we can use the various clustering algorithms to cluster our trajectory into different stable states.

In Figures S15 and 5, we have plotted two different properties important for our clustering metrics (RMSD relative to the NMR structure of the WT-HP35 protein 1YRF^60^ from the protein data bank and distance between the first 5 residues and the last 5 residues of the protein) on the y axis and the time on the x axis. We represent the different clusters using different colors in the plots. In figures S16 and S17 we represent the ground truth labels used for scoring our clustering metric-1. In figure 6, we represent the various scores obtained using our two different scoring metrics. From these scores, we can have an idea about the optimal number of clusters needed for clustering our data. As before when we had discussed for our Ising model, our scoring metric-1 is useful for getting the major stable states from our trajectory if we are not interested in the intermediates and just want a coarse classification of states, which in our protein folding problem is folded versus unfolded. Since our formulation for scoring metric-1 relies on pseudo ground truth labels for RMSD and distances, low RMSD/end-to-end-distances are mapped to a cluster (which physically represents the folded state) and high values of these properties are mapped to a different cluster (which physically represents the unfolded state). Accordingly, from our plots 6A and 6C, we find that the optimal number of clusters are 2 from our scoring metric-1 which can also be physically verified by the fact that RMSD and the distances ground truth labels from figures S16 and S17 align the best with the actual cluster labels in figures S15 and 5 for only 2 clusters.

**Figure 6:**
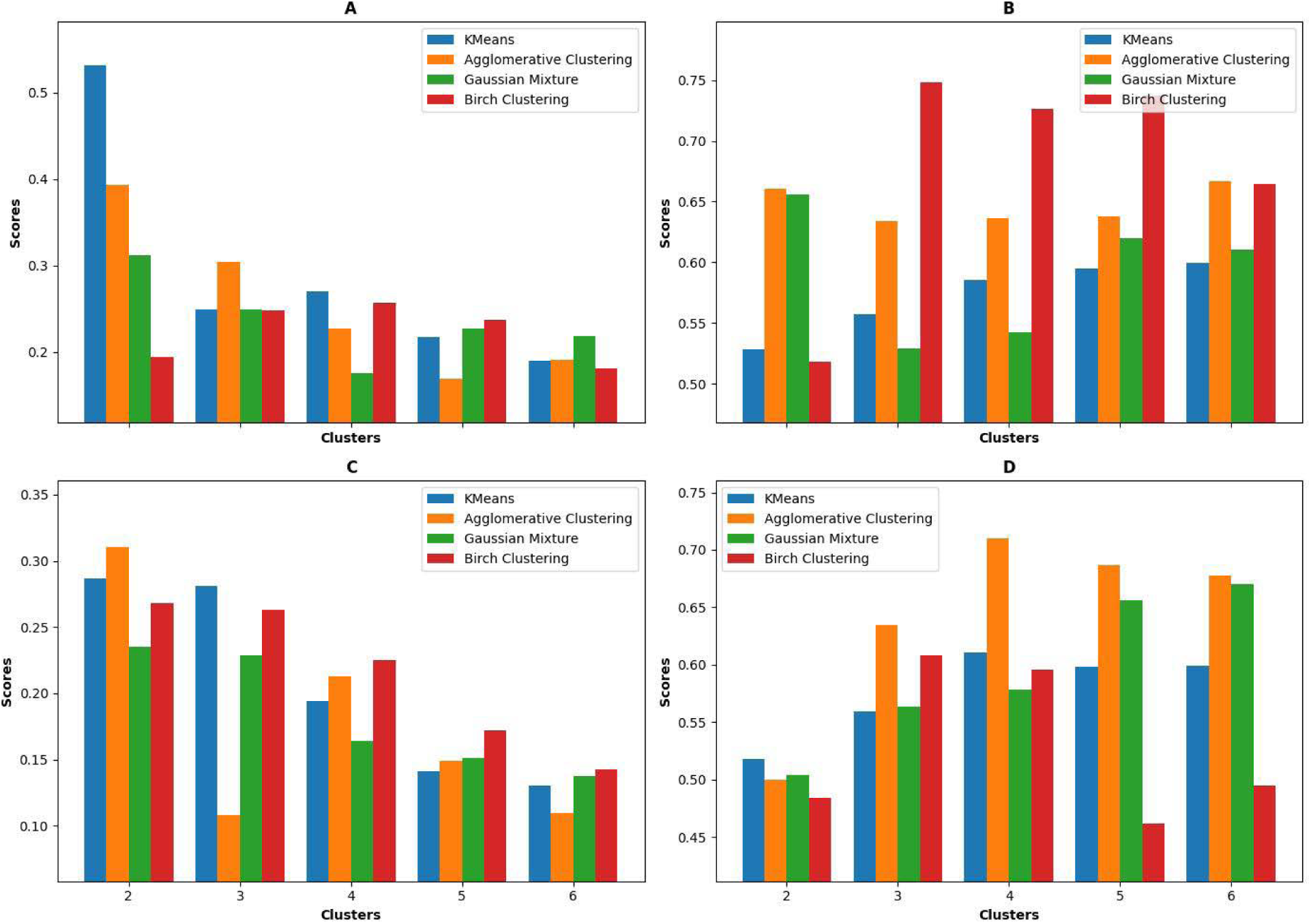
Scoring functions based on RMSD for the protein-folding simulation trajectories used for clustering. (A) and (B) denote the scoring functions-1 and 2 used on RMSD property while (C) and (D) denote the scoring functions-1 and 2 used with the distance property. Different clusters are marked using different colors.

On the other hand, if we are interested in also the intermediate states from our trajectory we must rely on scoring metric-2, which does not rely on ground truth labels but rather tries to minimize the spread of the properties for each of the clusters. For the scoring metric-2 from figure 8, we see that for RMSD Birch Clustering indeed gives 3 distinct clusters as our best metric overall for RMSD and we can see that physically too as there is the least spread overall between the different clusters as there is a very gradual transition from green to yellow in this cluster. We also can see that for distances 4 clusters is the optimal given by Agglomerative as we can physically see that 4 clusters reduce the spread amongst the various clusters. This also means that our clustering metric-2 can find intermediate states apart from the major states. These results of 3/4 optimal clusters are in close quantitative agreement with previous studies^23, 24, 61^. Five clusters were chosen by Wang et al. for constructing a Markov State Model on the simulations on this HP-35 protein ^23^. Chen et al. and Klem et al. chose 4 state models as well in their studies based on physical and chemical intutions^24, 61^

It is also noteworthy to mention that just considering RMSD/end-to-end-distances alone might not be a good measure of our scoring metric. We must consider an overall scoring function F (equation 1). In most practical cases, we might want to put more weight onto the end-to-end-distances property than the RMSD as they end to end distances provide a more direct measurement of how folded/unfolded the protein is. Accordingly, we have put a weight W_rmsd_=0.30 on the RMSD scores and a weight W_end-to-end-distances_=0.70 on the end-to-end distances scores and have accordingly obtained cluster scores as shown in figure S18. We further note here that the assignment weights are not ad-hoc and are just merely based on the physical intuitions about the systems that we are trying to analyze. This would lead us to having 2 major states from the scoring metric-1 (folded and unfolded) and 3-4 major states from metric-2 for most algorithms which also implies some intermediate states as well apart from the two major folded and unfolded states.

In addition, we have used our DBSCAN based scoring metrics on the DBSCAN algorithm. We show the various clustering results in figure S19 and S20. Interestingly, the physically relevant properties of distances do not show any clear difference in density of their properties across the folding. So to cluster the peptide folding trajectory with DBSCAN, we chose the RMSD of the peptide as its representation. An optimal clustering was obtained at min-points=900 and ε=0.5 or min-points=150 and ε=0.1 as shown in figure S21. In both optimal clusters, we see from figures S19 and S20, that there is a clear separation in the RMSD as well as the end-to-end-distances. We again would comment that it is not wise to compare between the scores achieved by DBSCAN against the other clustering algorithms as their formulations are different. However, using our method we can simultaneously determine both min-points and ε in case of peptide-folding.

### 3.3 System protein-ligand

The final system consists of our unbiased simulations from a protein-ligand trajectory for an enzyme Smlt1473 bound to a substrate ManA^35^. The details of the docking and the simulations are provided in Reference 35^35^. In our trajectories, the sugar is mostly stable and remains in the enzyme binding pocket to change its various conformations.

To convert the raw trajectory into a usable ML data, the distances between the important highly interacting residues of the protein and the 6 units of the sugar are used to form a 2-dimensional array representation (86×6) of the pocket of the protein bound to the sugar with each column representing the bound state of the sugar to the enzyme for a single time step. In this matrix, the 1^st^ dimension (86 rows) denotes the important residues, and the 2^nd^ dimension denotes a particular sugar unit (6 columns). Once we have this matrix, we combine all the data across all the time frames to form a 7500×86×6 matrix to train a VAE to again create a reduced one-dimensional representation (200 dimensional) of the binding pocket for each time frame, thus forming a 7500×200-dimensional matrix. We next consider clustering these reduced 1-D representations of size 200 for a length 750 ns (7500-time steps).

The various properties of interest that we can consider for forming our scoring metrics would be the RMSD of the protein structure, the RMSD of the sugar structure bound to the protein, the distances between the various catalytic sites on the sugar to the protein, as well as the various hydrogen bonds and salt-bridges for the sugar interacting with the enzyme. The methods for calculations of these various properties are again given in Reference 32^35^. These properties for the WT Smlt1473 protein bound to the ManA sugar are plotted in figure 7. According to these various properties, we run our property-based scoring algorithms and present our various results. We represent the various cluster labels found out by our various clustering algorithms in figures S22,8,9,S23, and S24 and the respective ground truth labels in S25-S29 as well for the various chosen properties for scoring metric-1 like RMSD of the protein backbone, RMSD of the sugar, one-dimensional principal component representations of the distances between various residues on the protein to the sugar, the presence or absence of hydrogen-bonds and salt-bridges between the residues of the protein to the subunits of the sugar respectively obtained by using Principal-Component-Analysis^62^. In figure 10, the various scores for the algorithms are presented.

**Figure 7:**
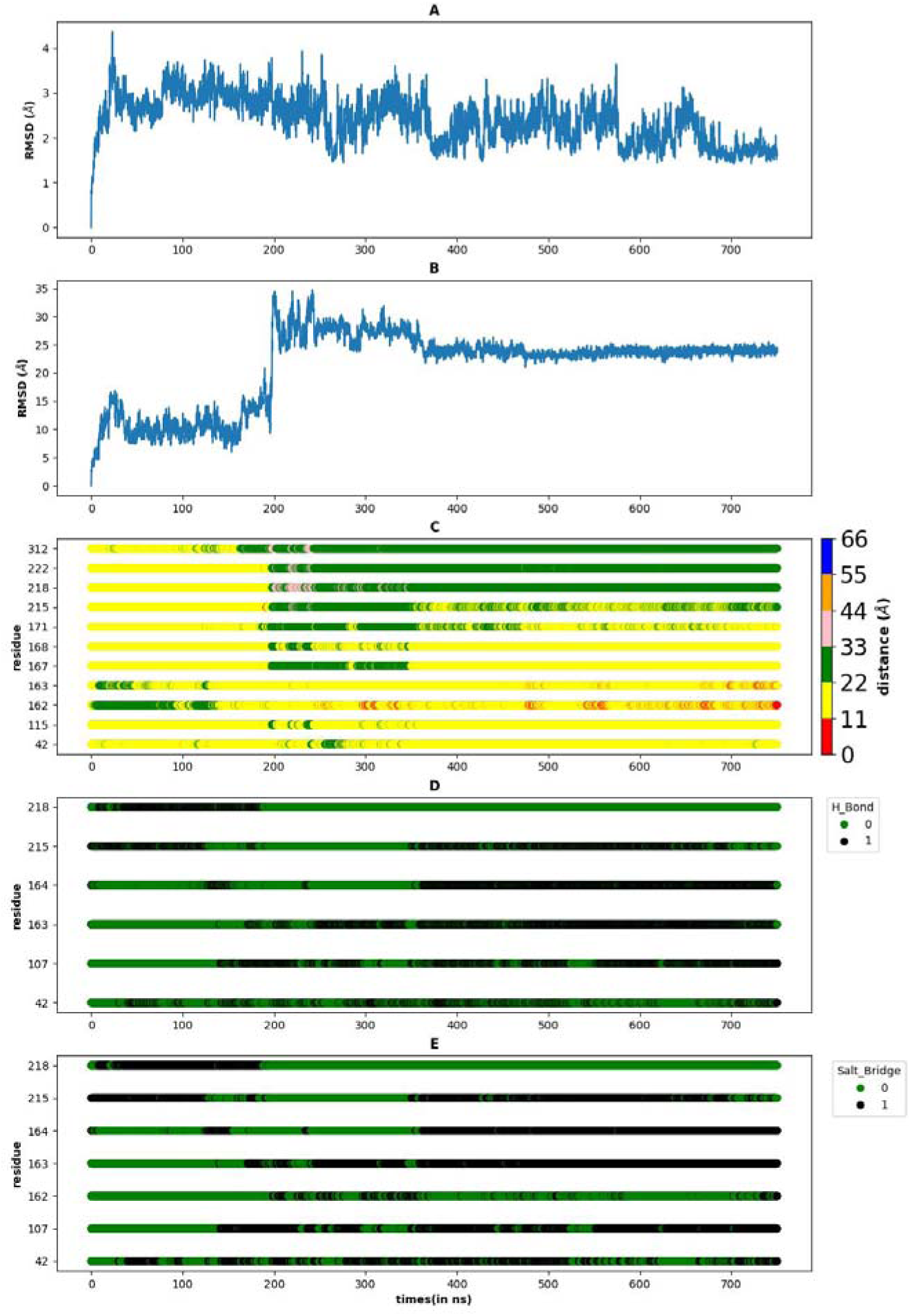
The various properties of the protein ligand simulation trajectories for Smlt1473 bound to Mana that we have. (A) denotes the RMSD of the protein plotted as a function of time. (B) denotes the RMSD of the ligand as a function of time. (C) denotes the distances between the various important residues of the protein to the sugar. The residues are shown on the y axis while the x axis denotes time in ns. (D) and (E) denotes the presence or absence of the hydrogen bond/salt bridges between the various residues and the sugar.

**Figure 8:**
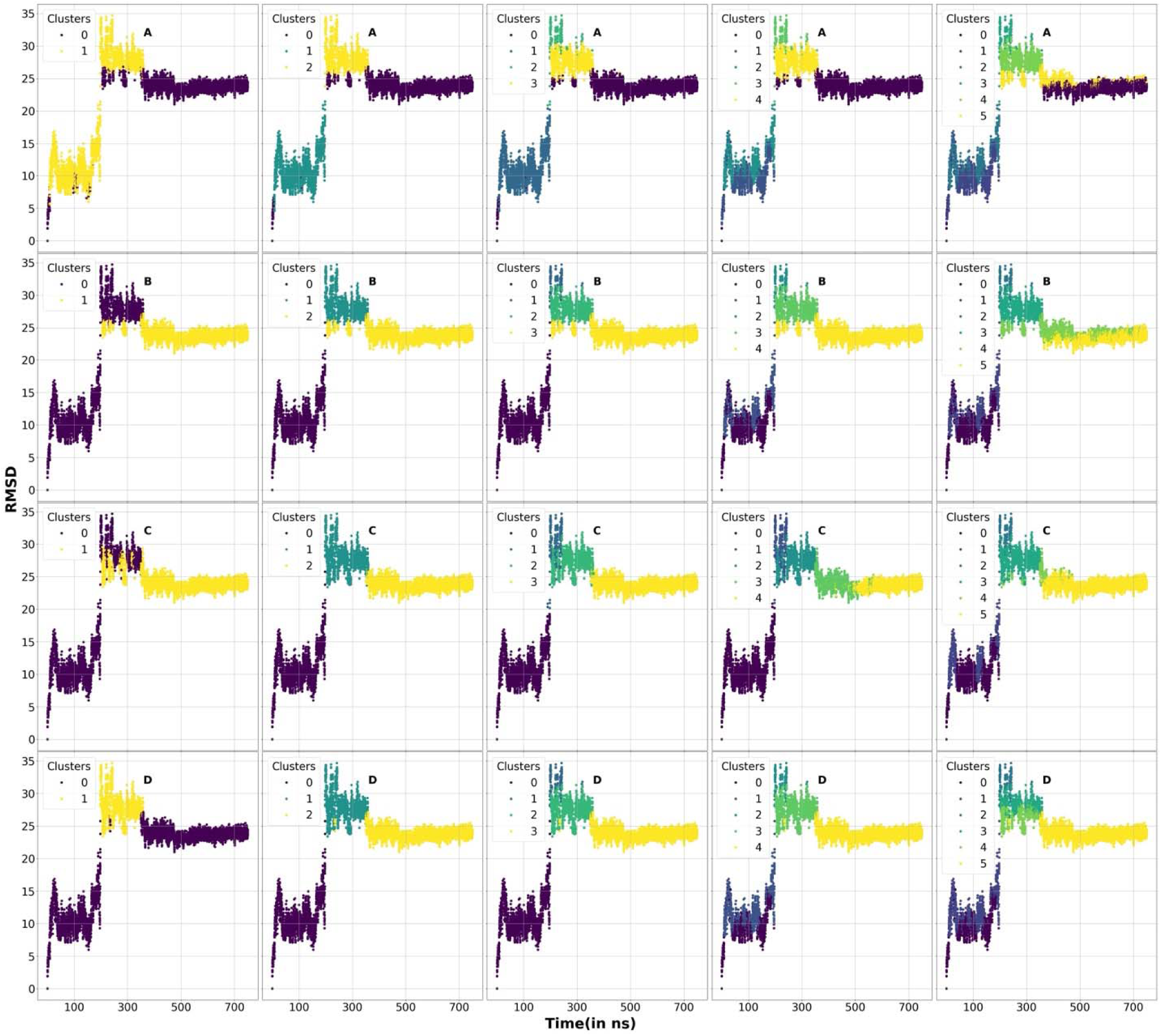
RMSD for the sugar properties of the protein-ligand simulation trajectories used for clustering for Smlt1473 bound to Mana. Different clusters are marked using different colors. A, B, C, and D sets denotes our 4 clustering algorithms K-Means, Agglomerative, BIRCH, and Agglomerative. Each column respectively denotes 2,3,4,5, and 6 clusters.

**Figure 9:**
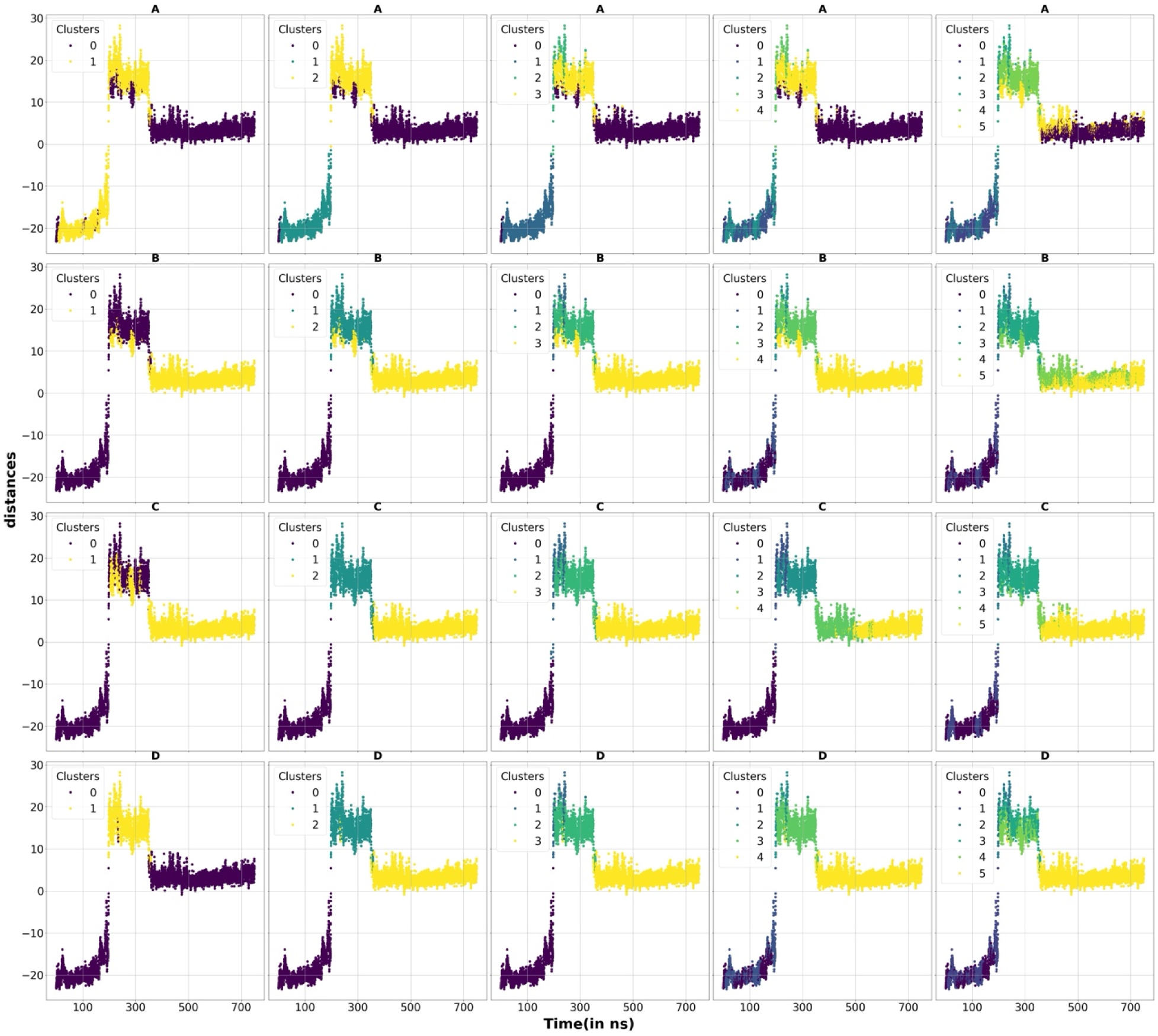
Reduced one-dimensional PCA component of protein-sugar distance properties of the protein-ligand simulation trajectories used for clustering for Smlt1473 bound to Mana. Different clusters are marked using different colors. A, B, C, and D sets denotes our 4 clustering algorithms K-Means, Agglomerative, BIRCH, and Agglomerative. Each column respectively denotes 2,3,4,5, and 6 clusters.

**Figure 10:**
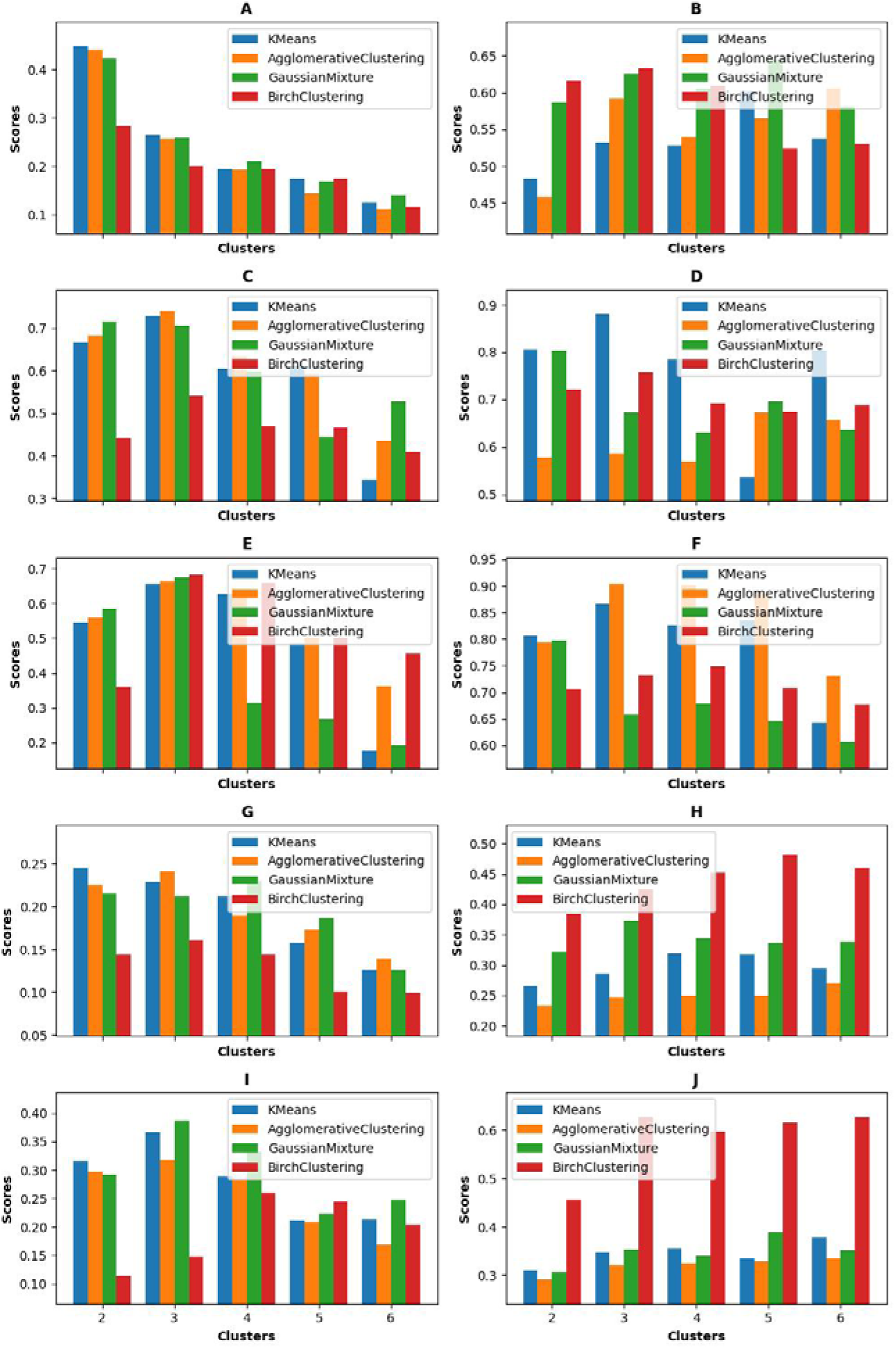
Scoring functions based on protein-ligand simulation trajectories used for clustering for Smlt1473 bound to Mana. (A) and (B) denotes the scoring functions-1 and 2 used on RMSD property for the backbone, (C) and (D) denotes the scoring functions-1 and 2 used on RMSD property of the sugar, (E) and (F) denotes the scoring functions-1 and 2 on the distances between the protein and the sugar, (G) and (H) denotes the hydrogen-bonds between the protein and the sugar while (I) and (J) denotes the salt-bridges between the protein and the sugar.

From the various scoring metrics, the RMSD of the protein backbone results in a total of 2 clusters for the scoring metric-1 again due to the best alignment whereas from the scoring metric-2 we get a total of 5 clusters for the backbone RMSD. For the ManA RMSD, the scoring function is able to identify a total of 3 clusters with both the scoring metrics which makes sense as 3 clusters align the best with both the ground truth labels and 3 clusters minimize the spread between the clusters. For the distances array, from the first scoring metric, we get an optimal of 3 clusters as our best scores as 3 clusters align the best with our ground truth labels for distances. From scoring metric-2, 3-5 clusters are optimal and this number of clusters also reduces the spread between the clusters. For hydrogen bonds and salt bridges, scoring metric-1 highlights 2-3 clusters as optimal as 2-3 clusters align the best with our actual cluster labels. From scoring metric-2, we get around 5 clusters for our optimal number of clusters for hydrogen bonds and 3 clusters for salt bridges as on looking carefully we see that 5 clusters and 3 clusters indeed does reduce the spread for the hydrogen bonds and salt bridges.

Our physical intuition with our protein-ligand system will be that the RMSD of the protein should not be an important property compared to RMSD of the sugar, the presence/absence of hydrogen bonds, or the distances between the catalytic residues to the sugar. In addition to this, the RMSD of the sugar and the distances of the sugar from the protein should have more importance in forming the scoring metrics than the presence or absence of hydrogen bonds/salt-bridges. So, we might decide to put more importance to these other properties other than the RMSD of the protein and thus form an overall scoring function F in equation 1 to determine the quality of our clustering. This would lead to 3-4 clusters from both scoring functions-1 and 2 as we have shown in figure S30 by taking a weight W_rmsd-sugar_=0.35, W_distances_=0.35, W_presence-of-h-bonds_=0.15, and W_presence-of-salt-bridges_=0.15. Again we would like to note that these weights are note ad-hoc but rather based on the physical intuitions about the system.

It is important to note the differences between the scores obtained through scoring functions-1 and 2. In this case, we see that unlike the cases for Ising model as well as protein folding/unfolding, we see here there are much less differences in the scores between the scoring functions-1 and 2. This is probably because in this case we used only one major stable state for clustering and thereafter scoring. This is why scoring function-1 was able to find a multitude of stable states that makes up this major stable state.

Finally, we also want to make the point that our algorithm is able to find the optimal clusters independent of the sampling achieved by MD. In our trajectory the ligand states are not sufficiently sampled from figure 7B. In spite of this, our clustering algorithms are able to find all the clusters in these poorly sampled spaces as well and our scoring metrics indeed points them out through the scores obtained as well. For example, all the three states sampled by the ligand in figure 7B are clustered optimally as shown in figure 8 and they are highlighted by our metrics. To further prove our point, we have also analyzed with our scoring functions a protein ligand dissociation trajectory which composes of two major states and our clustering algorithm only finds two major states like with the protein-folding/Ising model cases. Further details are provided in the SI.

Finally, we have analyzed our systems using DBSCAN algorithm and shown the various results for the various properties in figures S31 to S35. We show finally in figure S36 how our scoring metrics using DBSCAN is indeed able to find out the optimal min-points and ε combination for clustering using DBSCAN. The optimal min-points and ε value that we got with our scoring algorithms in figure S36A and S36B. Namely the optimal parameters obtained are min-points=40, ε =0.05 or min-points=80, ε =0.06 from scoring metric A or or min-points=80, ε =0.06 from scoring metric B. On observing the corresponding property plots especially, the ligand and the distances PCA property plots from figures S32 and S33 we see that these sets of parameter values indeed give us the best combination of clusters. Finally, we would comment again it would be unwise to compare the DBSCAN based scoring algorithms against the other scoring algorithms as they are designed differently.

## 4 Discussions and Conclusions

We have described two possible physically interpretable metrics that can quantify the quality of clustered data. All the clustering metrics available in literature are related to the separation of the clusters as well as the similarity of the members of each cluster based on the reduced one-dimensional order parameter space used for clustering itself. Since these metrics are based on the clustered dimension itself, they clearly lack any true physical interpretation. We have brought forth two metrics that take into consideration the microscopic and macroscopic properties of the system and ranks clustering algorithms accordingly. We find that the ranked clustering data also matches with our physical intuition about the systems.

For the two different metrics that we have developed, the thought processes behind these two metrics are a bit different. For our first metric (metric-1), we consider hypothetical ground truth labels and look at how well our original cluster labels align with these ground truth labels. The better these pseudo ground truth labels are aligned with respect to the original cluster labels the better score obtained from metric-1. For our second metric (metric-2), we look at how well is the sub-properties within each cluster are like each other by performing a pseudo clustering on these sub properties. For very distinct sub properties, metric-2 will give us back poor results. Finally, we also introduce an overall scoring function based on all important properties (equations 1-3).

We have considered 5 different clustering algorithms in our paper (i) K-Means (ii) Gaussian Mixture, (iii) Agglomerative, (iv) BIRCH, and (v) DBSCAN. The clustering algorithms that we have used are coupled with a dimensionality reduction algorithm, a VAE (Variational Auto-Encoder). To perform our clustering, we present our systems in a multi-dimensional matrix. To present our system in a form appropriate for 1-D clustering, we run our VAE on top of this and get a reduced 1D representation for our system. We then use this reduced 1D representation to perform our clustering based on different clustering algorithms and get our different clustering results.

The various systems considered are: (i) a simple well studied system called the Ising model, (ii) WT-HP35 protein folding and unfolding, (iii) protein-ligand unbiased simulated trajectory from our own previous study, and (iv) protein-ligand dissociated trajectory. We have presented our Ising model as a 30 x30-dimensional matrix, our simulated WT-HP35 protein as a 35×35 dimensional matrix and our protein ligand simulate trajectory with a 86×6 dimensional matrix. VAE on this multimdimensional matrix is run and create some 1D representations using our clustering algorithms.

For the Ising model, the four properties that we considered are the energy, magnetization, heat capacity, and susceptibility. Energy and magnetization are microscopic properties of each Ising system, and we used a normal algorithm to perform our clustering. Heat capacity and susceptibility are macroscopic properties, and we perform an average pooling operation on the microstate representations at each temperature to create our representation for the macrostate. After this we perform the clustering using different algorithms and using different number of clusters and finally score them using our two different scoring metrics. Both our scoring metrics clustered two distinct states for our paramagnetic and ferromagnetic lattices whereas the entire temperature scan forms a specific cluster apart from the transition region. This matches with our physical intuitions about the Ising model.

For the wildtype HP35 protein, the properties that we have considered are the RMSD of the protein and the distance between the first 5 and the last 5 residues of the protein. After this we perform the clustering using different algorithms and using different number of clusters and finally score them using our two different scoring metrics. Our scoring metric-1 clusters two distinct folded and unfolded states while our scoring metric-2 is able to identify the 4 different states which is likely able to define two intermediate states in between the folded and unfolded states.

For the wildtype protein-ligand simulations for our own Smlt1473 bound to the sugar, the various properties that we have considered are the RMSD of the protein, RMSD of the sugar, the protein-ligand contact distances, the H-bond, as well as the salt-bridges forming between the various residues of the protein. From these scores, we can obtain 3-4 different optimal clusters for the sugar binding to the protein as we had discussed in section 3.3. In addition to this, we have also analyzed a FKBP protein bound to DMSO. We show that our scoring algorithm is able to correctly identify the 2 states, bound and dissociated states from the trajectory.

Finally, we show how our clustering algorithms can be extended to clustering algorithms that do not take number of clusters as an input to the clustering algorithm, i.e., Density Based Clustering algorithms (DBSCAN). We show that using our scoring metrics we can select the optimal min-points and ε for any system. We further show that the results obtained match with the physical intuition about the systems.

It is also very important to highlight the differences between our scoring metrics-1 and 2. Our scoring metric-1 is very helpful in scoring the major stable states of the system as it uses ground truth labels to quantify the score. Whereas scoring metric-2 does not use ground truth labels but instead looks at the spread of the properties within each cluster, this can find major stable states as well as intermediates between these two major states. However, if all the states in the system are stable like our considered protein ligand system, without any metastable states/intermediates our there would be less difference between our scoring functions.

Finally, it is also important to compare our results against some recent studies on clustering quality and method development work. In a recent study, a kinetics-based clustering algorithm was implemented to segment the protein folding pathway into distinct states^56^. First, a proper Collective Variable is identified for the protein folding pathway. Then this collective variable undergoes downsampling and then a BEAST algorithm^63^ is applied to divide the CV trajectory into segments. Then each of these segments are standardized and normalized and finally these segments are projected onto 3-dimensional coordinates using PCA. Then finally using DBSCAN algorithm, these points are clustered into distinct states. Finally, to evaluate the clustering they used a Kuhn-Munkres Label Matching algorithm^64, 65^ to match the clusters with certain ground truth data which they obtained by using the Deep-TDA algorithm^66, 67^. In our case, we have developed a method which is more physically interpretable and does not rely on actual ground truth labels to build a clustering metric.

Lastly, our scoring metrics that we have developed are not only limited to MD simulation trajectories and generally applicable. For example, this metric can also be used to analyze the clustering quality of a collection of books for example based on different properties of the books like the number of pages, the number of words, the language on which it was written and so on. As shown here was well, these metrics not only would help determining the optimal number of clusters but also choose between different clustering algorithms.

## Acknowledgments

The high-performance computing cluster used for this study was Zaratan maintained by the Division of Information Technology at the University of Maryland. This work was in part supported by NSF (CHE2003912). The authors would like to thank D.E. Shaw Research for making the trajectories available which are reported in this paper.

## 5 Data and Code Availability

All codes associated with this work are uploaded to https://github.com/mondalkinjal/Clustering-Metrics

## References

1. Y. Wang, J. M. Lamim Ribeiro, and P. Tiwary, Current Opinion in Structural Biology 61 (2020) 139.

2. F. Noé et al., Annual review of physical chemistry 71 (2020) 361.

3. H. Sidky, W. Chen, and A. L. Ferguson, Molecular Physics 118 (2020) e1737742.

4. S. Kaptan, and I. Vattulainen, Advances in physics: X 7 (2022) 2006080.

5. J. Zhang et al., Journal of Chemical Theory and Computation 19 (2023) 4338.

6. C. Scherer et al., Journal of Chemical Theory and Computation 16 (2020) 3194.

7. S. John, and G. Csányi, The Journal of Physical Chemistry B 121 (2017) 10934.

8. A. Glielmo et al., Chemical Reviews 121 (2021) 9722.

9. L. Herron, et al., arXiv preprint arXiv:2308.14885 (2023)

10. Y. Wang, L. Herron, and P. Tiwary, Proceedings of the National Academy of Sciences 119 (2022) e2203656119.

11. Z. Belkacemi et al., The Journal of Chemical Physics 159 (2023)

12. S. Doerr, et al., arXiv preprint arXiv:1710.10629 (2017)

13. G. A. Tribello, and P. Gasparotto, Frontiers in molecular biosciences 6 (2019) 46.

14. F. Trozzi, X. Wang, and P. Tao, The Journal of Physical Chemistry B 125 (2021) 5022.

15. D. Suárez, and N. Díaz, Journal of chemical information and modeling 60 (2020) 5815.

16. A. Wolf, and K. N. Kirschner, Journal of molecular modeling 19 (2013) 539.

17. V. C. de Souza, L. Goliatt, and P. V. C. Goliatt, in 2017 IEEE Latin American Conference on Computational Intelligence (LA-CCI) (IEEE, 2017), pp. 1.

18. M. Ghorbani et al., The Journal of Chemical Physics 155 (2021)

19. S. Ishiai et al., Journal of chemical theory and computation 20 (2024) 819.

20. M.-K. Hsieh, and J. B. Klauda, The Journal of Physical Chemistry B 128 (2023) 150.

21. X. Liu et al., Journal of Chemical Theory and Computation 20 (2024) 665.

22. D. Wang, and P. Tiwary, The Journal of Chemical Physics 154 (2021)

23. D. Wang et al., Journal of Chemical Theory and Computation (2024)

24. L. Chen et al., Journal of Chemical Theory and Computation (2024)

25. S. a. T. V. a. W. S. Bray, Journal of Chemical Information and Modeling 62 (2022) 4591.

26. D. Ray, and M. Parrinello, Proceedings of the National Academy of Sciences 121 (2024) e2313542121.

27. V. Tänzel, M. Jäger, and S. Wolf, Journal of Chemical Theory and Computation (2024)

28. J. Shao et al., Journal of chemical theory and computation 3 (2007) 2312.

29. A. Rosenberg, and J. Hirschberg, in Proceedings of the 2007 joint conference on empirical methods in natural language processing and computational natural language learning (EMNLP-CoNLL)2007), pp. 410.

30. T. Caliński, and J. Harabasz, Communications in Statistics-theory and Methods 3 (1974)

31. P. J. Rousseeuw, Journal of computational and applied mathematics 20 (1987) 53.

32. D. L. Davies, and D. W. Bouldin, IEEE transactions on pattern analysis and machine intelligence (1979) 224.

33. B. M. McCoy, and T. T. Wu, The two-dimensional Ising model (Harvard University Press, 1973),

34. S. Piana, K. Lindorff-Larsen, and D. E. Shaw, Proceedings of the National Academy of Sciences 109 (2012) 17845.

35. K. Mondal, et al., bioRxiv (2024) 2024.09.24.614745.

36. M. Tschannen, O. Bachem, and M. Lucic, arXiv preprint arXiv:1812.05069 (2018)

37. D. P. Kingma, arXiv preprint arXiv:1312.6114 (2013)

38. K. Z. Takahashi, Physical Chemistry Chemical Physics 25 (2023) 658.

39. P. Ravindra, Z. Smith, and P. Tiwary, Molecular Systems Design & Engineering 5 (2020) 339.

40. Z. Smith et al., The Journal of Chemical Physics 149 (2018)

41. M. Ahmed, R. Seraj, and S. M. S. Islam, Electronics 9 (2020) 1295.

42. A. M. Ikotun et al., Information Sciences 622 (2023) 178.

43. K. P. Sinaga, and M.-S. Yang, IEEE access 8 (2020) 80716.

44. N. Liu et al., Information Sciences 557 (2021) 170.

45. N. Monath et al., in Proceedings of the 27th ACM SIGKDD Conference on knowledge discovery & data mining2021), pp. 1245.

46. E. K. Tokuda, C. H. Comin, and L. d. F. Costa, Physica A: Statistical mechanics and its applications 585 (2022) 126433.

47. A. D. Fontanini, and J. Abreu, in 2018 IEEE Power & energy society general meeting (PESGM) (IEEE, 2018), pp. 1.

48. T. Zhang, R. Ramakrishnan, and M. Livny, ACM sigmod record 25 (1996) 103.

49. T. Zhang, R. Ramakrishnan, and M. Livny, Data mining and knowledge discovery 1 (1997) 141.

50. G. Yu, G. Sapiro, and S. Mallat, IEEE Transactions on Image Processing 21 (2011) 2481.

51. M. Gogebakan, IEEE Access 9 (2021) 159987.

52. K. Khan et al., in The fifth international conference on the applications of digital information and web technologies (ICADIWT 2014) (IEEE, 2014), pp. 232.

53. O. Lemke, and B. G. Keller, The Journal of Chemical Physics 145 (2016)

54. S. Liu, et al., bioRxiv (2021) 2021.06.09.447666.

55. D. Deng, in 2020 7th international forum on electrical engineering and automation (IFEEA) (IEEE, 2020), pp. 949.

56. M. a. R. D. a. N. S. a. R. U. a. P. M. a. B. G. Faran, Journal of Chemical Theory and Computation 20 (2024) 5428.

57. M. Ester, et al., A density based algorithm for discovering clusters in large spatial databases with noise (1996) 226.

58. J. Sander et al., Data mining and knowledge discovery 2 (1998) 169.

59. S. Lee, et al., bioRxiv (2024)

60. T. K. Chiu et al., Proceedings of the National Academy of Sciences 102 (2005) 7517.

61. H. Klem, G. M. Hocky, and M. McCullagh, Journal of chemical theory and computation 18 (2022) 3218.

62. H. Abdi, and L. J. Williams, Wiley interdisciplinary reviews: computational statistics 2 (2010) 433.

63. K. Zhao et al., Remote Sensing of Environment 232 (2019) 111181.

64. Y. Chen et al., Information Sciences 433–434 (2018) 510.

65. H. W. Kuhn, Naval Research Logistics Quarterly 2 (1955) 83.

66. D. Ray, E. Trizio, and M. Parrinello, The Journal of Chemical Physics 158 (2023) 204102.

67. E. Trizio, and M. Parrinello, The Journal of Physical Chemistry Letters 12 (2021) 8621.

