## Supplementary Information for "Physically Interpretable Performance Metrics for Clustering"

**Scoring Metric-1 extended for DBSCAN algorithm**

Using DBSCAN algorithm we need to specify the min-points and ε to obtain our cluster labels. Using our algorithm, we can show how reliable our clustering is and thus choose the optimal min-points and ε.

Let’s say that using our clustering algorithm we have clustered all the datapoints that we have into certain cluster labels L. Since our scoring metric-1 relies on choosing proper ground truth labels for comparing our actual cluster labels to the ground truth labels, we first need to get proper pseudo ground truth labels for the various properties. To obtain the pseudo ground truth labels, we set some possible values of ε and min-points do a two-dimensional scan of the pseudo ground truth labels min-points and ε for each element p that belong to the important set of properties P_imp_. Then for all these scanned pseudo ground truth labels for the property p, we rank them using a score (SC) which is a product of the Silhouette score (sc) that we obtain by using only the clustered datapoints that belong to a specific cluster and fraction of the ground truth labels that belong to a specific cluster (f_clustered_). We note here that if there is only one type of cluster then we set the silhouette score (sc) to be 0. Then we choose the pseudo ground truth labels that has the highest score S. Finally, we use these pseudo ground truth labels as our ground truths for the property p (PL).

Next we iterate through each cluster j in the ground truth label and for each cluster j we gather the frame numbers in PL that are equal to j (F_j_). Then from our original cluster labels L, we find the subset $L_{f_{j}}$ $\subseteq$ L for all elements (frame numbers) f in F_j_. Then we find the most frequent number m from $L_{f_{j}}$ that maps to j from PL (ground truth labels). Then we define the two different fractions $f_{p_{mj}}$ which denotes the probability of occurrence of this number m in $L_{f_{j}},$ as well as probability of occurrence of F_j_ itself in the cluster labels L (i.e. excluding the unclustered points) which equals $f_{p_{j}}$. So based on this j to m mapping, we can define our alignment product score f_j_= $f_{p_{mj}}\times f_{p_{j}}$. Again if multiple j’s from the pseudo ground truth labels are mapped to the same m, then we just retain the maximum f_j_ and set all other f_j_’s to 0. Then we sum over f_j_’s to get our final score F. Thus, our final scoring function is SC$\times$F. Henceforth our final scoring function S_p_= SC$\times$F. We finally give a tabular description of our algorithm below.

**Algorithm-1 for DBSCAN**

1. Inputs: ε and min-points for the original cluster labels

2. Cluster Labels using algorithm to get cluster labels L = {L_1_, L_2,_ L_3_, …., L_n_} for N = {1, 2_,_ 3, …., n} trajectory frames

3. Calculate Property Values P = {P_1_, P_2,_ P_3_, …., P_n_}

4. Initialize ε-list and min-points-list for property P.

5. for ε_pseudo_ in ε-list

for min-points_pseudo_ in min-points-list

b. run DBSCAN on P with inputs ε_pseudo_ and min-points_pseudo_ to get pseudo ground truth labels PL= {PL_1_, PL_2,_ PL_3_, …., PL_n_}

c. Get the SC= Silhouette score (sc) obtained by using only the clustered datapoints that do not belong to a specific cluster* fraction of the ground truth labels that belong to a specific cluster (f_clustered_)

6. Get the ground truth labels PL with the maximal score SC

7. for cluster no j= 1, 2, 3, … k

a. Find F_j_$\subseteq$ N (frame numbers) for which PL = j

b. Find $L_{f_{j}}$ $\subseteq$ L for all elements f in F_j_

c. Find cluster element with most frequent occurrence in $L_{f_{j}}$ (m)

d. Calculate $f_{p_{mj}}$= probability of occurrence of m in $L_{f_{j}}$

e. Calculate $f_{p_{j}}$ = probability of occurrence of F_j_ in the cluster labels

f. f_j_= $f_{p_{mj}}\times f_{p_{j}}$

6. For each m and the corresponding j’s that are mapped to it, retain the maximum f_j_.

Set others to 0.

7. Final Score S_p_= SC$\times\sum_{j=1}^{k} f_{j}$

**Scoring Metric-2 extended for DBSCAN algorithm**

For our scoring metric-2, we again scan through min-points and ε to obtain our cluster labels. Here we show how using our scoring metric we can achieve the optimal min-points and ε. Let’s say that using our DBSCAN algorithm with inputs min-points and ε, we have clustered our data into cluster labels L. Now let’s say we have property values P for a property p which belongs to an important set of properties P_imp_. Then we iterate through each cluster j in L and find out frame numbers in L that belong to cluster j. Considering a subset of frames F_j_$\subseteq$ N (all the frames), we select properties $P_{f_{j}}$ from P, for each element/frame number f in F_j_. Once we have these sub-properties $P_{f_{j}}$ we do a clustering using for them to get the most dominant sub-property $P_{\mathrm{sub}_{f_{j}}}$. In order to do this, we perform a two-dimensional scan over several ε and min-points values to get our best choice for these parameters. In order to choose the best choice for ε and min-points, we again rank our ε and min-points values using the score SC that we had discussed in metric-1. So, we rank these ε and min-points values using the score SC again which is a product of the Silhouette score (sc) that is obtained by using only the clustered datapoints that fall to a specific cluster and the fraction of the datapoints within $P_{f_{j}}$that fall within a specific cluster (f_clustered_). We note that if there is only one type of cluster, we set the silhouette score (sc) to be 0. Then we choose the ε and min-points values that has the highest score SC. Finally based on these ε and min-points values as well as the subsequent clusters that we get when we try to subcluster $P_{f_{j}}$, we try to find out the most dominant subproperty $P_{\mathrm{sub}_{f_{j}}}$. If the score SC is less than a certain cutoff (sc-cutoff) we choose that further sub clustering is no longer possible and $P_{\mathrm{sub}_{f_{j}}}$ equals the whole set of sub-properties $P_{f_{j}}$.Else, we look at the majorly occurring subcluster label and we declare $P_{\mathrm{sub}_{f_{j}}}$as having the most majorly dominant sub property. Once we have done this we can again define the two fractions $f_{p_{j}}$which denotes the probability of occurrence of F_j_ in N, as well as $f_{p_{\mathrm{subj}}}$ that denotes the probability of occurrence of the most dominant subproperty $P_{\mathrm{sub}_{f_{j}}}$ within this subset F_j_. We can then take the product f_j_ =($f_{p_{j}}\times f_{p_{\mathrm{subj}}}$) to denote the weighted score of how majorly dominant each dominant subproperty $P_{\mathrm{sub}_{f_{j}}}$ is. After this, we can sum over the product f_j_ to get our final scoring term S_p_.

**Algorithm-2 for DBSCAN**

1. Inputs: min-points and ε. Score-cutoff (sc-cutoff)

2. Cluster Labels using DBSCAN to get cluster labels L = {L_1_, L_2,_ L_3_, …., L_n_} for N = {1, 2_,_ 3, …., n} trajectory frames

3. Calculate Property Values P = {P_1_, P_2,_ P_3_, …., P_n_}

4. Initialize ε-list and min-points-list for property P.

4. for cluster_no j= 1, 2, 3, … k

a. Find F_j_$\subseteq$ N (frame numbers) for which L = j

b. Find $P_{f_{j}}$ $\subseteq P (sub-properties) for$all elements f in F_j_

c. Sub-cluster $P_{f_{j}}$ using DBSCAN by performing a scan over ε-list and min-points-list.

d. Get Scores (SC) for ε_pseudo_ and min-points_pseudo_ in ε-list and min-points-list. SC=Silhouette score (sc) obtained by using only the clustered datapoints that do not belong to a specific cluster$\times$ fraction of the ground truth labels that belong to a specific cluster (f_clustered_) (sc=0 if there is only 1 cluster

e. Find the maximal Score (SC-max) and subsequent cluster labels.

f. If SC-max<sc-cutoff=0.30 (chosen here), no further sub-clustering possible: $P_{\mathrm{sub}_{f_{j}}}$=$P_{f_{j}}$	and goto h.

g. Look at optimal clusters through sc-max and the most dominant cluster= $P_{\mathrm{sub}_{f_{j}}}$

h. Calculate $f_{p_{j}}$= probability of occurrence of F_j_

i. Calculate $f_{p_{\mathrm{subj}}}$= probability of occurrence of $P_{\mathrm{sub}_{f_{j}}}$

j. f_j_ = $f_{p_{j}}\times$ $f_{p_{\mathrm{subj}}}$

5. Final Score S_p_= $\sum_{j=1}^{k} f_{j}$

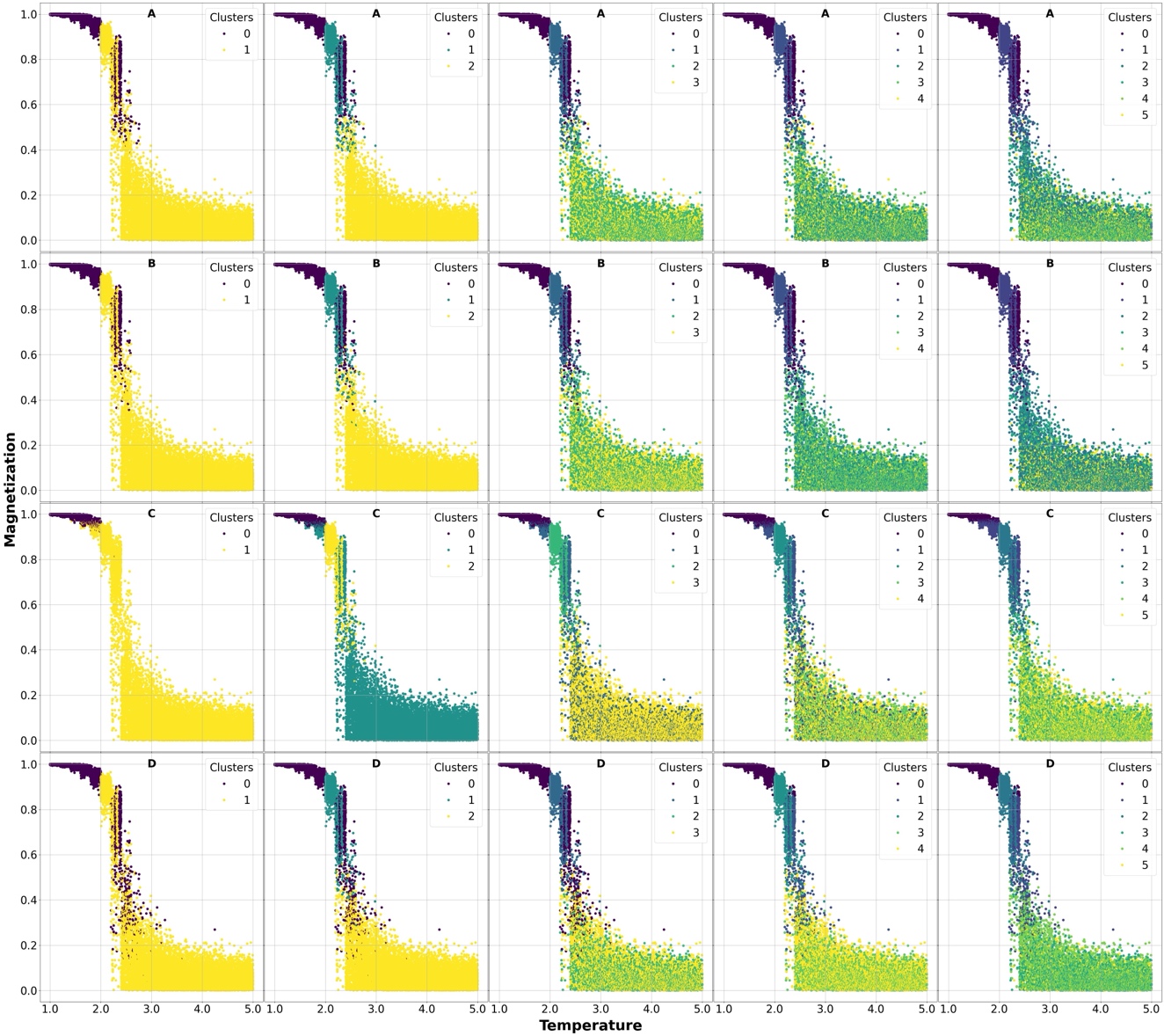

**Figure S1**: Magnetization properties of the different Ising model samples used for clustering using our first approach. Different clusters are marked using different colors. A, B, C, and D sets denotes our 4 clustering algorithms K-Means, Agglomerative, BIRCH, and Agglomerative. Each column respectively denotes 2,3,4,5, and 6 clusters.

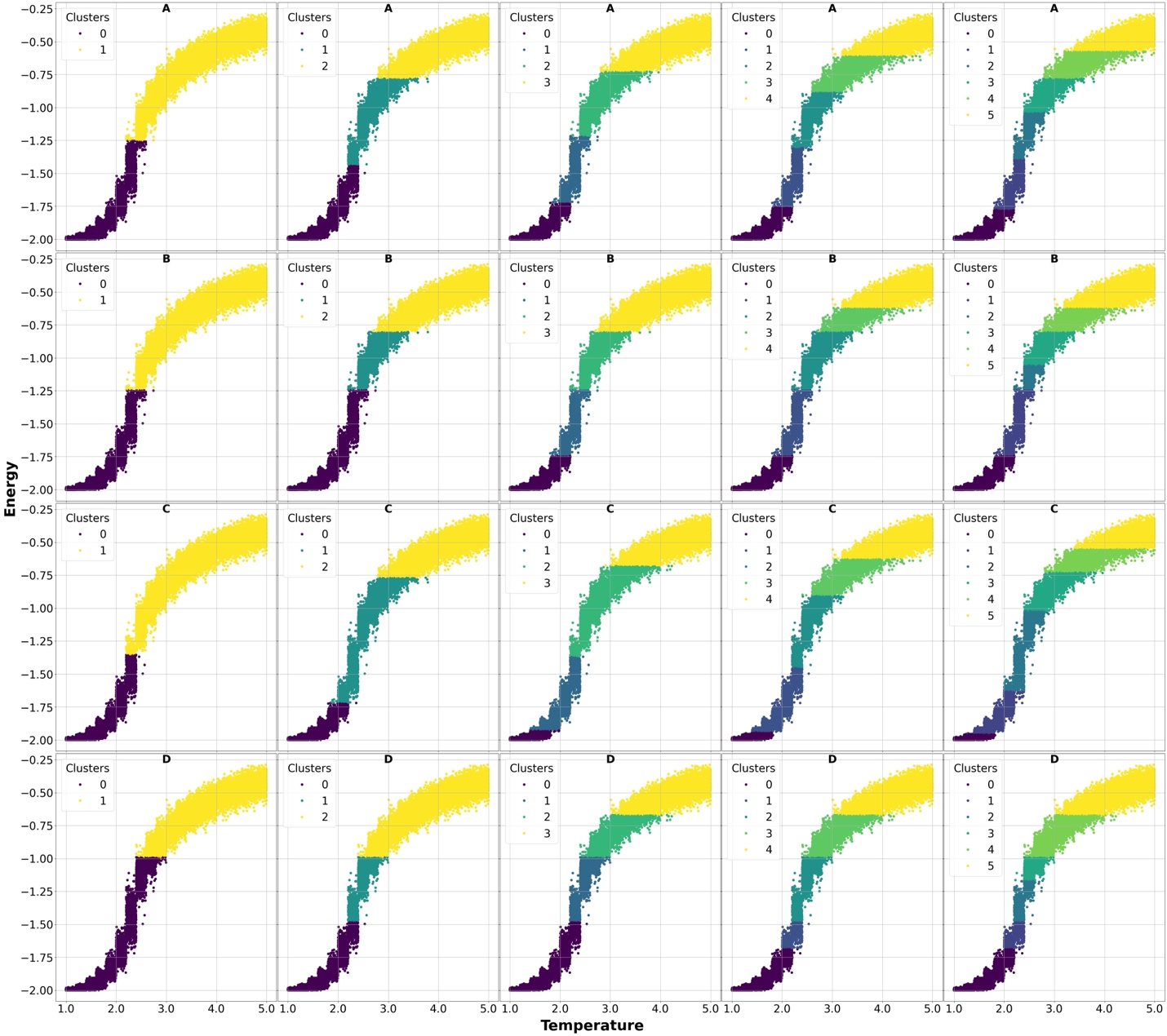

**Figure S2**: Actual Energy clusters found based on our ground truth label assignment (described in scoring metric-1). A, B, C, and D sets denotes our 4 clustering algorithms K-Means, Agglomerative, BIRCH, and Agglomerative. Each column respectively denotes 2,3,4,5, and 6 clusters

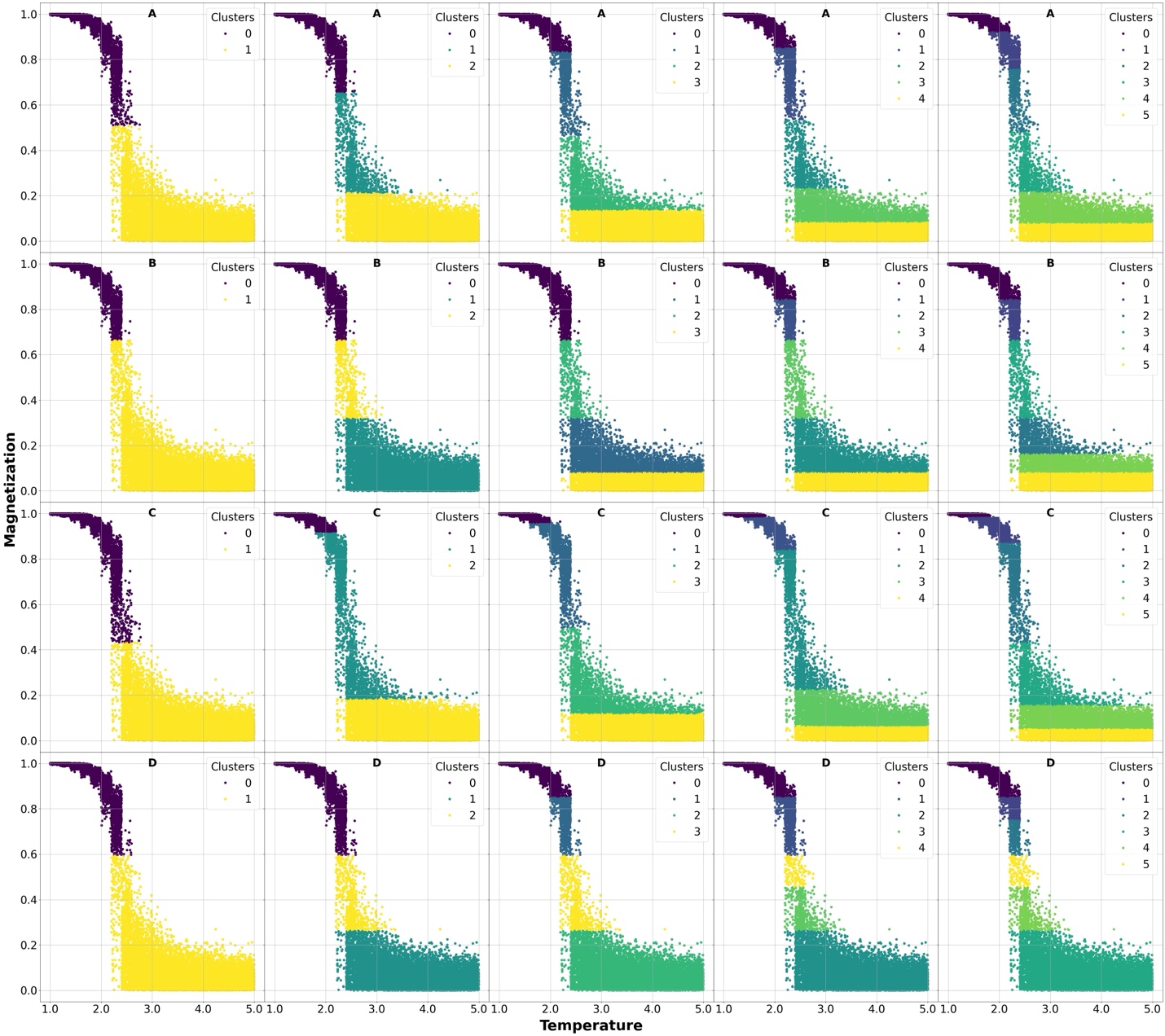

**Figure S3**: Actual Magnetization clusters found based on our ground truth label assignment (described in scoring metric-1). A, B, C, and D sets denotes our 4 clustering algorithms K-Means, Agglomerative, BIRCH, and Agglomerative. Each column respectively denotes 2,3,4,5, and 6 clusters.

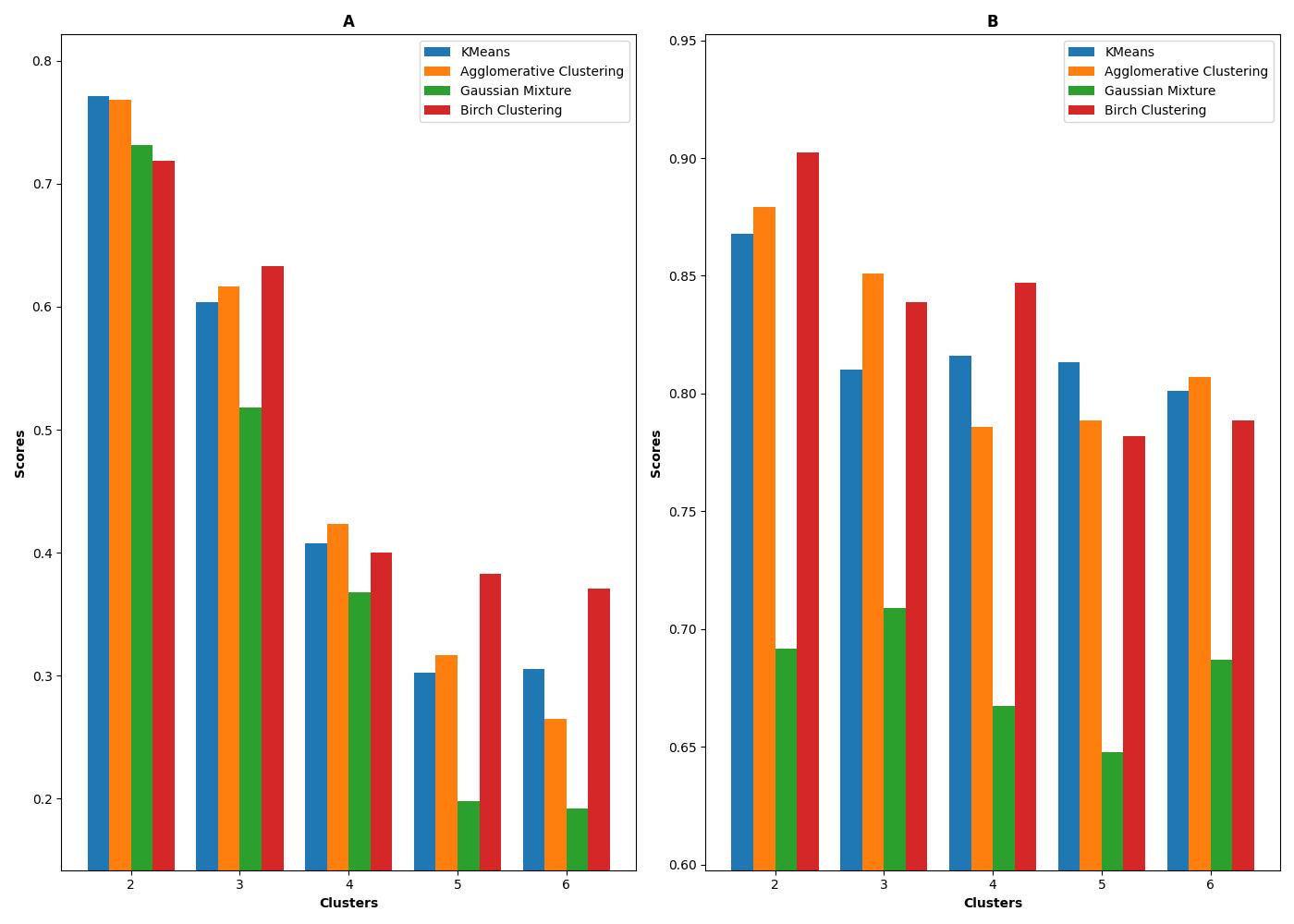

**Figure S4**: Overall scores based on our scoring function-1 (A) and scoring function-2 (B) based on the scores on energy and magnetization properties shown in figure 3. The weights Wenergy and Wmagnetization are both set to 0.5.

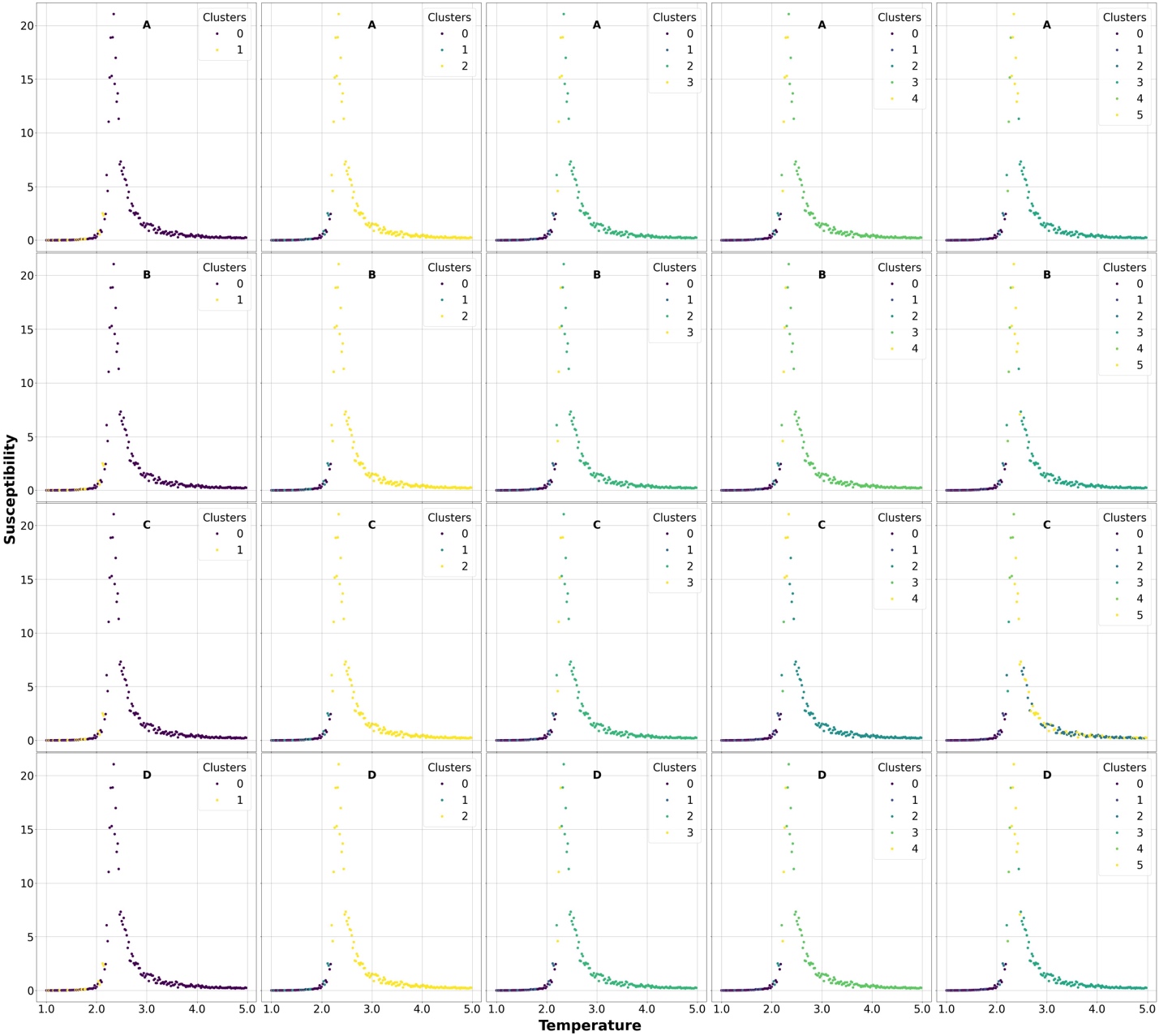

**Figure S5**: Susceptibility properties of the different Ising model samples used for clustering using our first approach. Different clusters are marked using different colours. A, B, C, and D sets denotes our 4 clustering algorithms K-Means, Agglomerative, BIRCH, and Agglomerative. Each column respectively denotes 2,3,4,5, and 6 clusters.

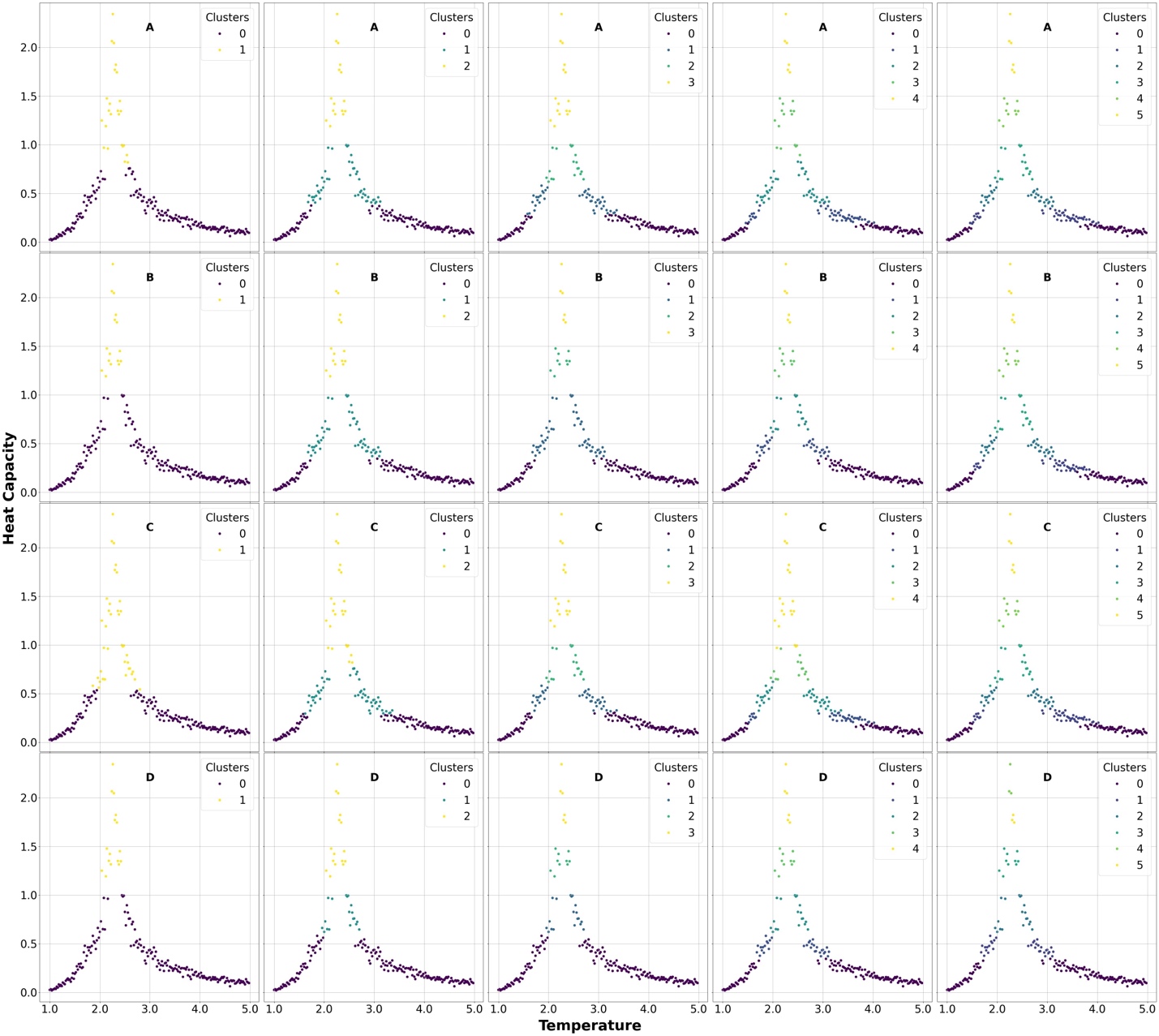

**Figure S6**: Actual Heat Capacity clusters found based on our ground truth label assignment (described in scoring metric-1). A, B, C, and D sets denotes our 4 clustering algorithms K-Means, Agglomerative, BIRCH, and Agglomerative. Each column respectively denotes 2,3,4,5, and 6 clusters.

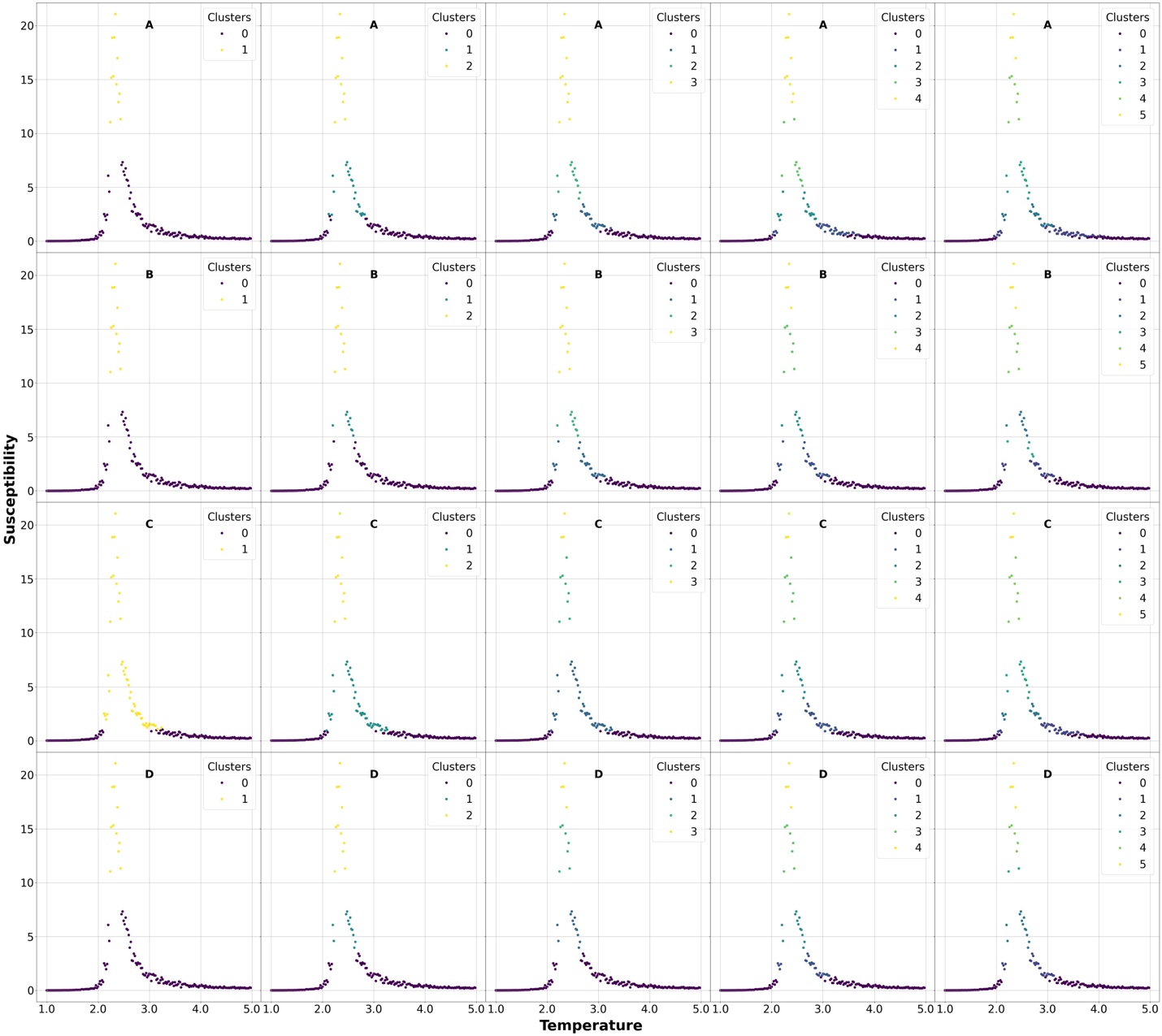

**Figure S7**: Actual Susceptibility clusters found based on our ground truth label assignment (described in scoring metric-1). A, B, C, and D sets denotes our 4 clustering algorithms K-Means, Agglomerative, BIRCH, and Agglomerative. Each column respectively denotes 2,3,4,5, and 6 clusters.

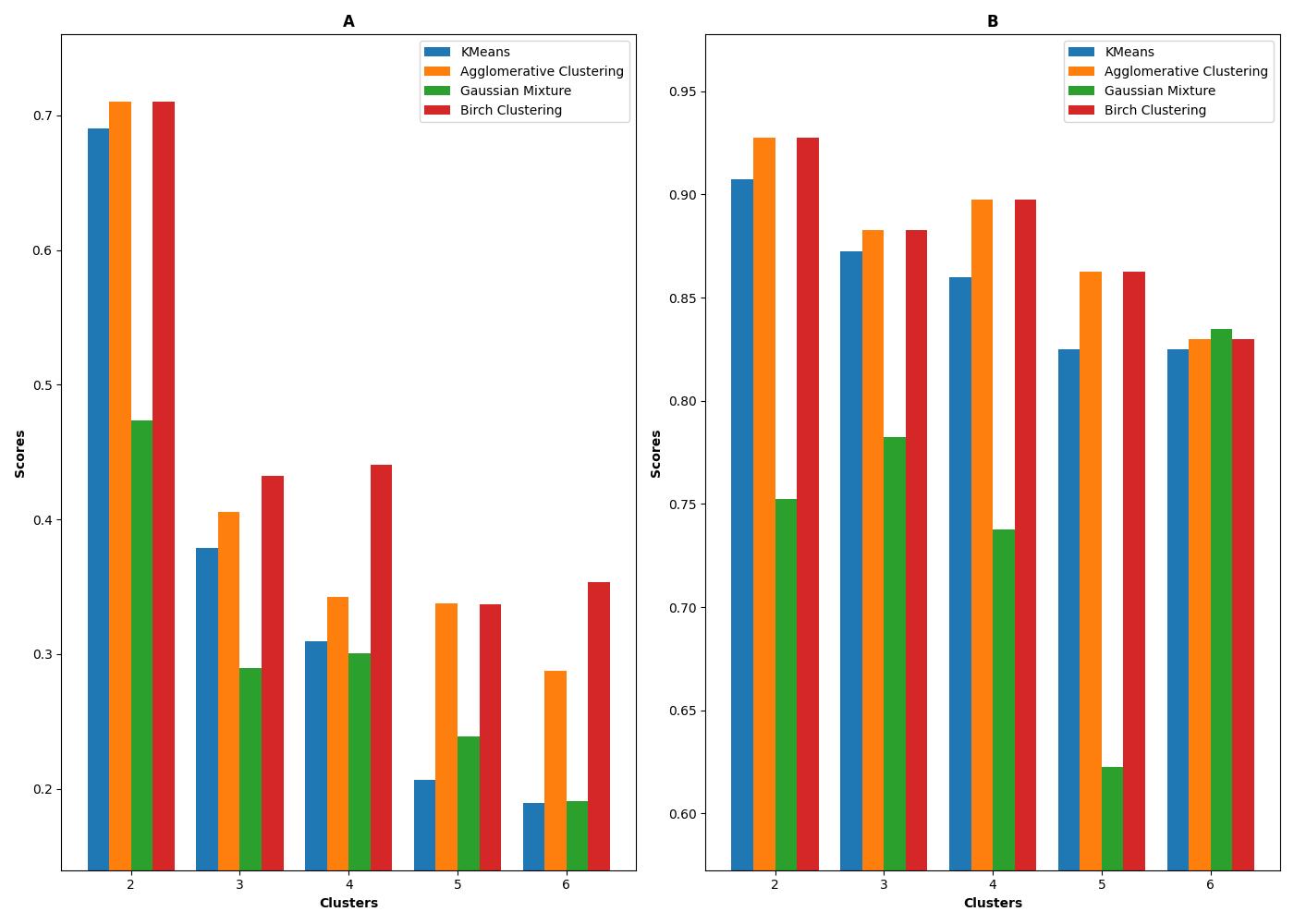

**Figure S8**: Overall scores based on our scoring function-1 (A) and scoring function-2 (B) based on the scores on heat capacity and susceptibility properties shown in figure 5. The weights Wheat-capacity and Wsusceptibility are both set to 0.5.

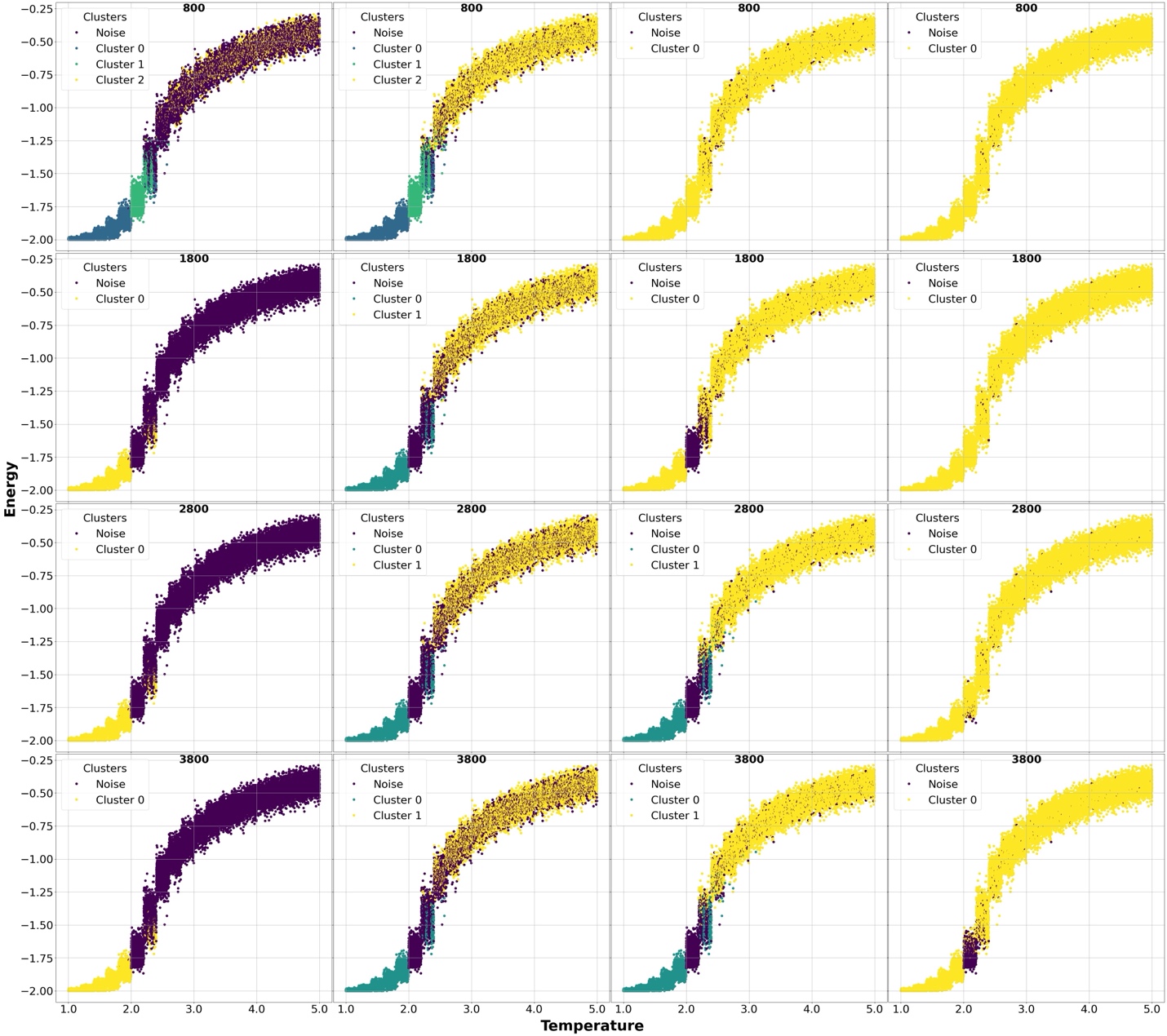

**Figure S9:** Energy properties of the different Ising model samples used for clustering by DBSCAN algorithm using our first approach. Different clusters as well as the unclustered points(noise) are marked using different colors. The min-points used for the clustering are shown in labels. Each row represents the min-points used in labels with increasing ε values=0.4, 0.5,0.6, and 0.7.

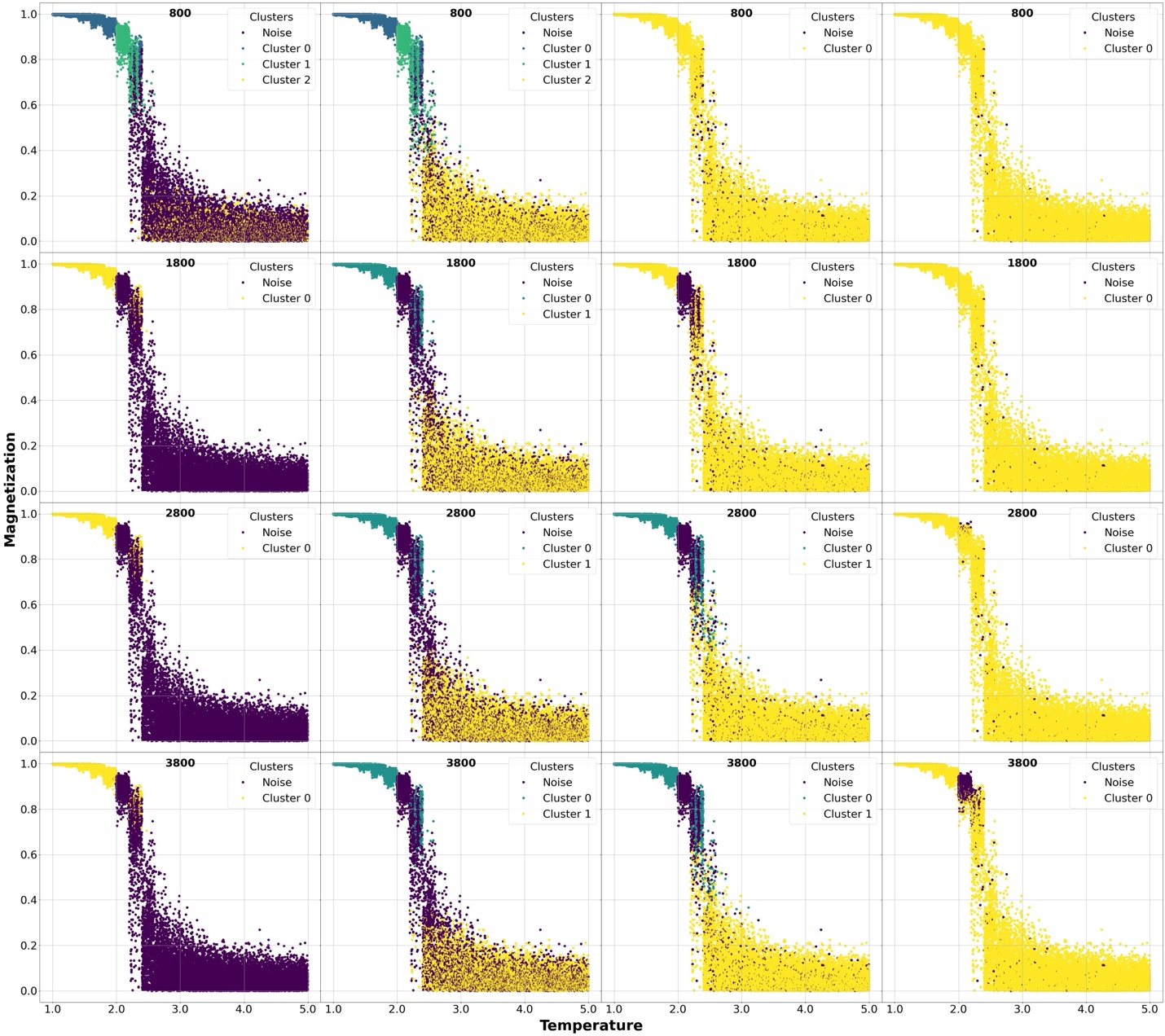

**Figure S10:** Magnetization properties of the different Ising model samples used for clustering by DBSCAN algorithm using our first approach. Different clusters as well as the unclustered points(noise) are marked using different colors. The min-points used for the clustering are shown in labels. Each row represents the min-points used in labels with increasing ε values=0.4, 0.5,0.6, and 0.7.

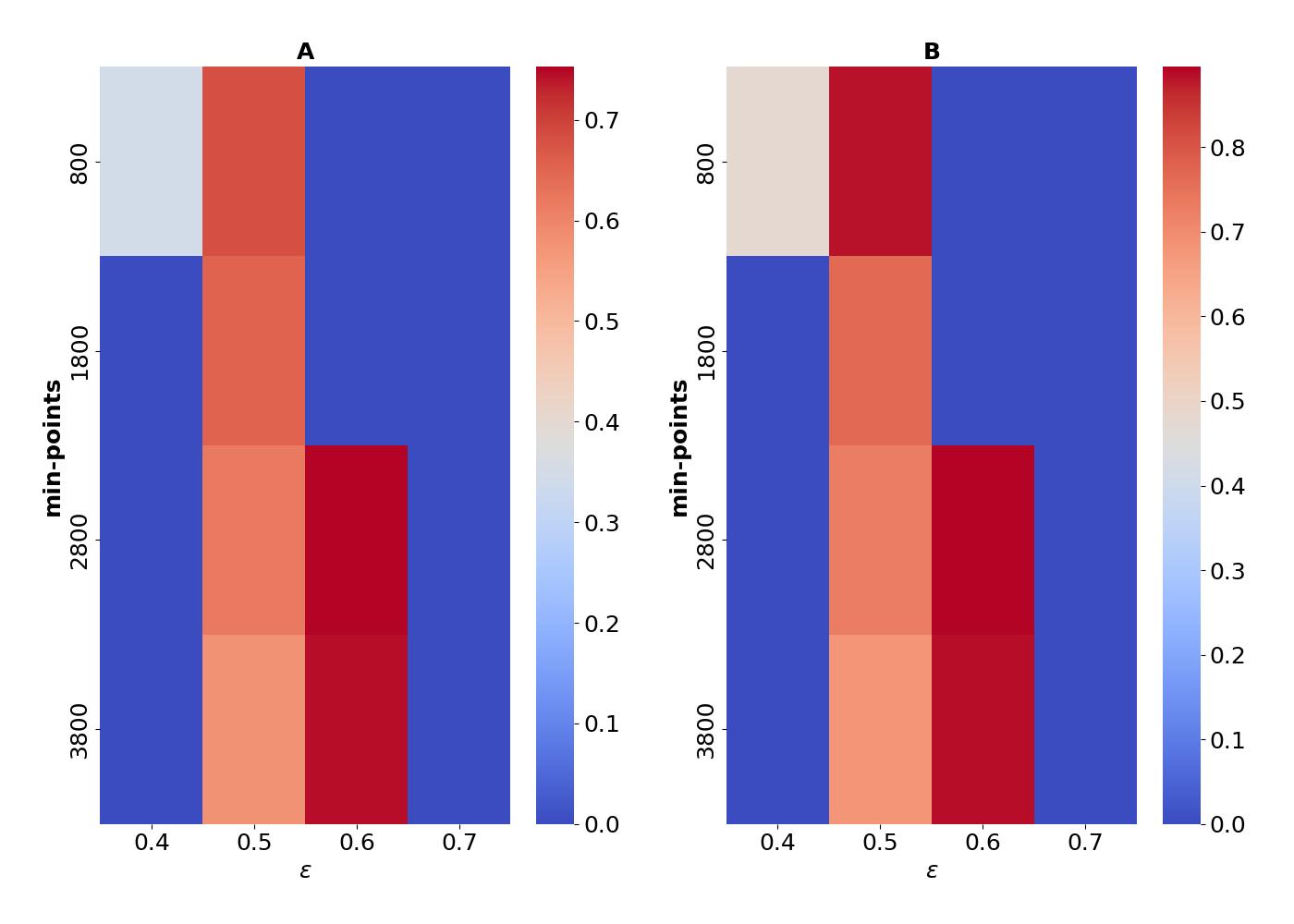

**Figure S11:** A two dimensional heatmap showing the scores obtained by our scoring metric-1(A) and scoring metric-2(B) for DBSCAN algorithm obtained by setting equal weights to energy(0.5) and magnetization (0.5) to our overall scoring function(Equation-1 in the main manuscript).

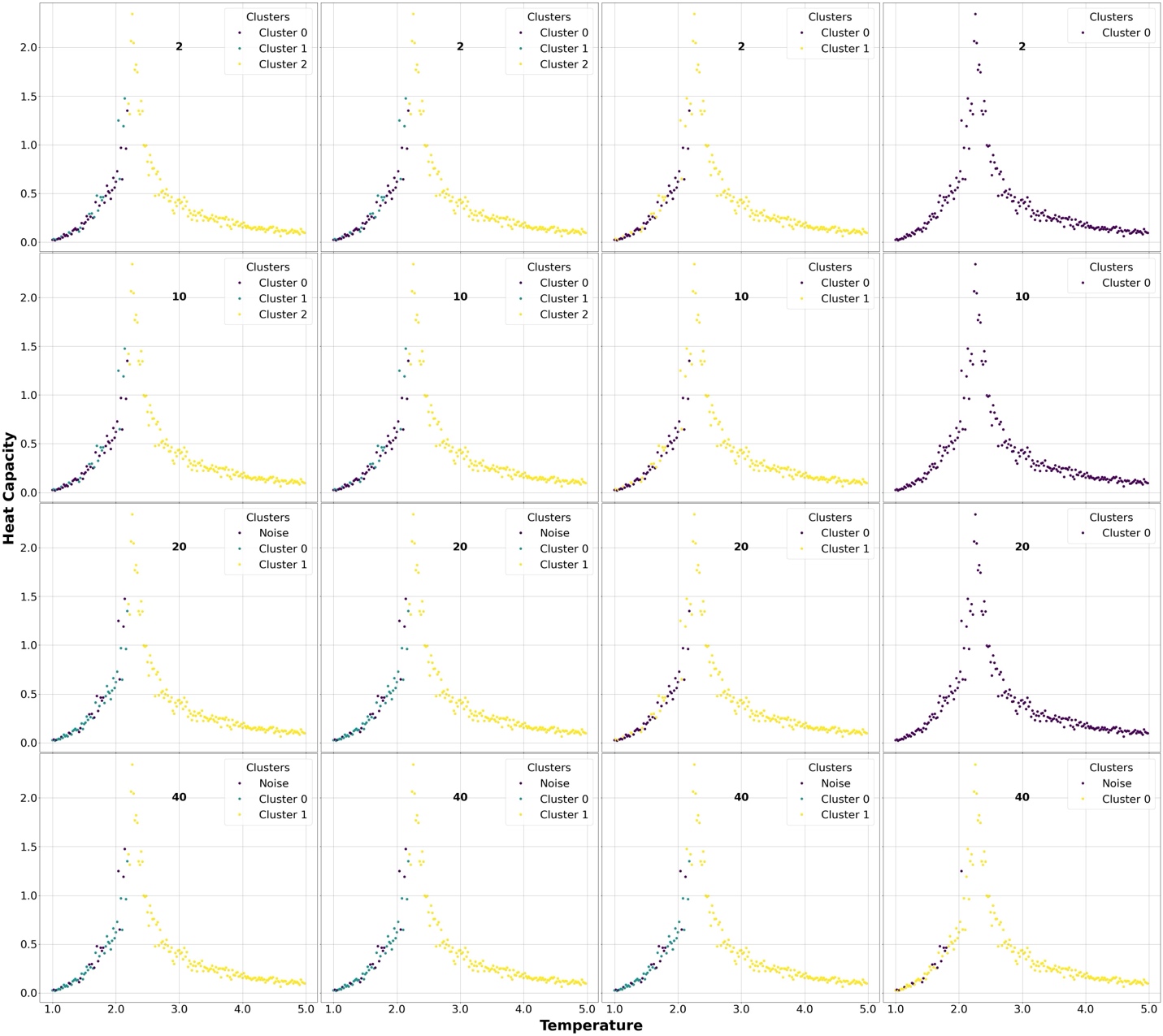

**Figure S12:** Heat Capacity properties of the different Ising model samples used for clustering by DBSCAN algorithm using our second approach. Different clusters as well as the unclustered points(noise) are marked using different colors. The min-points used for the clustering are shown in labels. Each row represents the min-points used in labels with increasing ε values=0.4, 0.5,0.6, and 0.7

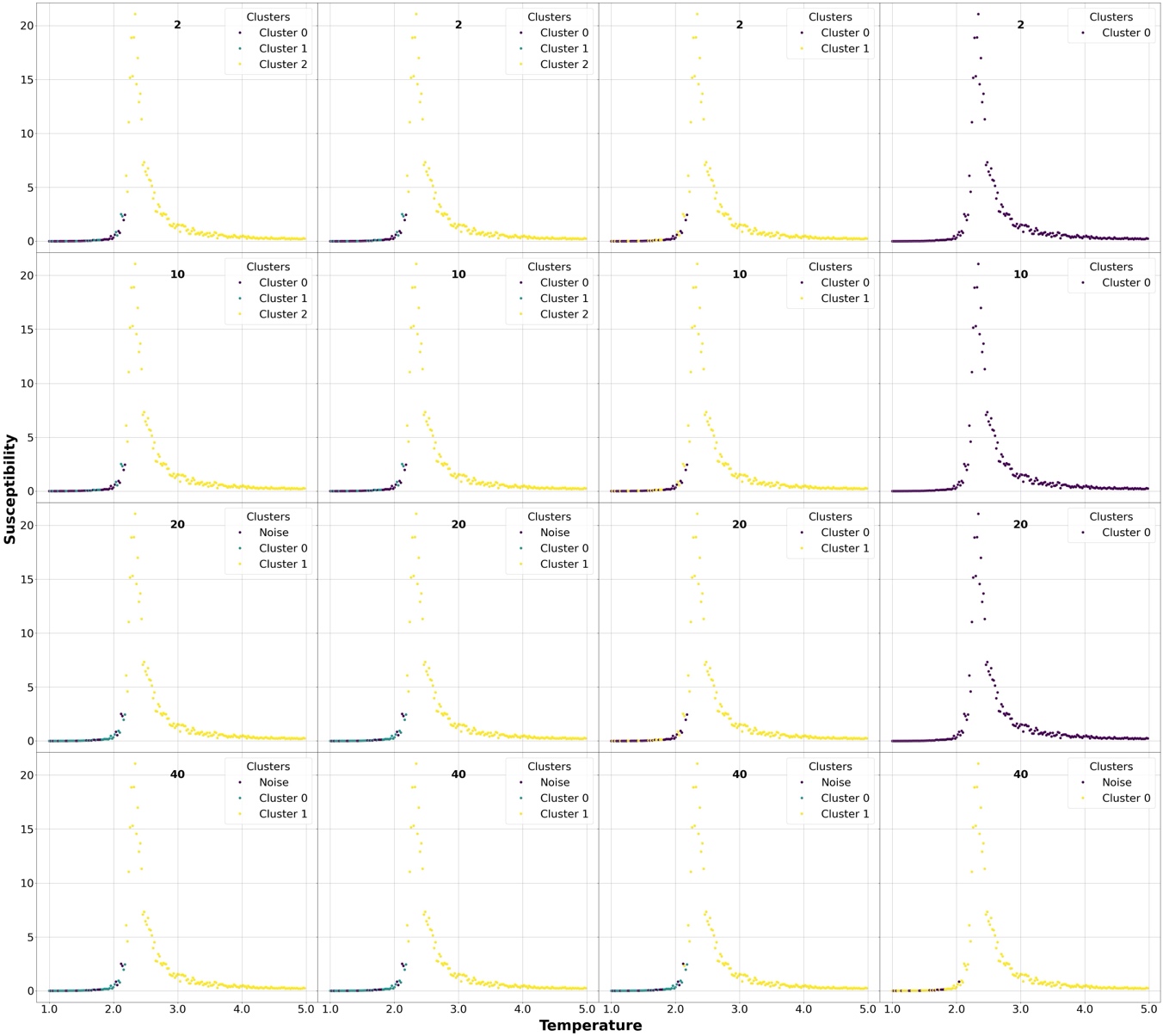

**Figure S13:** Susceptibility properties of the different Ising model samples used for clustering by DBSCAN algorithm using our second approach. Different clusters as well as the unclustered points(noise) are marked using different colors. The min-points used for the clustering are shown in labels. Each row represents the min-points used in labels with increasing ε values=0.4, 0.45,0.5, and 0.55.

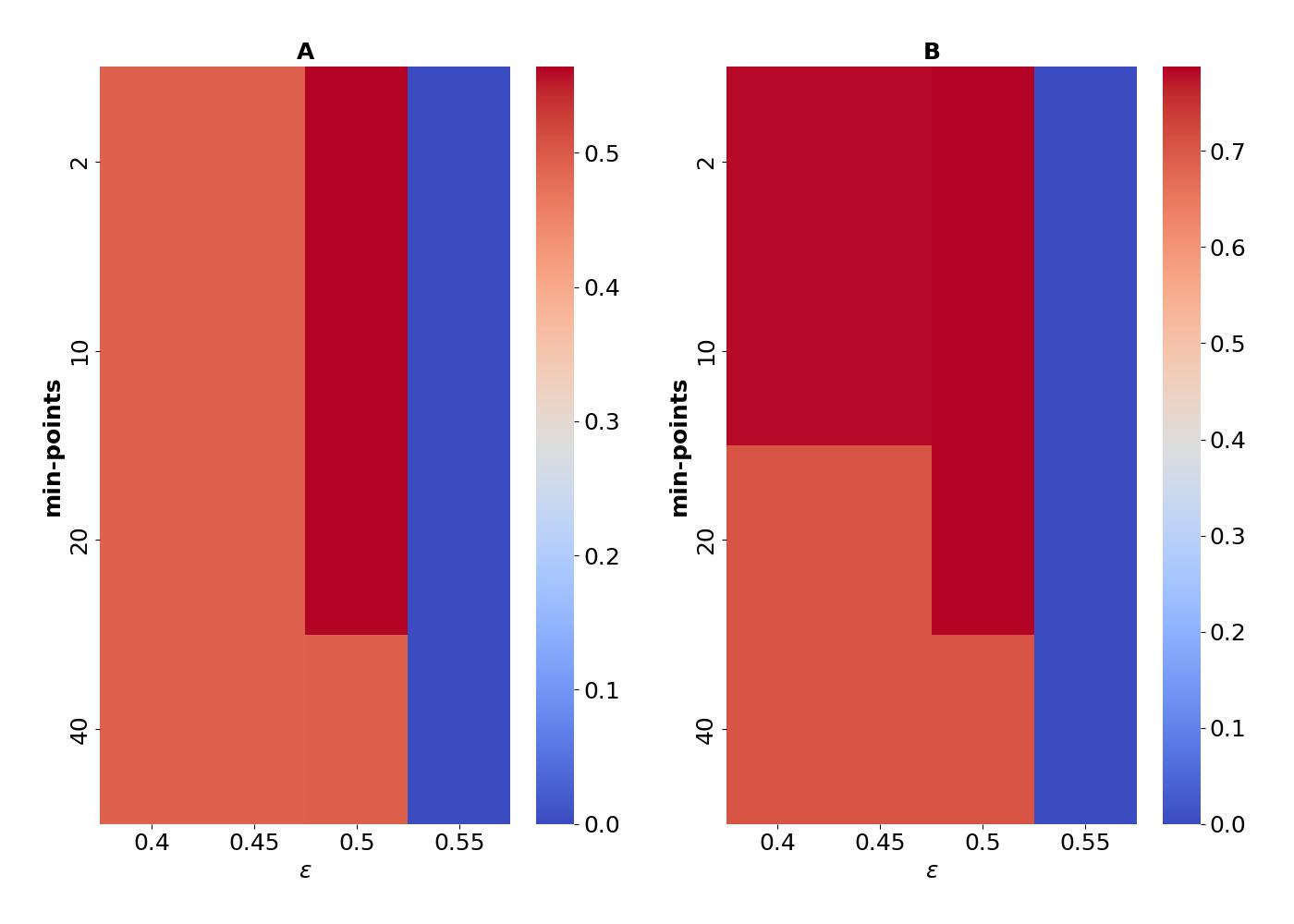

**Figure S14:** A two dimensional heatmap showing the scores obtained by our scoring metric-1(A) and scoring metric-2(B) for DBSCAN algorithm obtained by setting equal weights to heat capacity(0.5) and susceptibility (0.5) to our overall scoring function(Equation-1 in the main manuscript).

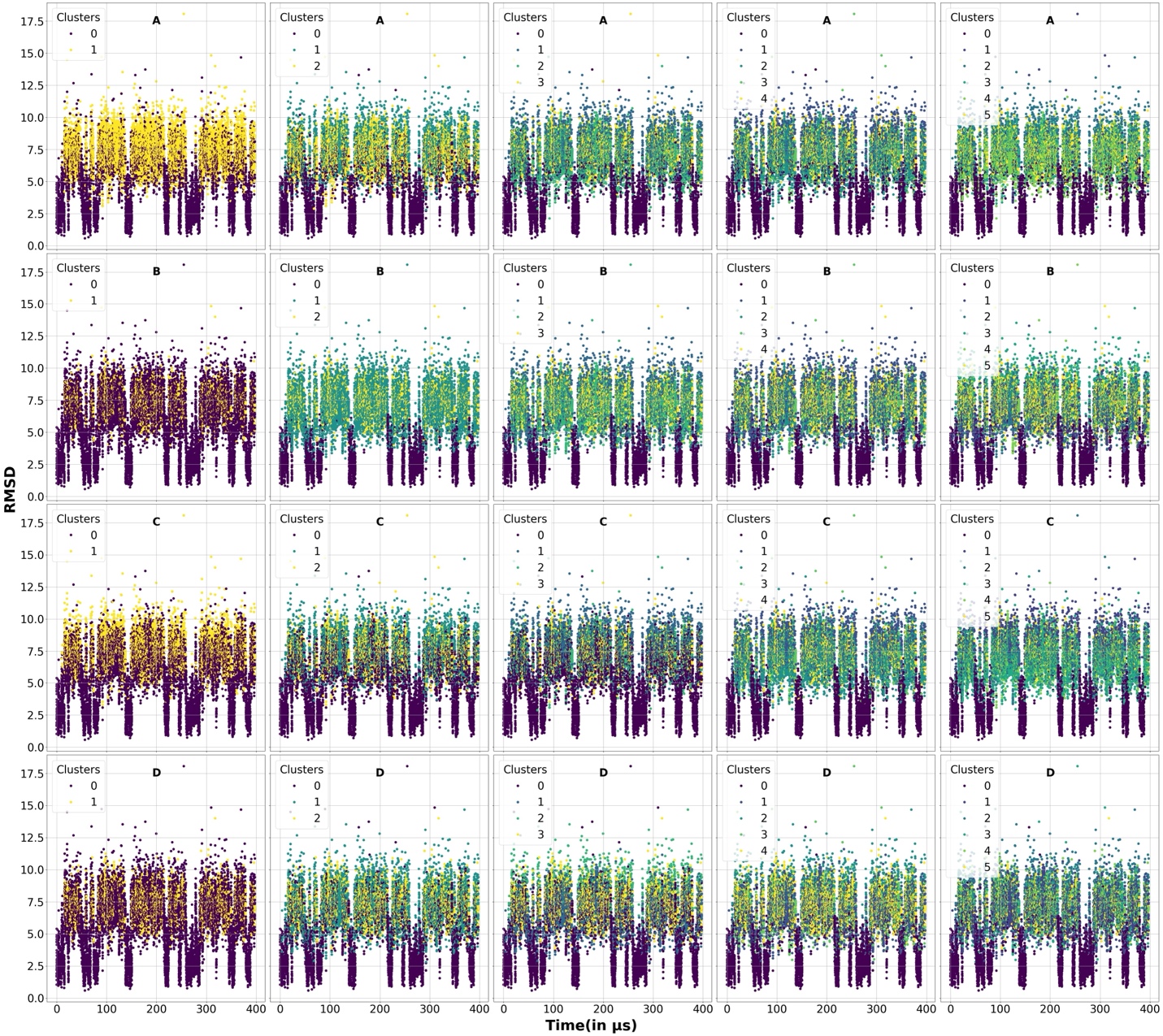

**Figure S15**: RMSD properties of the protein folding simulation trajectories used for clustering. Different clusters are marked using different colors. A, B, C, and D sets denote our 4 clustering algorithms K-Means, Agglomerative, BIRCH, and Agglomerative. Each column respectively denotes 2,3,4,5, and 6 clusters.

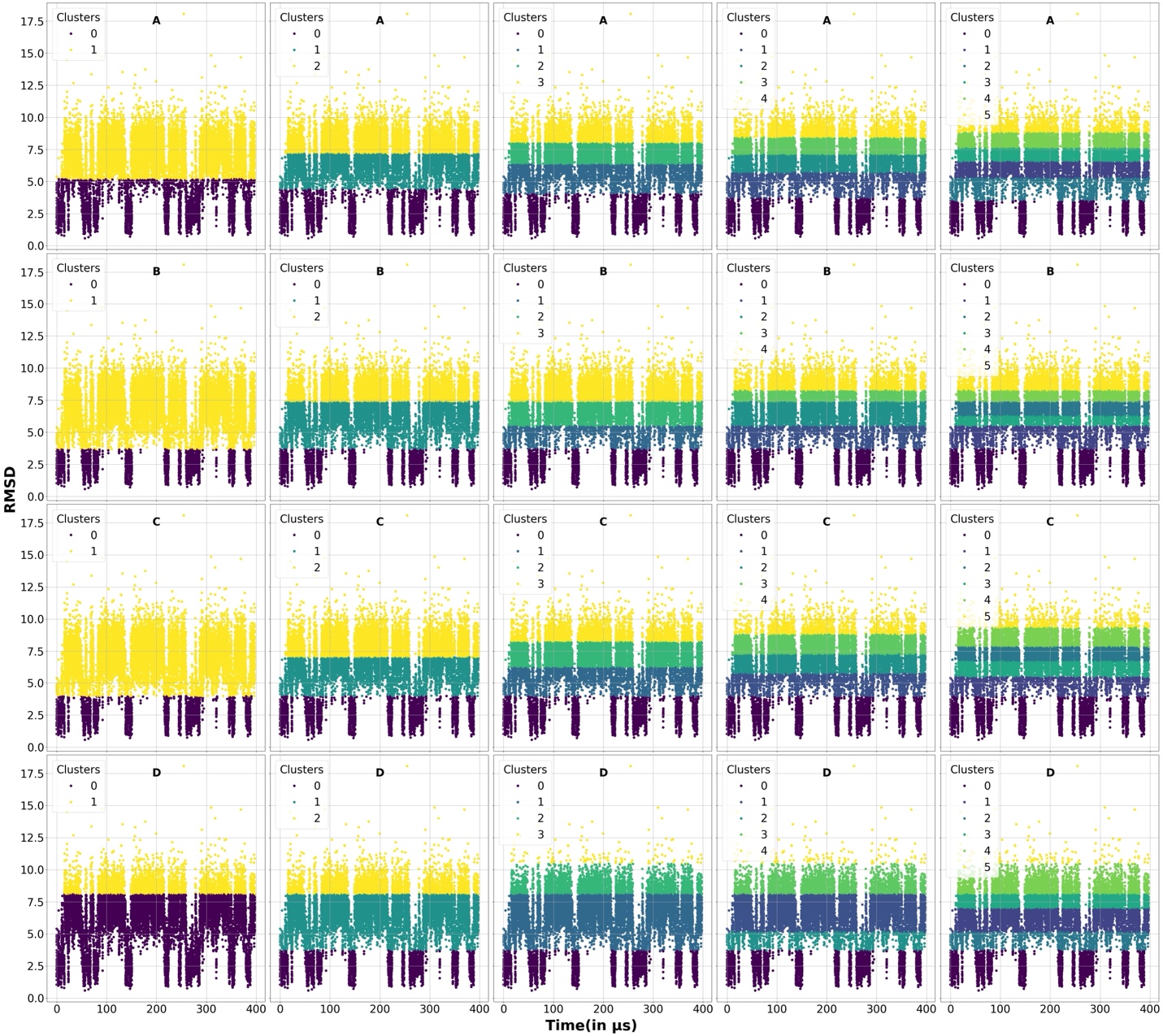

**Figure S16**: Actual RMSD for the protein clusters found based on our ground truth label assignment for metric-1. A, B, C, and D sets denotes our 4 clustering algorithms K-Means, Agglomerative, BIRCH, and Agglomerative. Each column respectively denotes 2,3,4,5, and 6 clusters.

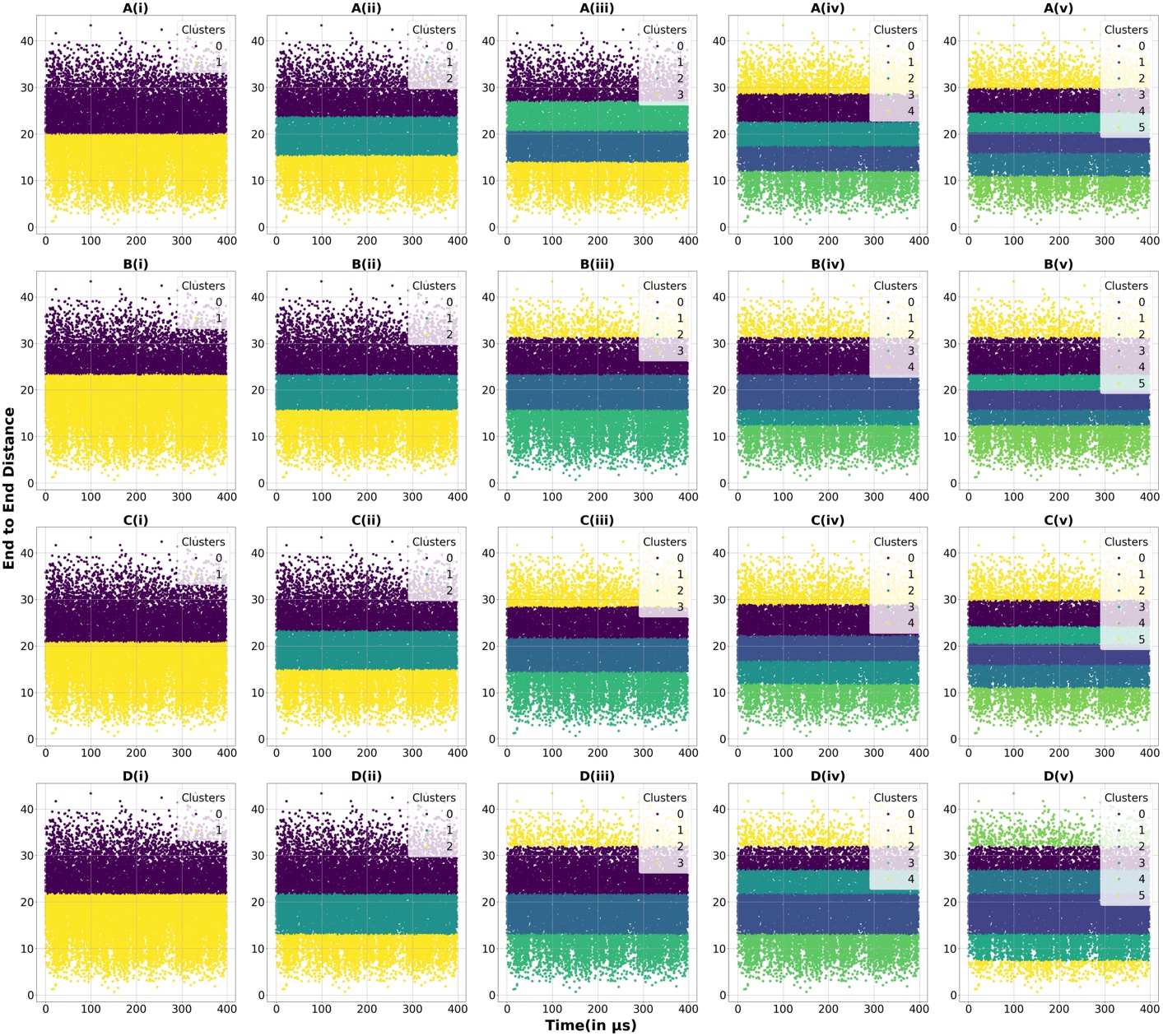

**Figure S17**: Actual end-to-end distances clusters found based on our ground truth label assignment. A, B, C, and D sets denotes our 4 clustering algorithms K-Means, Agglomerative, BIRCH, and Agglomerative. Each column respectively denotes 2,3,4,5, and 6 clusters.

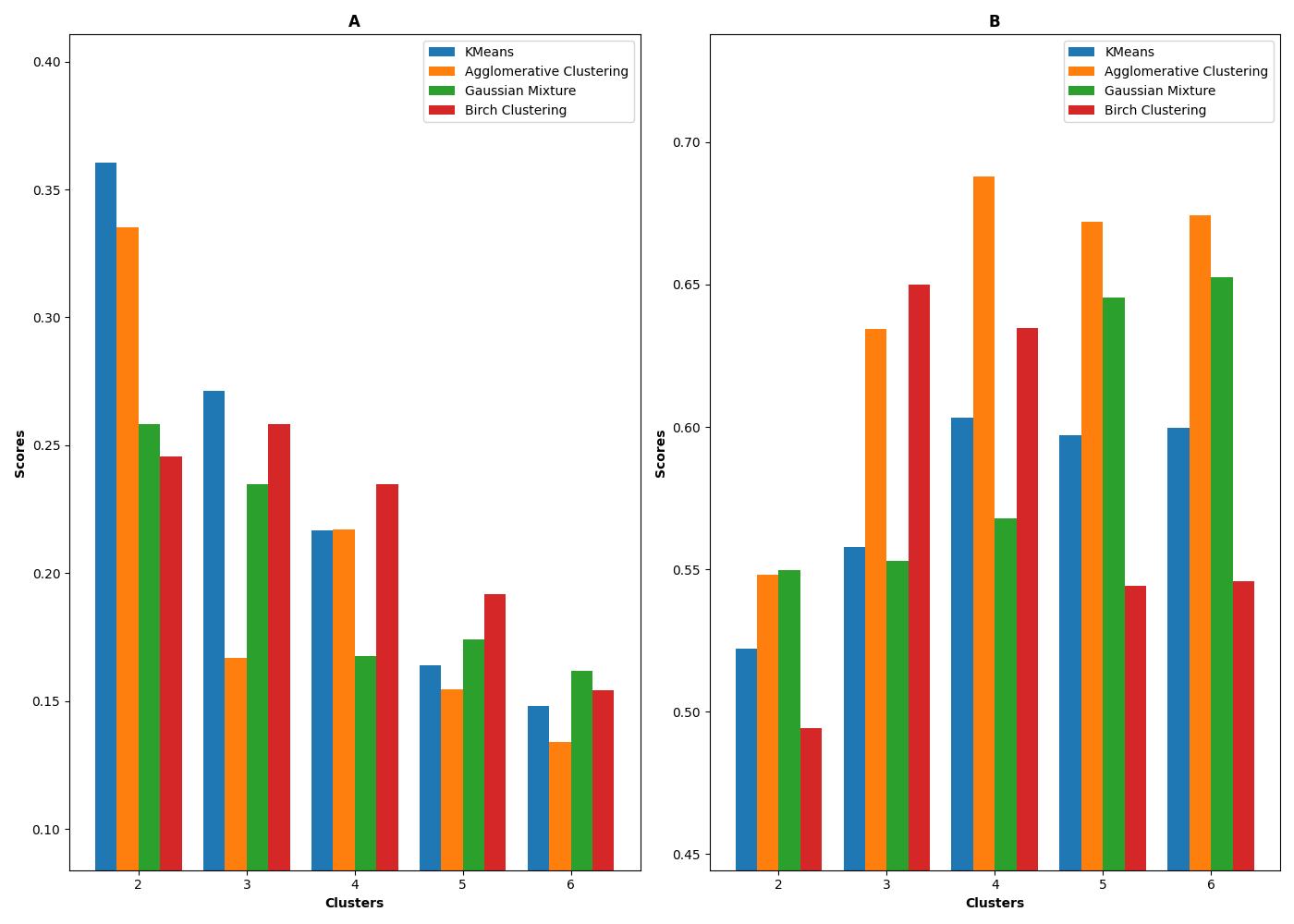

**Figure S18**: Overall scores based on our scoring function-1 (A) and scoring function-2 (B) based on the scores on rmsd and end-to-end distance properties shown in figure 8 for the protein-folding simulations. The weights for rmsd and end-to-end-distances are set to 0.3 and 0.7 respectively.

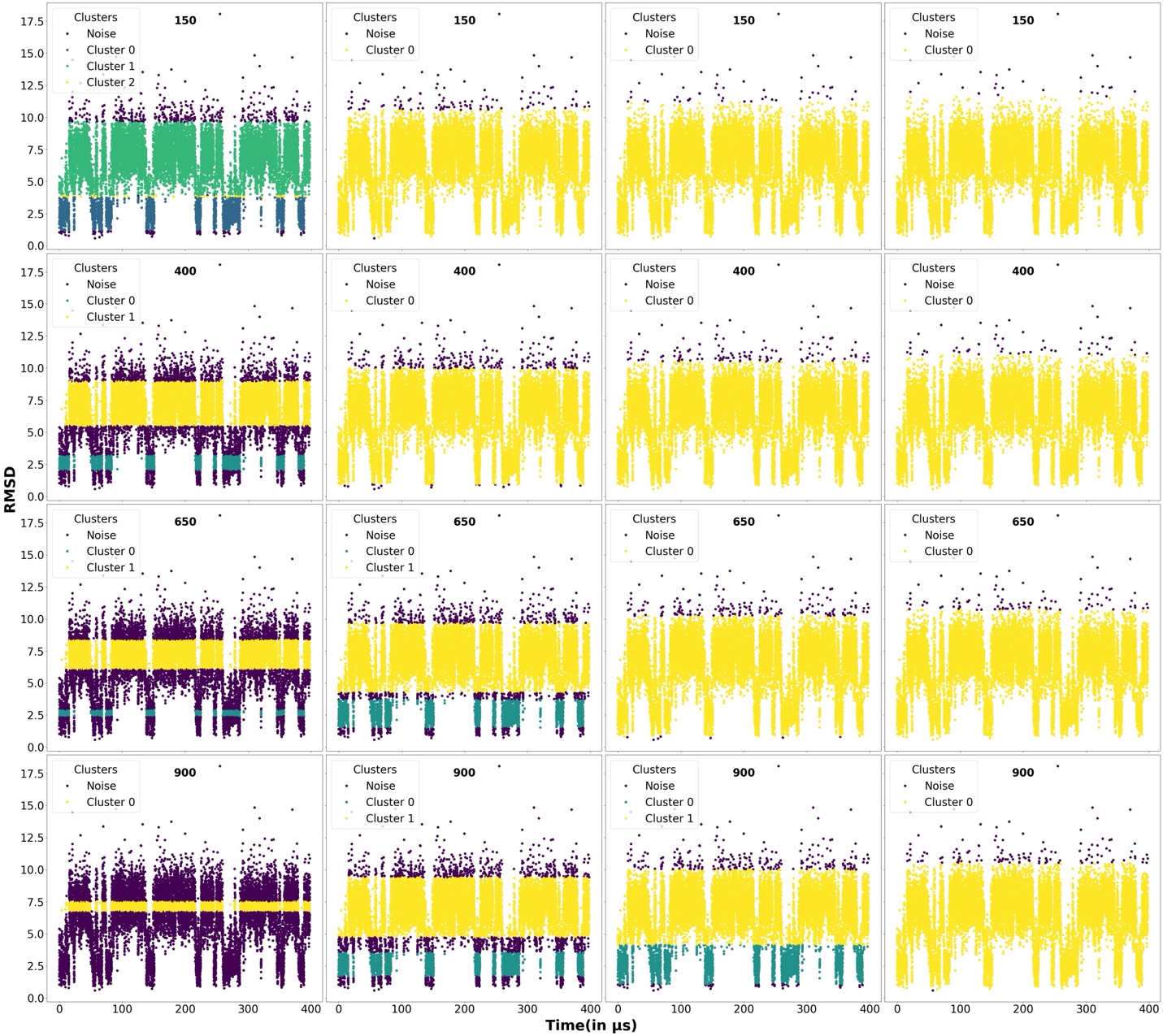

**Figure S19:** RMSD properties of the different protein-folding samples used for clustering by DBSCAN algorithm. Different clusters as well as the unclustered points(noise) are marked using different colors. The min-points used for the clustering are shown in labels. Each row represents the min-points used in labels with increasing ε values=0.1, 0.3, 0.5, and 0.7.

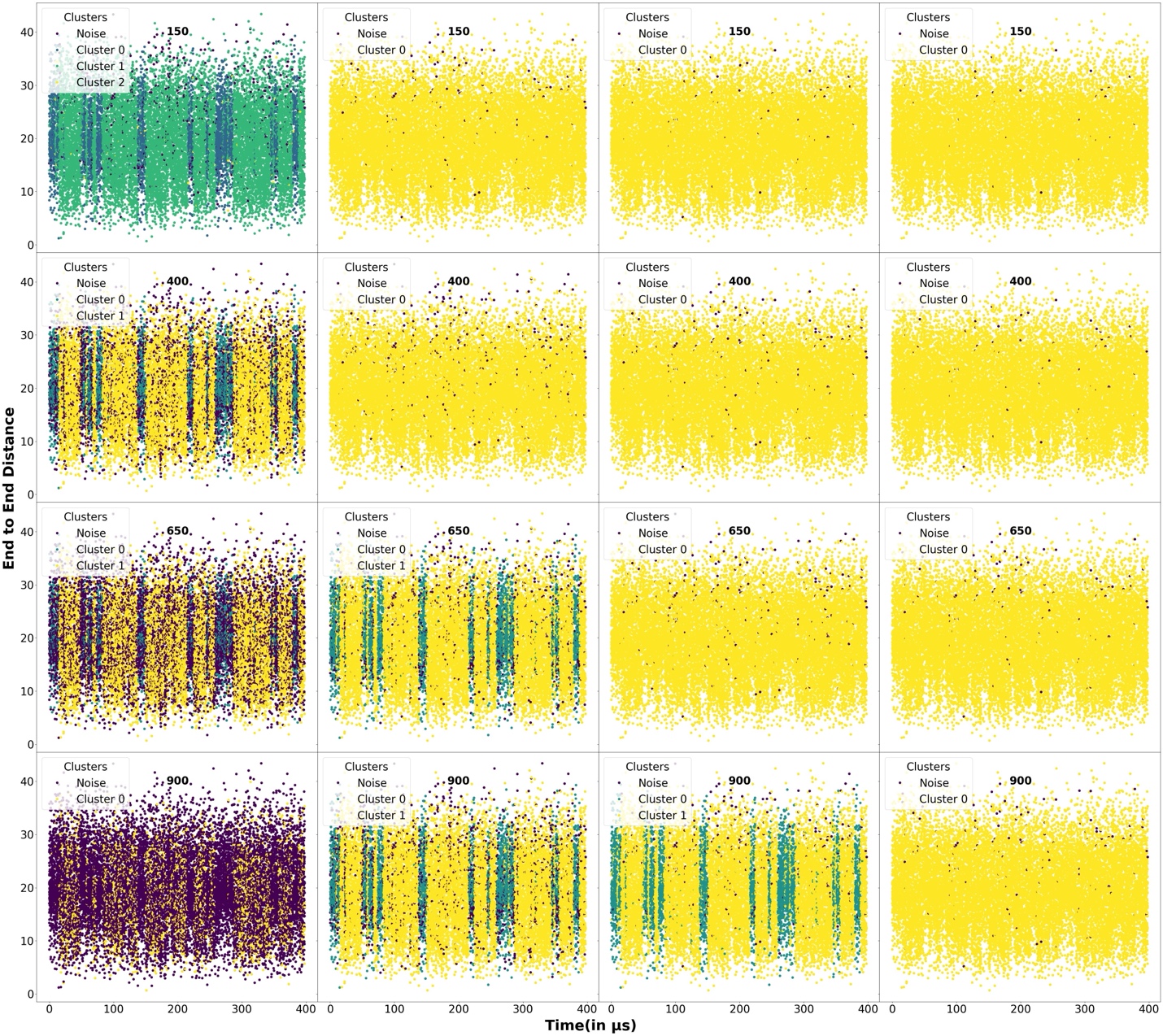

**Figure S20:** End-to-end-distance properties of the different protein-folding samples used for clustering by DBSCAN algorithm using our second approach. Different clusters as well as the unclustered points(noise) are marked using different colors. The min-points used for the clustering are shown in labels. Each row represents the min-points used in labels with increasing ε values=0.1, 0.3, 0.5, and 0.7.

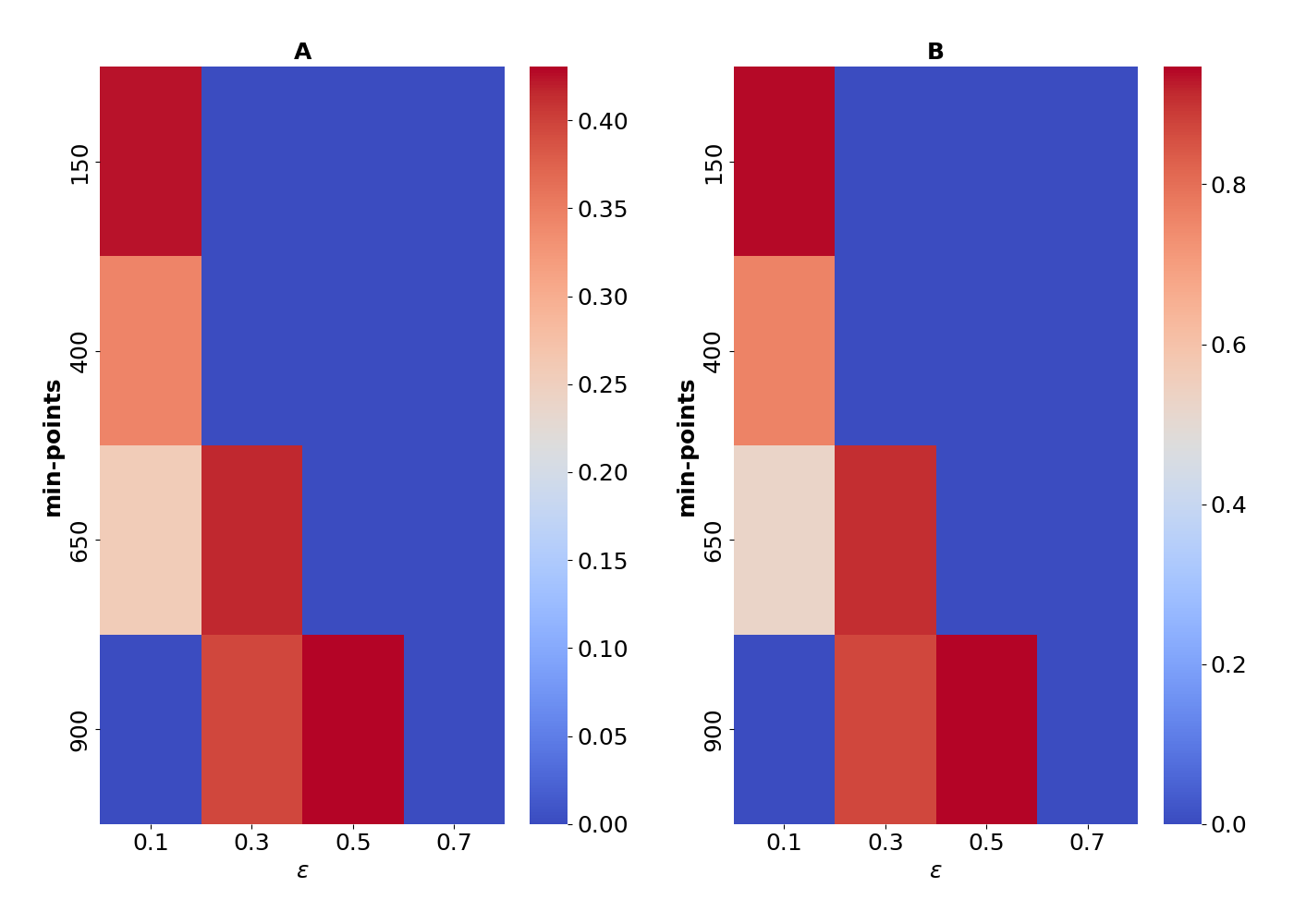

**Figure S21:** A two dimensional heatmap showing the scores obtained by our scoring metric-1(A) and scoring metric-2(B) for DBSCAN algorithm obtained by setting a weights of 0.30 to RMSD and 0.70 to end-to-end-distances to our overall scoring function(Equation-1 in the main manuscript).

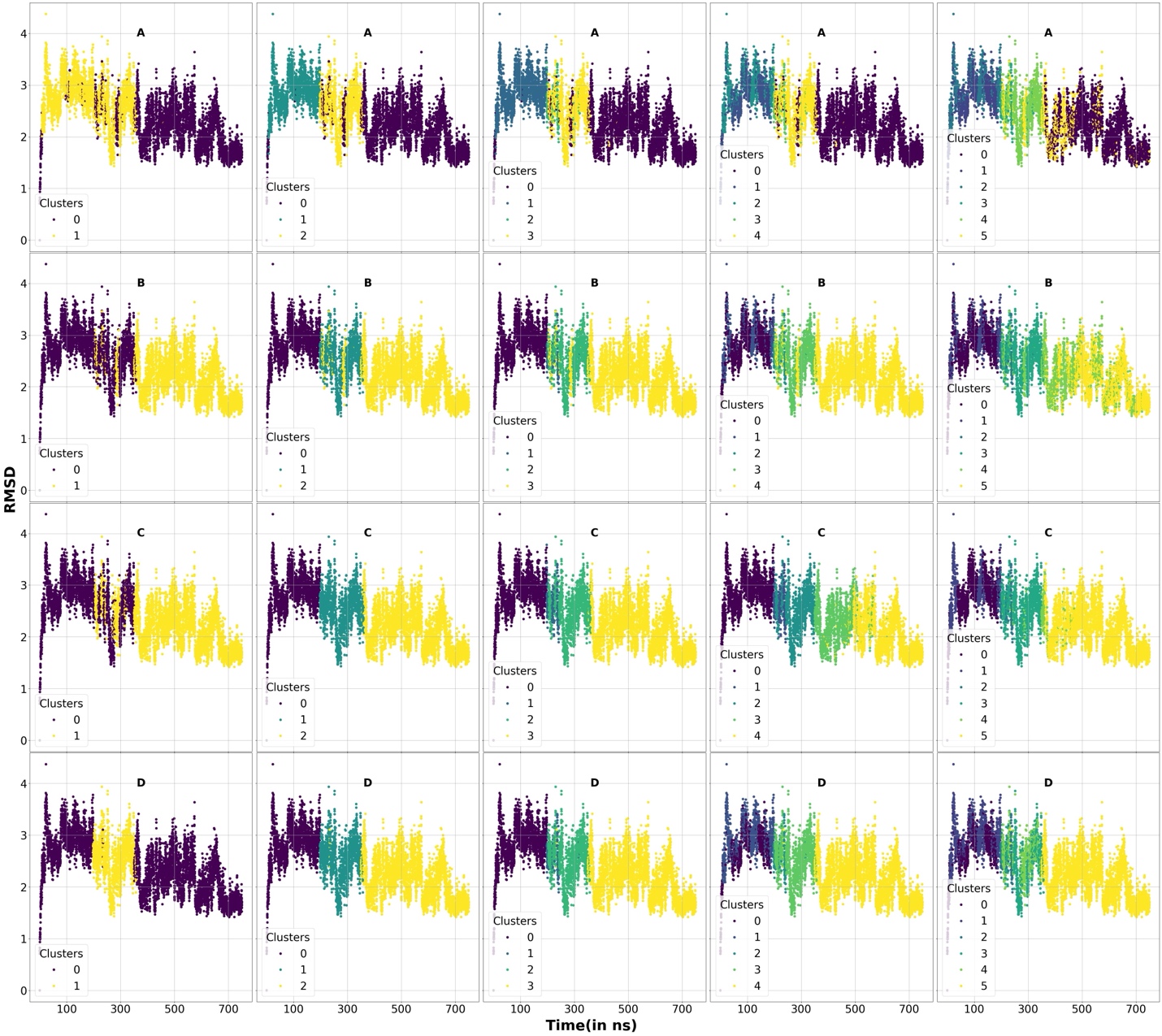

**Figure S22**: RMSD for the protein properties of the protein-ligand simulation trajectories used for clustering for Smlt1473 bound to Mana. Different clusters are marked using different colors. A, B, C, and D sets denotes our 4 clustering algorithms K-Means, Agglomerative, BIRCH, and Agglomerative. Each column respectively denotes 2,3,4,5, and 6 clusters.

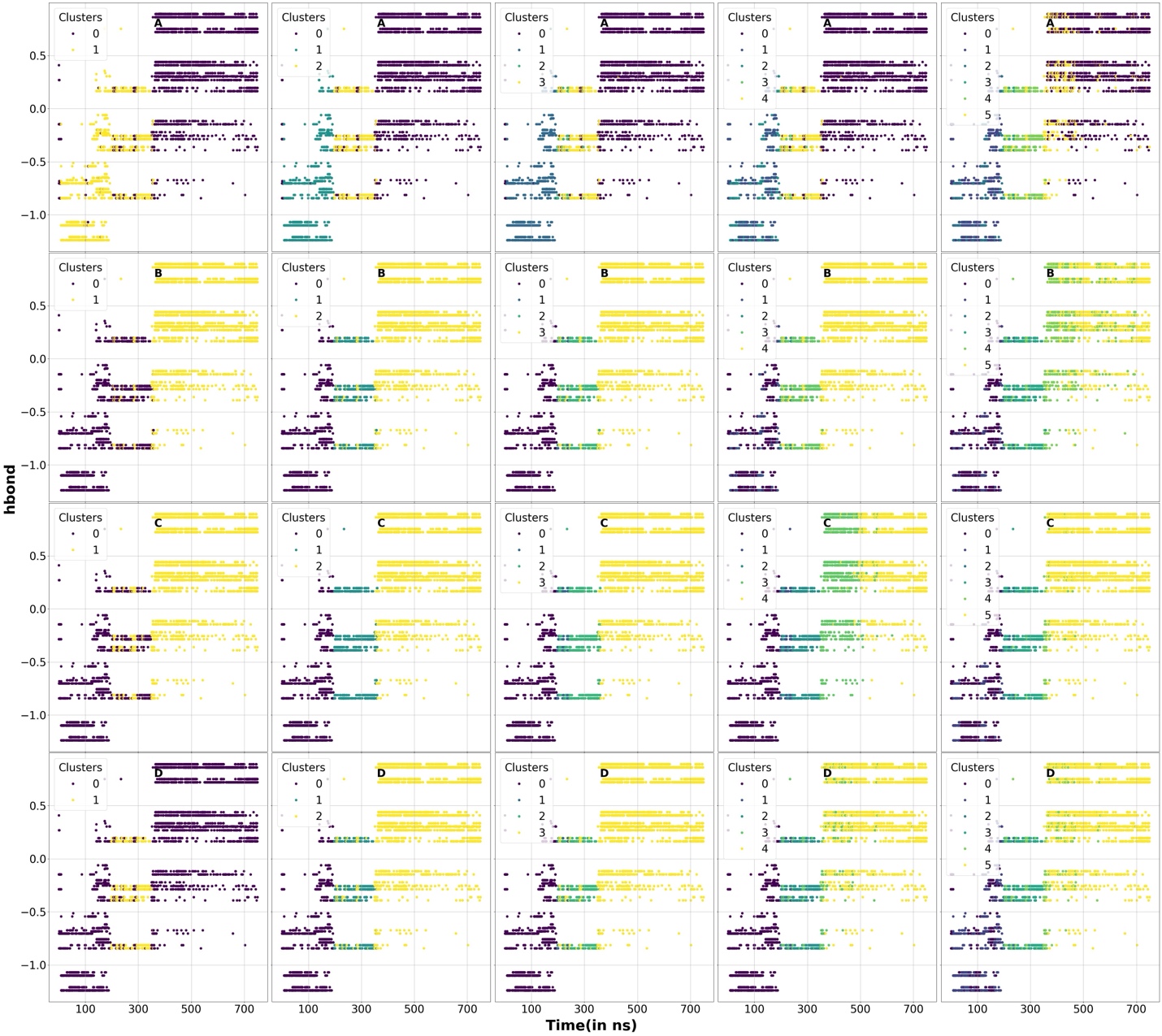

**Figure S23**: Reduced one-dimensional PCA component of presence or absence of hydrogen bonds between the protein and the sugar for the protein-ligand simulation trajectories used for clustering for Smlt1473 bound to Mana. Different clusters are marked using different colors. A, B, C, and D sets denotes our 4 clustering algorithms K-Means, Agglomerative, BIRCH, and Agglomerative. Each column respectively denotes 2,3,4,5, and 6 clusters.

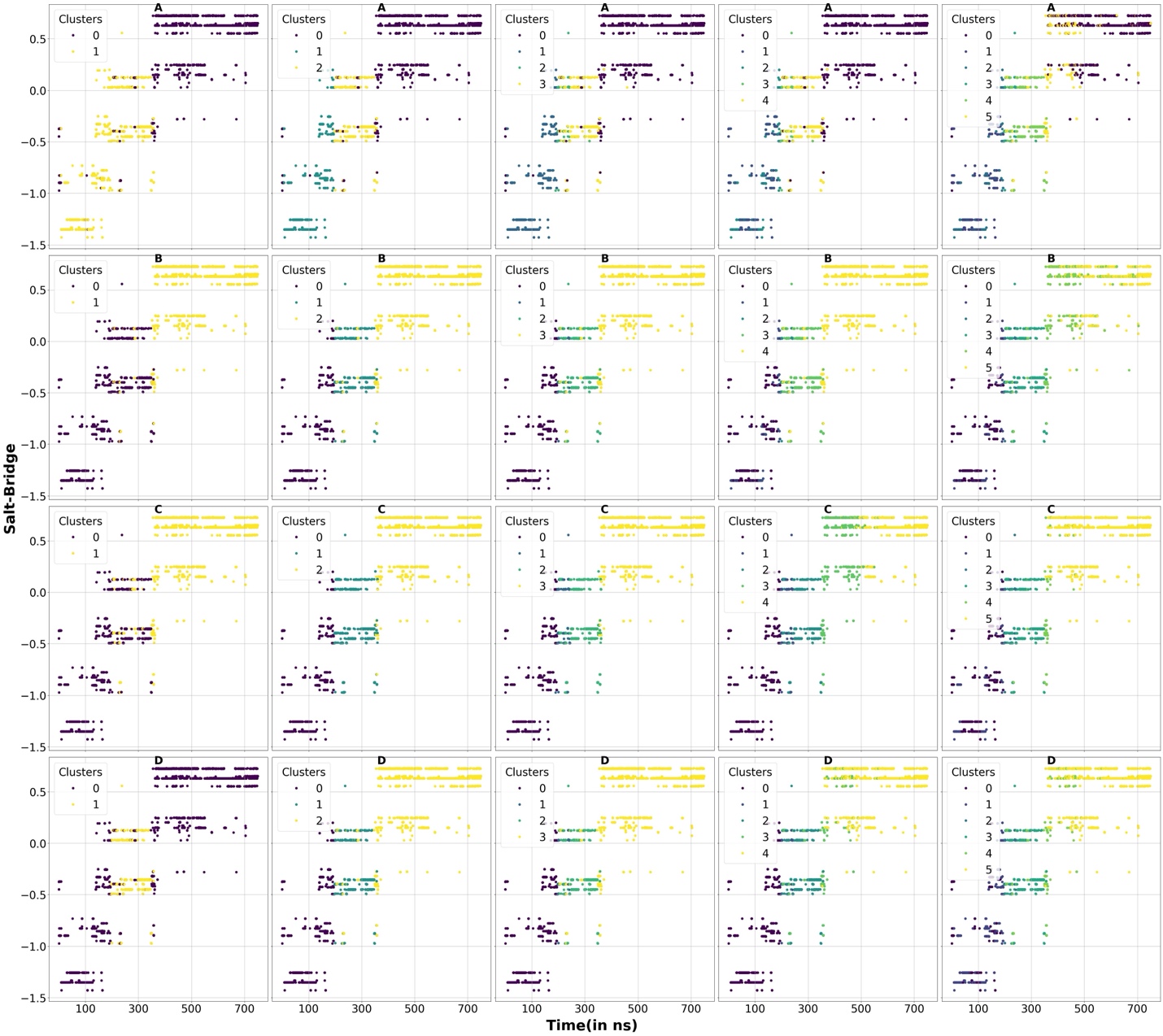

**Figure S24**: Reduced one-dimensional PCA component or presence or absence of salt bridges between the protein and the sugar for the protein-ligand simulation trajectories used for clustering for Smlt1473 bound to Mana. Different clusters are marked using different colors. A, B, C, and D sets denotes our 4 clustering algorithms K-Means, Agglomerative, BIRCH, and Agglomerative. Each column respectively denotes 2,3,4,5, and 6 clusters.

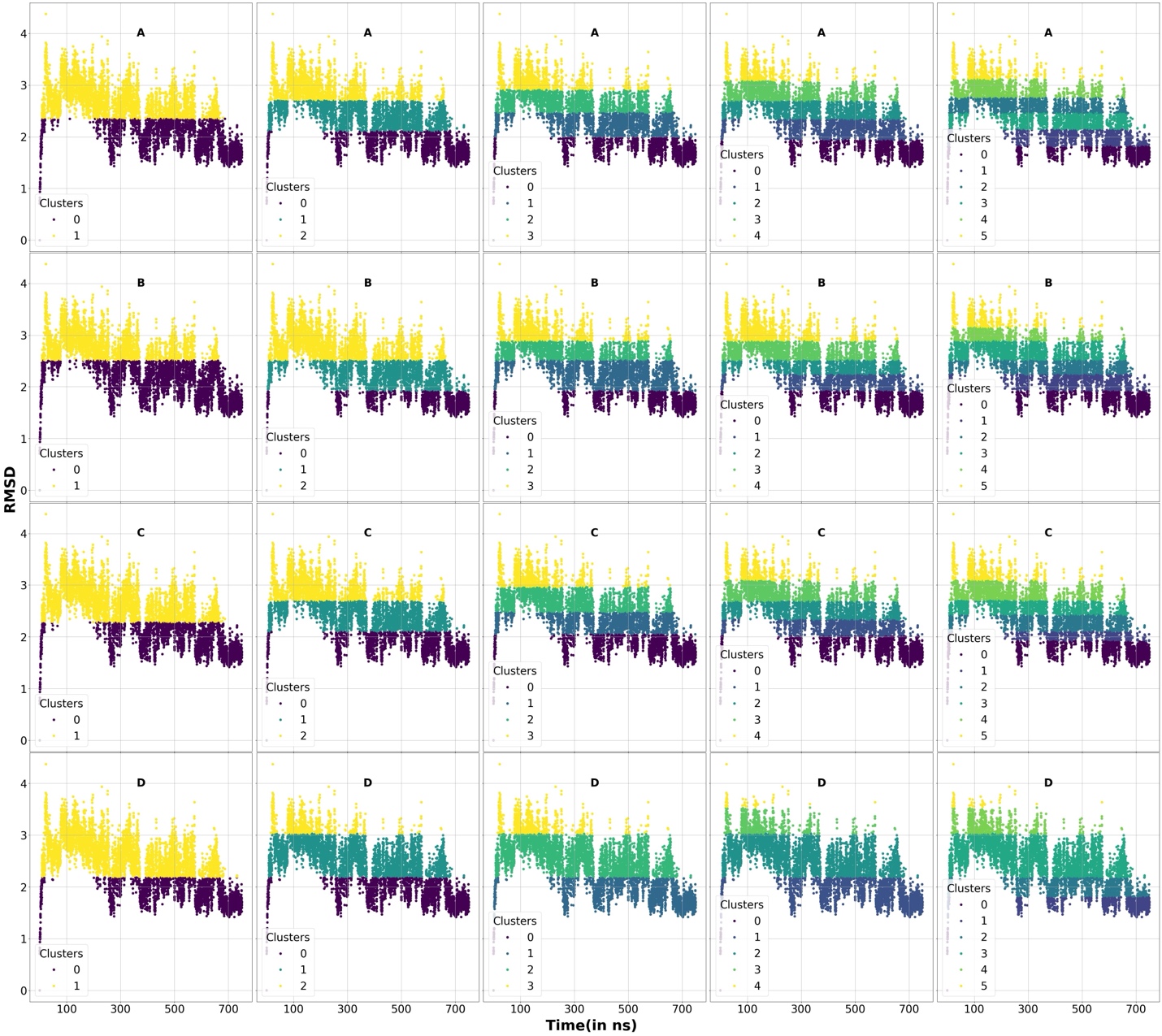

**Figure S25**: A graphical representation of the cluster states for the ground truth label for the RMSD of the protein backbone (for scoring metric-1) as a function of time for Smlt1473 bound to Mana. A, B, C, and D sets denotes our 4 clustering algorithms K-Means, Agglomerative, BIRCH, and Agglomerative. Each column respectively denotes 2,3,4,5, and 6 clusters.

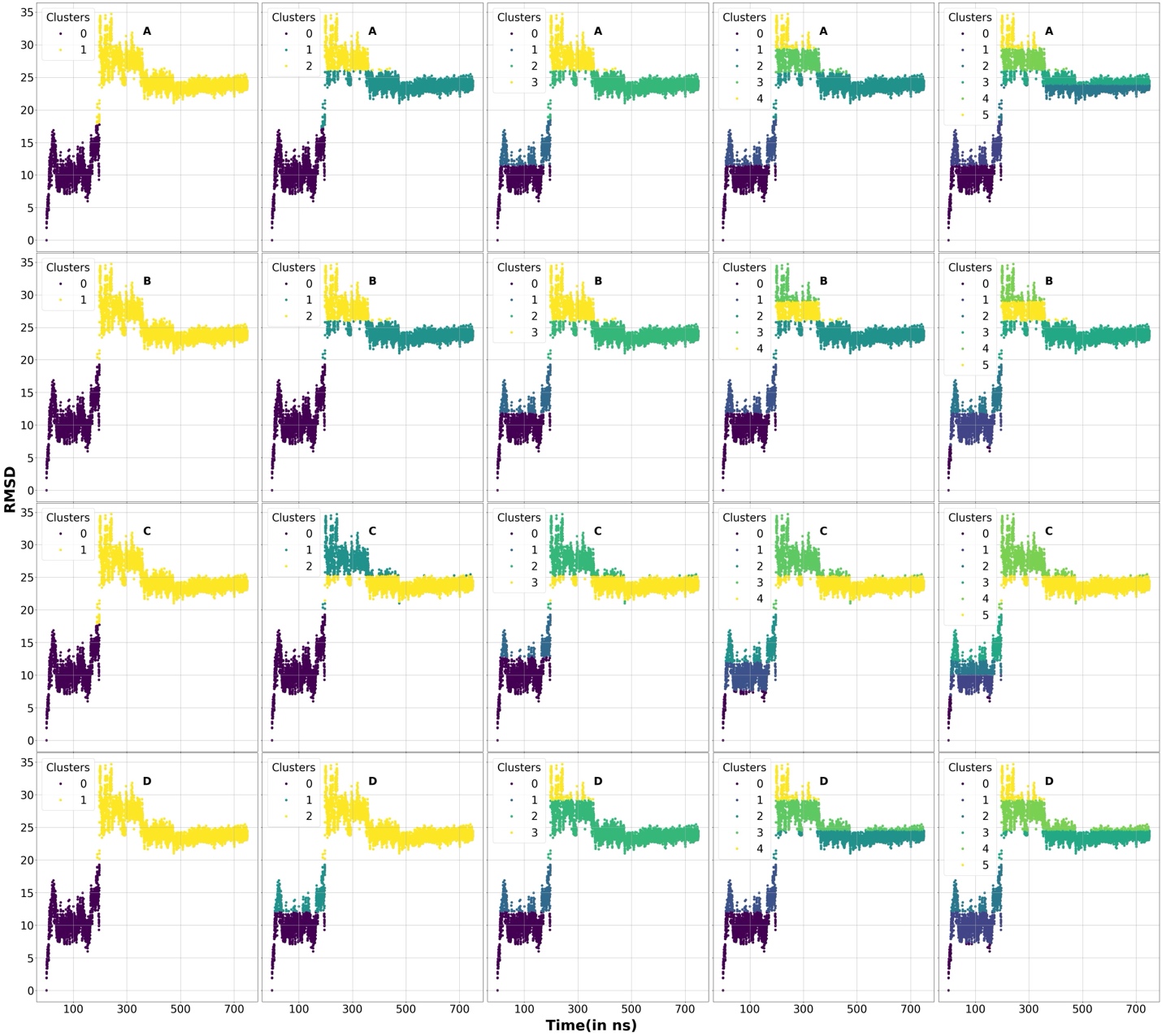

**Figure S26**: A graphical representation of the cluster states for the ground truth labels for the RMSD of the sugar backbone (for scoring metric-1) as a function of time for Smlt1473 bound to Mana. A, B, C, and D sets denotes our 4 clustering algorithms K-Means, Agglomerative, BIRCH, and Agglomerative. Each column respectively denotes 2,3,4,5, and 6 clusters.

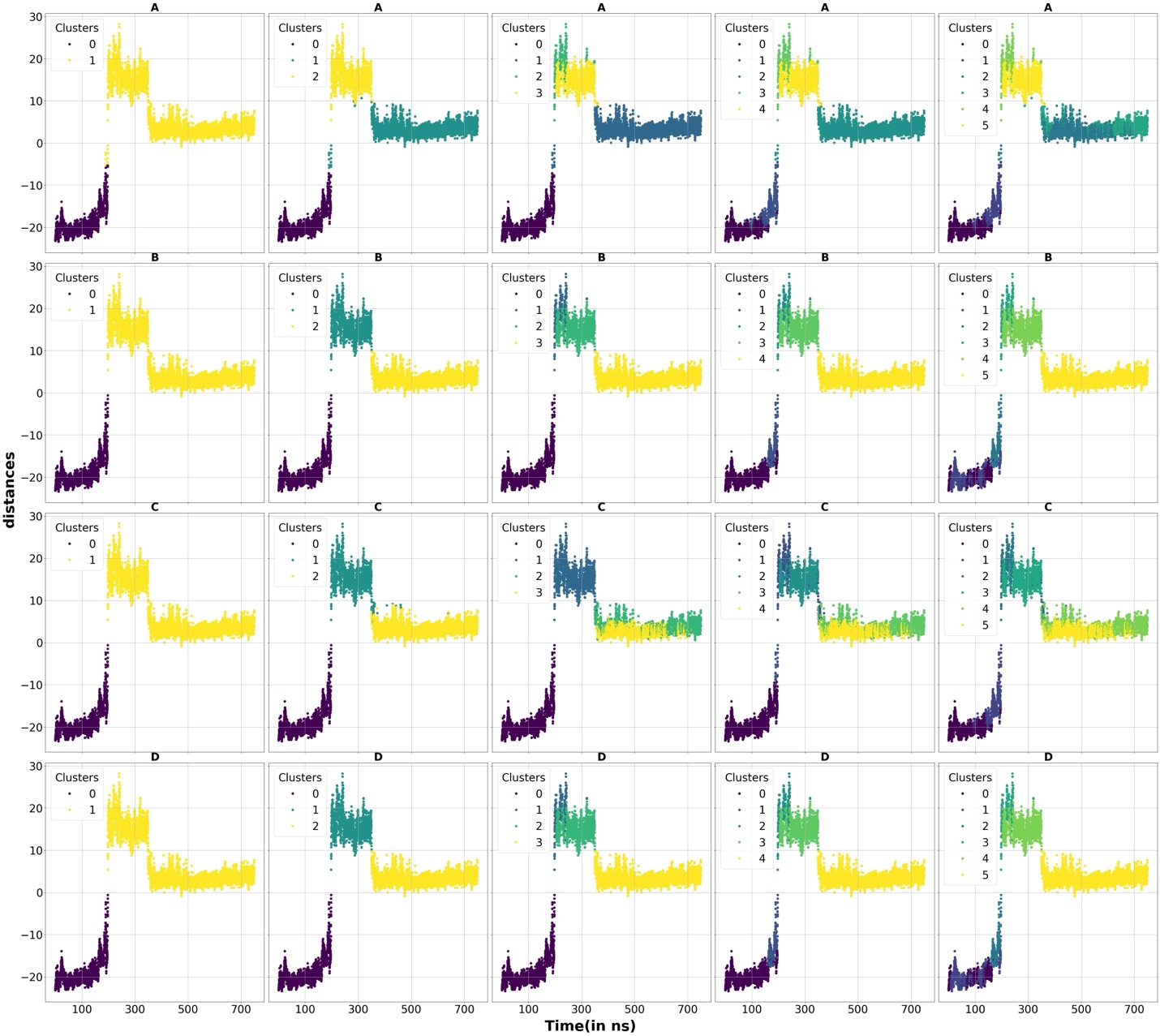

**Figure S27**: A graphical representation of the cluster states for the ground truth labels for the distances-resolved into one principal component (for scoring metric-1) as a function of time for Smlt1473 bound to Mana. A, B, C, and D sets denotes our 4 clustering algorithms K-Means, Agglomerative, BIRCH, and Agglomerative. Each column respectively denotes 2,3,4,5, and 6 clusters.

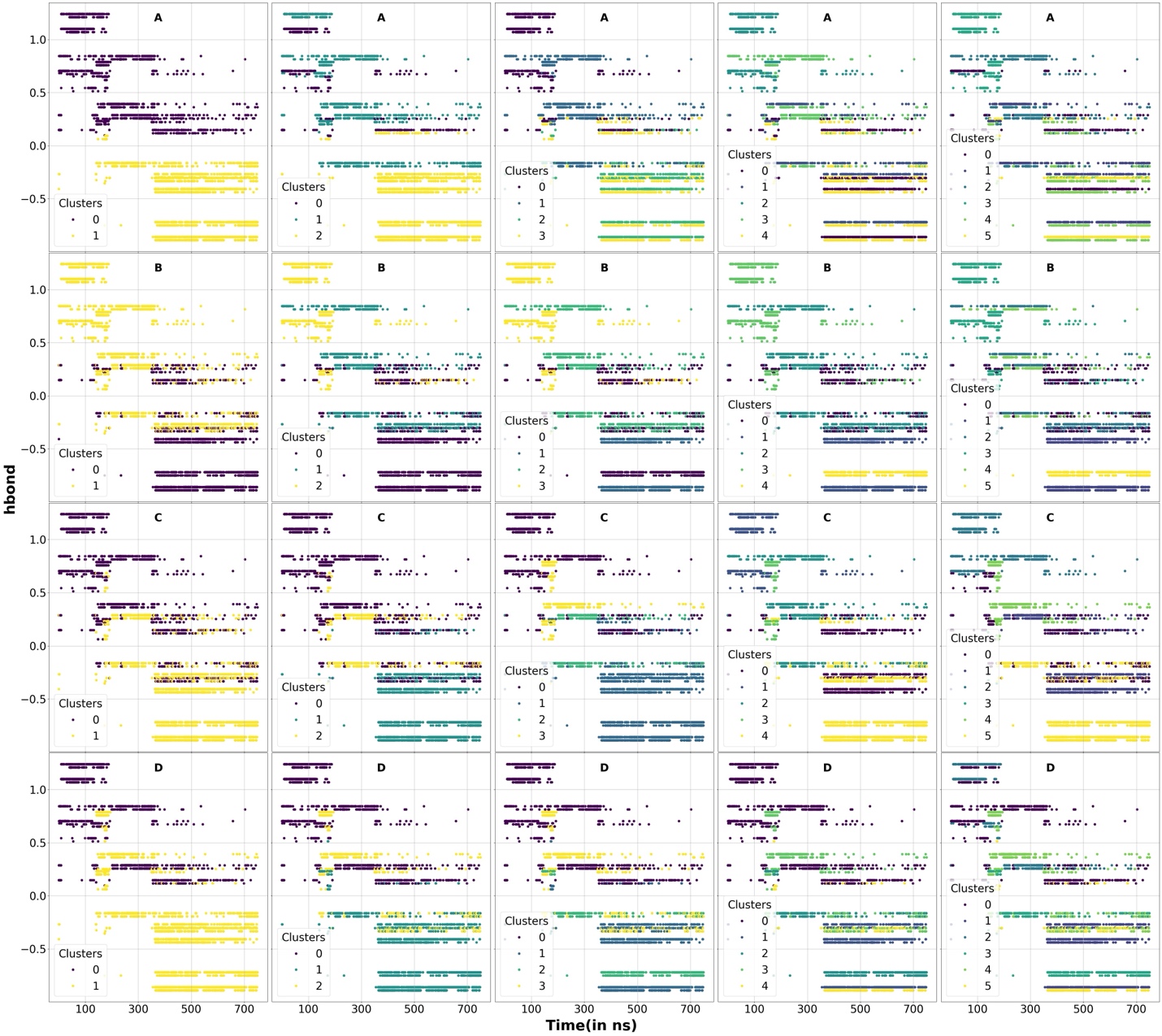

**Figure S28**: A graphical representation of the cluster states for the ground truth labels for the hydrogen-bonds resolved into one principal component (for scoring metric-1) as a function of time for Smlt1473 bound to Mana. A, B, C, and D sets denotes our 4 clustering algorithms K-Means, Agglomerative, BIRCH, and Agglomerative. Each column respectively denotes 2,3,4,5, and 6 clusters.

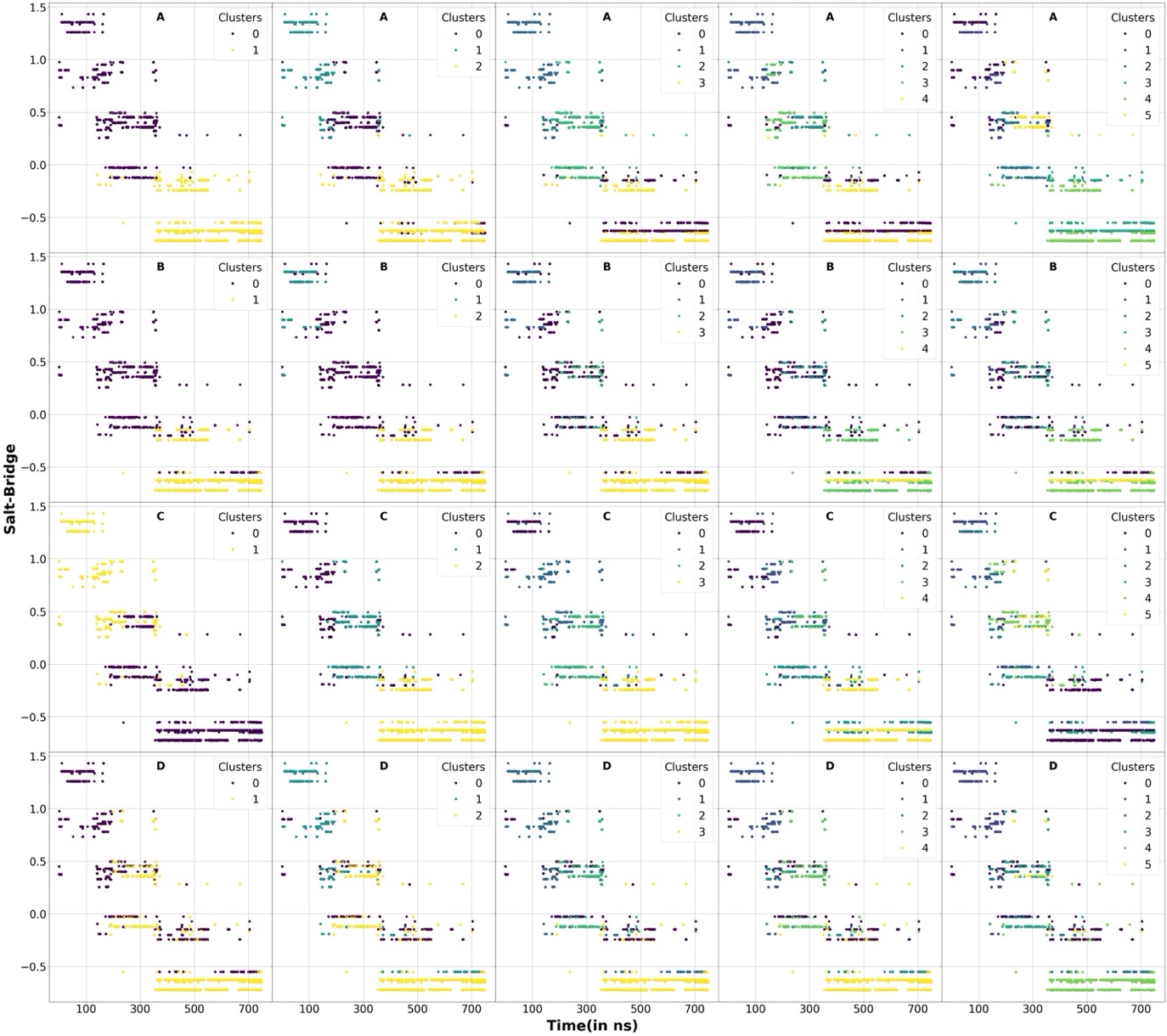

**Figure S29**: A graphical representation of the cluster states for the ground truth labels for the salt-bridges resolved into one principal component (for scoring metric-1) as a function of time for Smlt1473 bound to Mana. A, B, C, and D sets denotes our 4 clustering algorithms K-Means, Agglomerative, BIRCH, and Agglomerative. Each column respectively denotes 2,3,4,5, and 6 clusters.

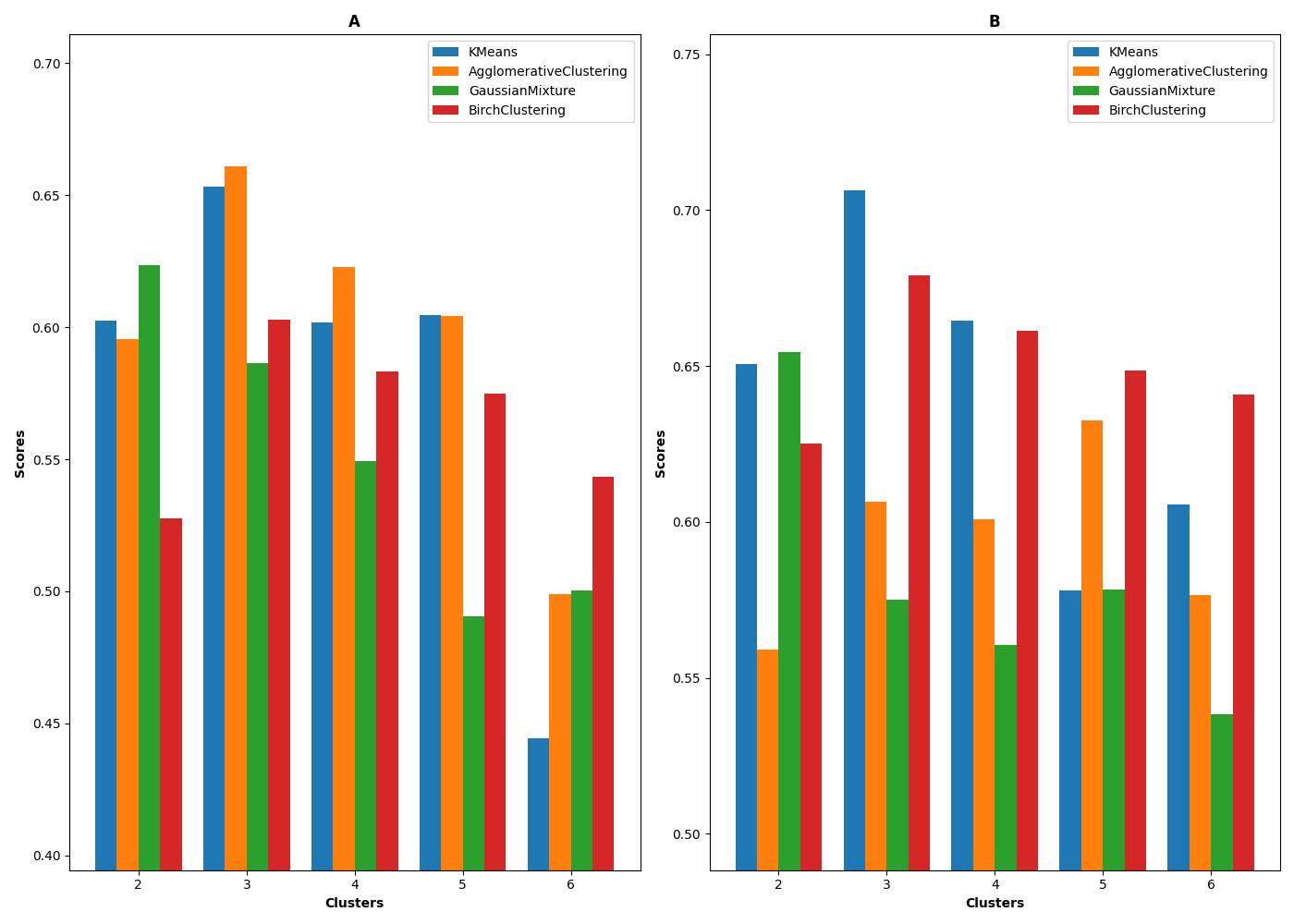

**Figure S30**: Overall scores based on our scoring function-1 (A) and scoring function-2 (B) based on the scores on RMSD of the sugar, distance properties of the sugar from the protein, as well as the presence and absence of hydrogen bonds/salt-bridges shown in figure 15 for the protein-ligand simulations for Smlt1473 bound to Mana. The weights W_rmsd-sugar_, W_distances_, W_presence-of-h-bonds_, and W_presence-of-salt-bridges_ are set to 0.35 and 0.35, 0.15, and 0.15 respectively.

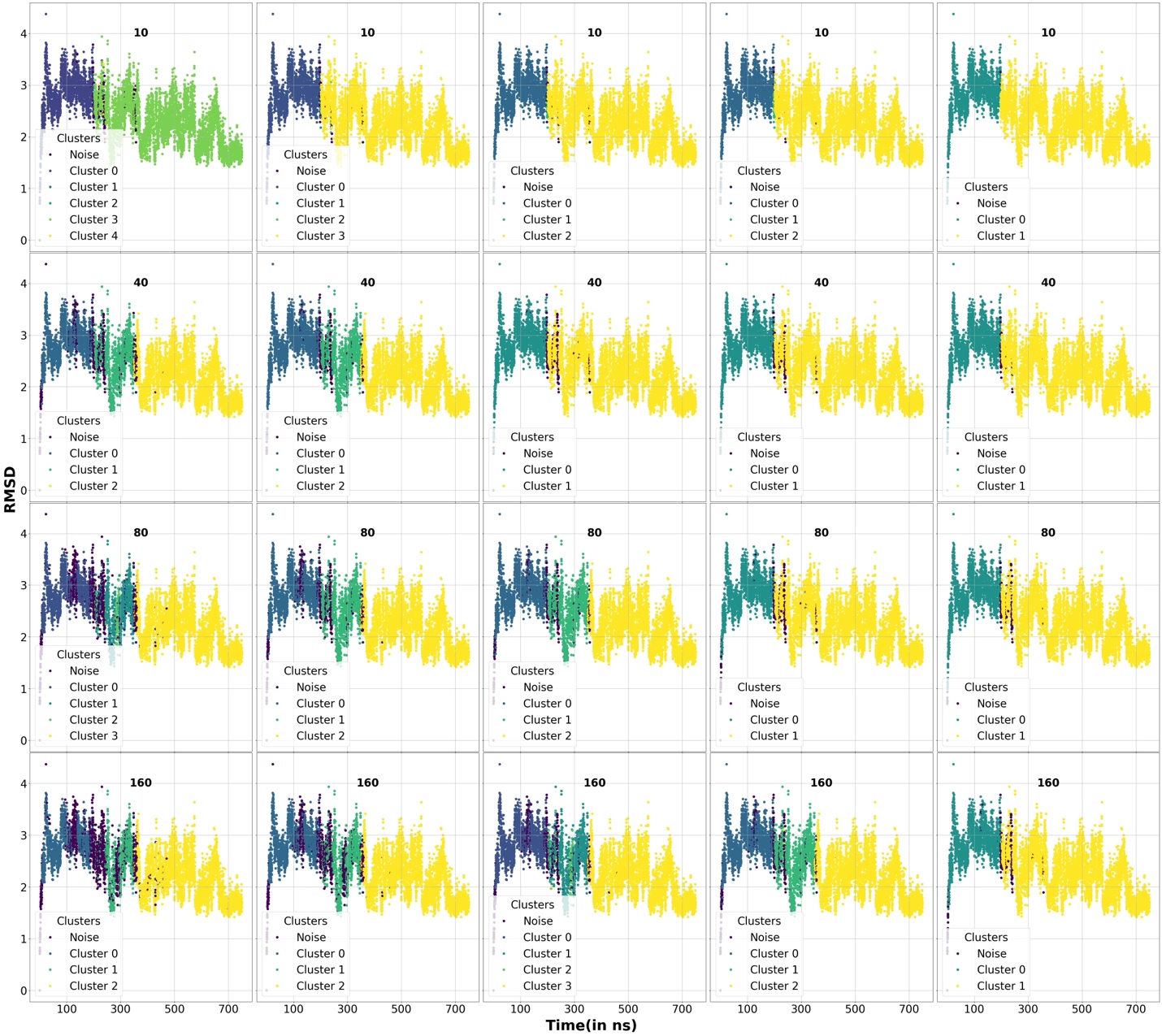

**Figure S31:** RMSD of the protein properties of the different protein-ligand samples used for clustering by DBSCAN algorithm for the Sml1473 bound to Mana system. Different clusters as well as the unclustered points(noise) are marked using different colors. The min-points used for the clustering are shown in labels. Each row represents the min-points used in labels with increasing ε values=0.04, 0.05, 0.06, 0.07, and 0.09.

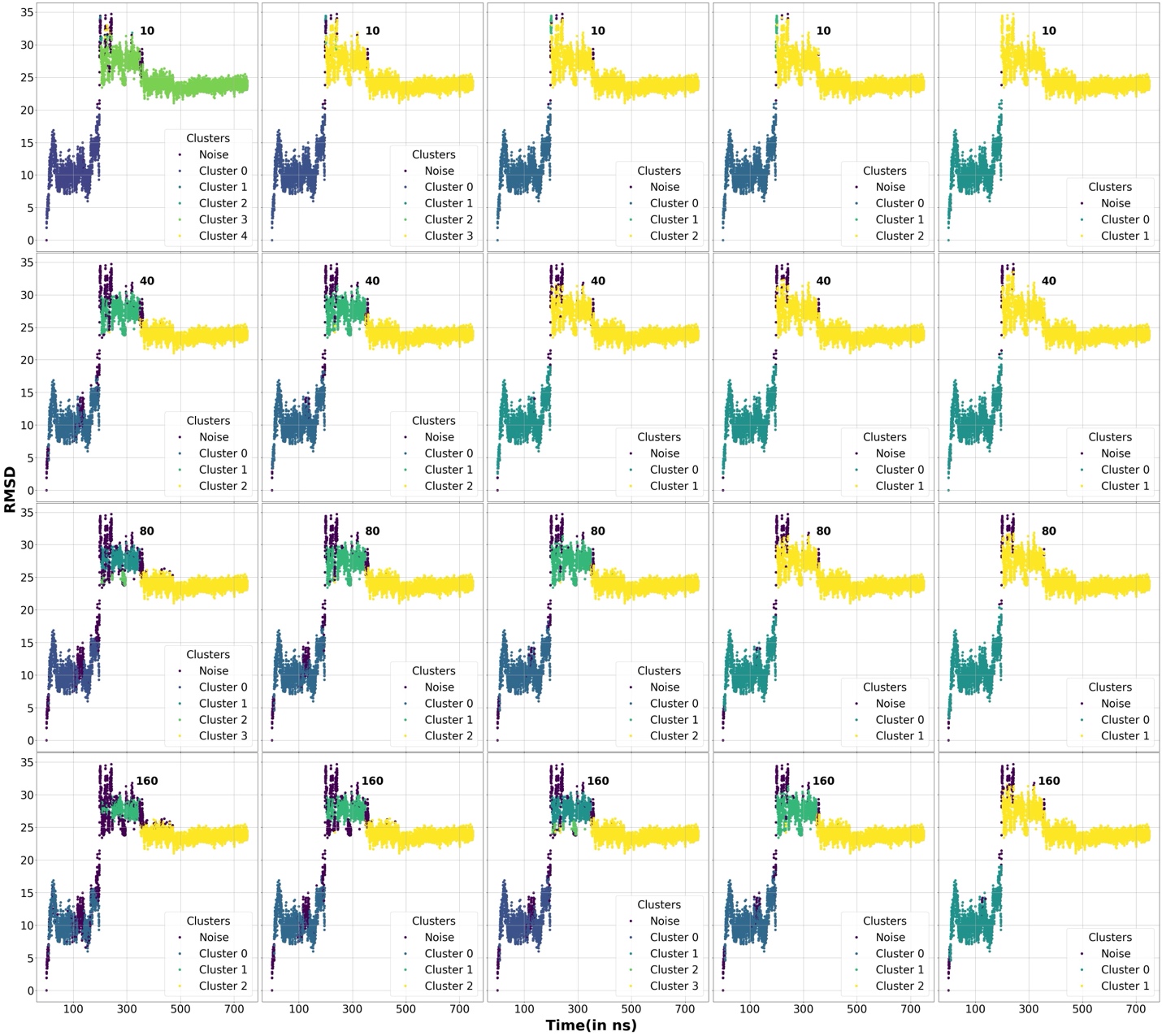

**Figure S32:** RMSD of the ligand properties of the different protein-ligand samples used for clustering by DBSCAN algorithm for the Smlt1473 bound to Mana system. Different clusters as well as the unclustered points(noise) are marked using different colors. The min-points used for the clustering are shown in labels. Each row represents the min-points used in labels with increasing ε values=0.04, 0.05, 0.06, 0.07, and 0.09.

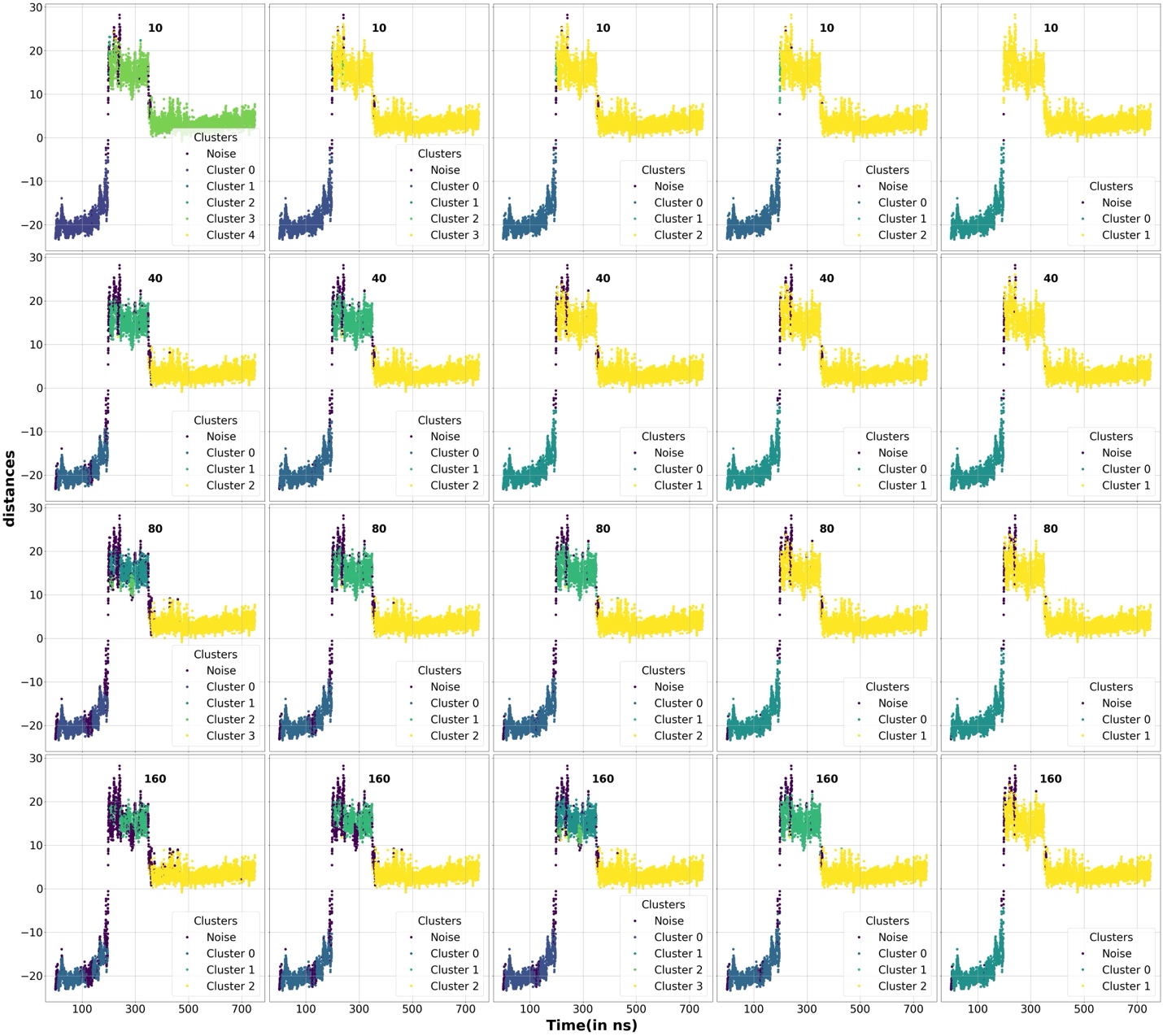

**Figure S33:** A dimensional PCA component of the distance properties of the different protein-ligand samples used for clustering by DBSCAN algorithm for the Sml1473 bound to Mana system. Different clusters as well as the unclustered points(noise) are marked using different colors. The min-points used for the clustering are shown in labels. Each row represents the min-points used in labels with increasing ε values=0.04, 0.05, 0.06, 0.07, and 0.09.

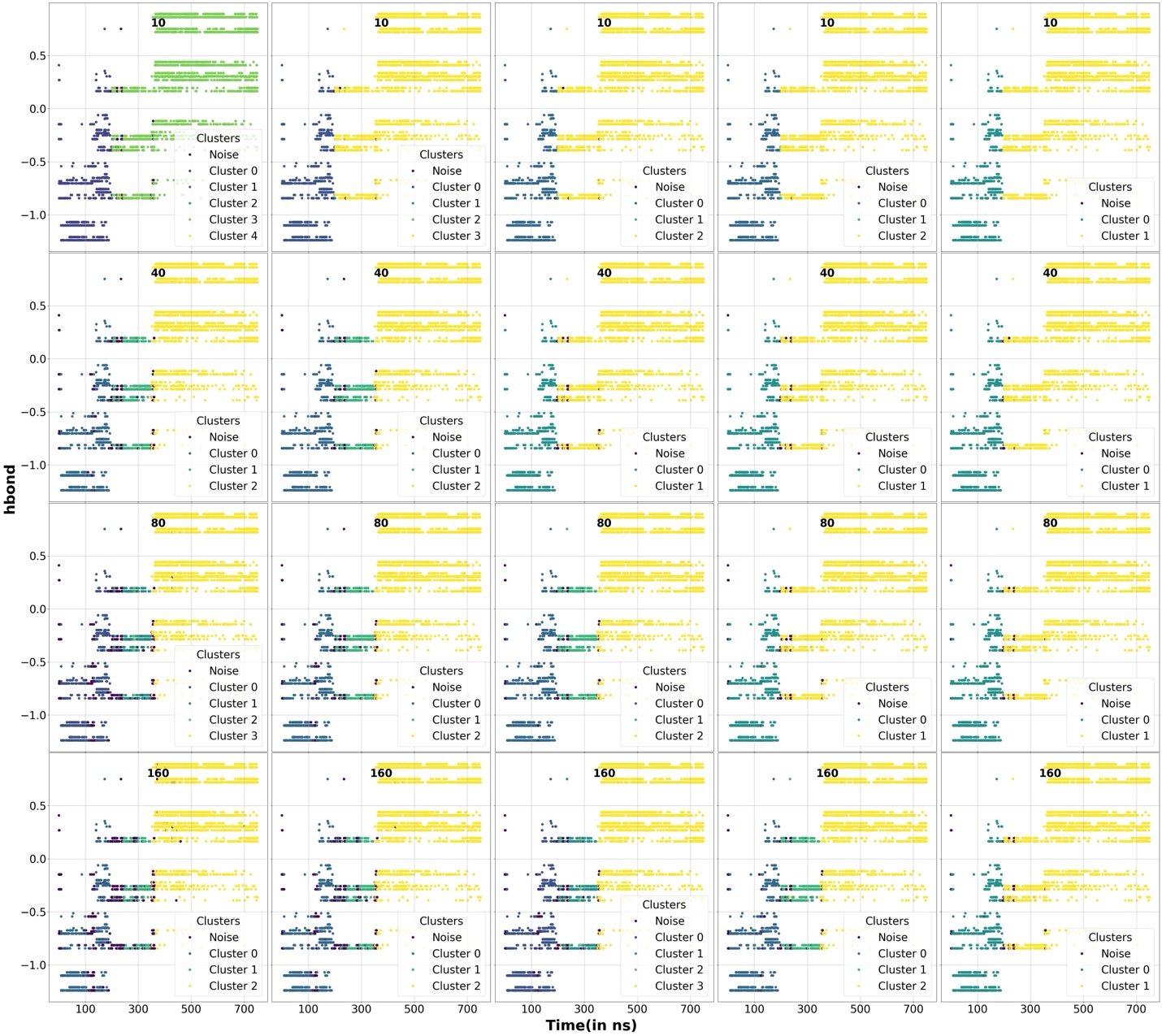

**Figure S34:** A dimensional PCA component of the hydrogen-bond properties of the different protein-ligand samples used for clustering by DBSCAN algorithm for the Smlt1473 bound to Mana system. Different clusters as well as the unclustered points(noise) are marked using different colors. The min-points used for the clustering are shown in labels. Each row represents the min-points used in labels with increasing ε values=0.04, 0.05, 0.06, 0.07, and 0.09.

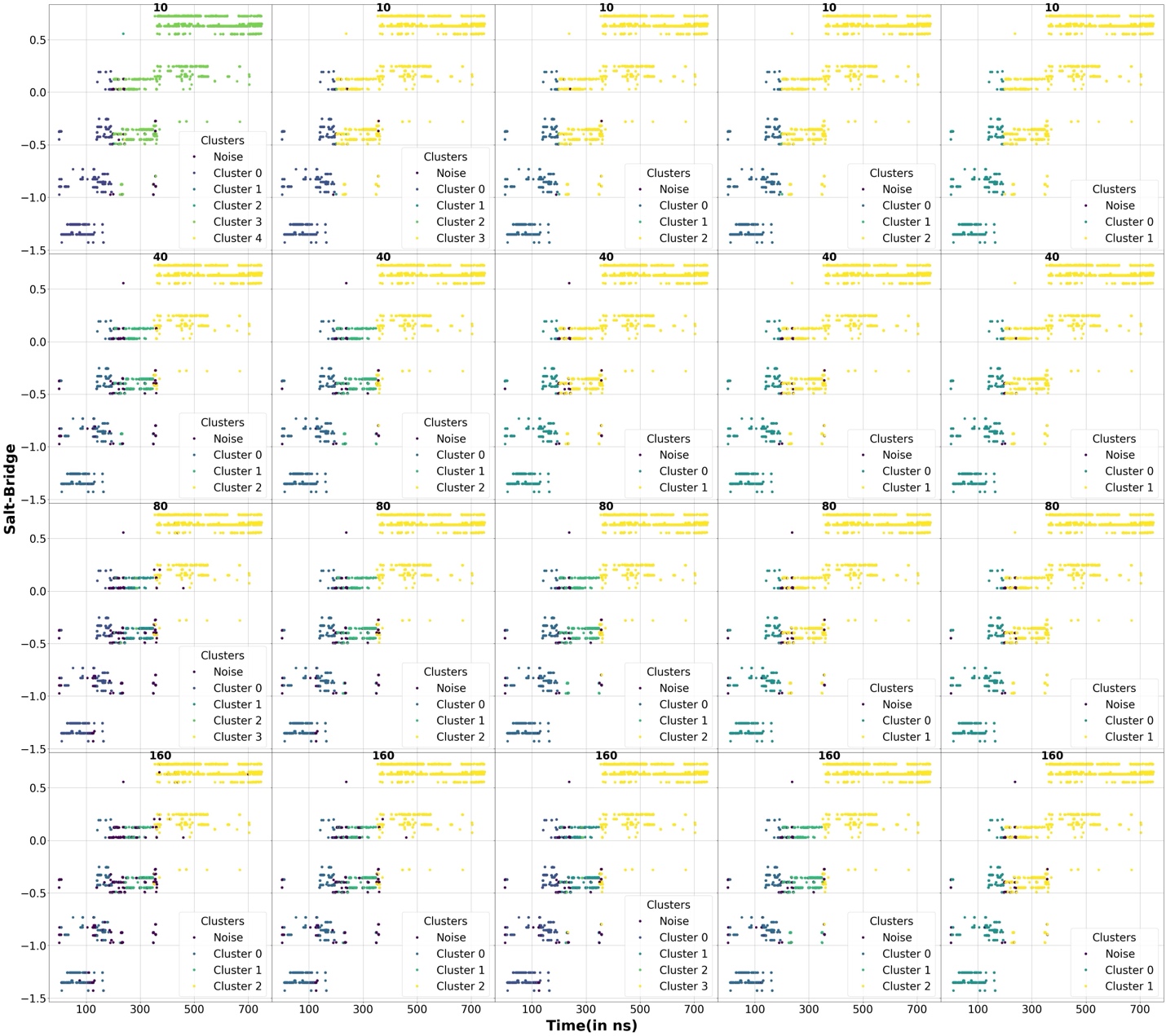

**Figure S35:** A dimensional PCA component of the salt-bridge properties of the different protein-ligand samples used for clustering by DBSCAN algorithm for the Smlt1473 bound to Mana system. Different clusters as well as the unclustered points(noise) are marked using different colors. The min-points used for the clustering are shown in labels. Each row represents the min-points used in labels with increasing ε values=0.04, 0.05, 0.06, 0.07, and 0.09.

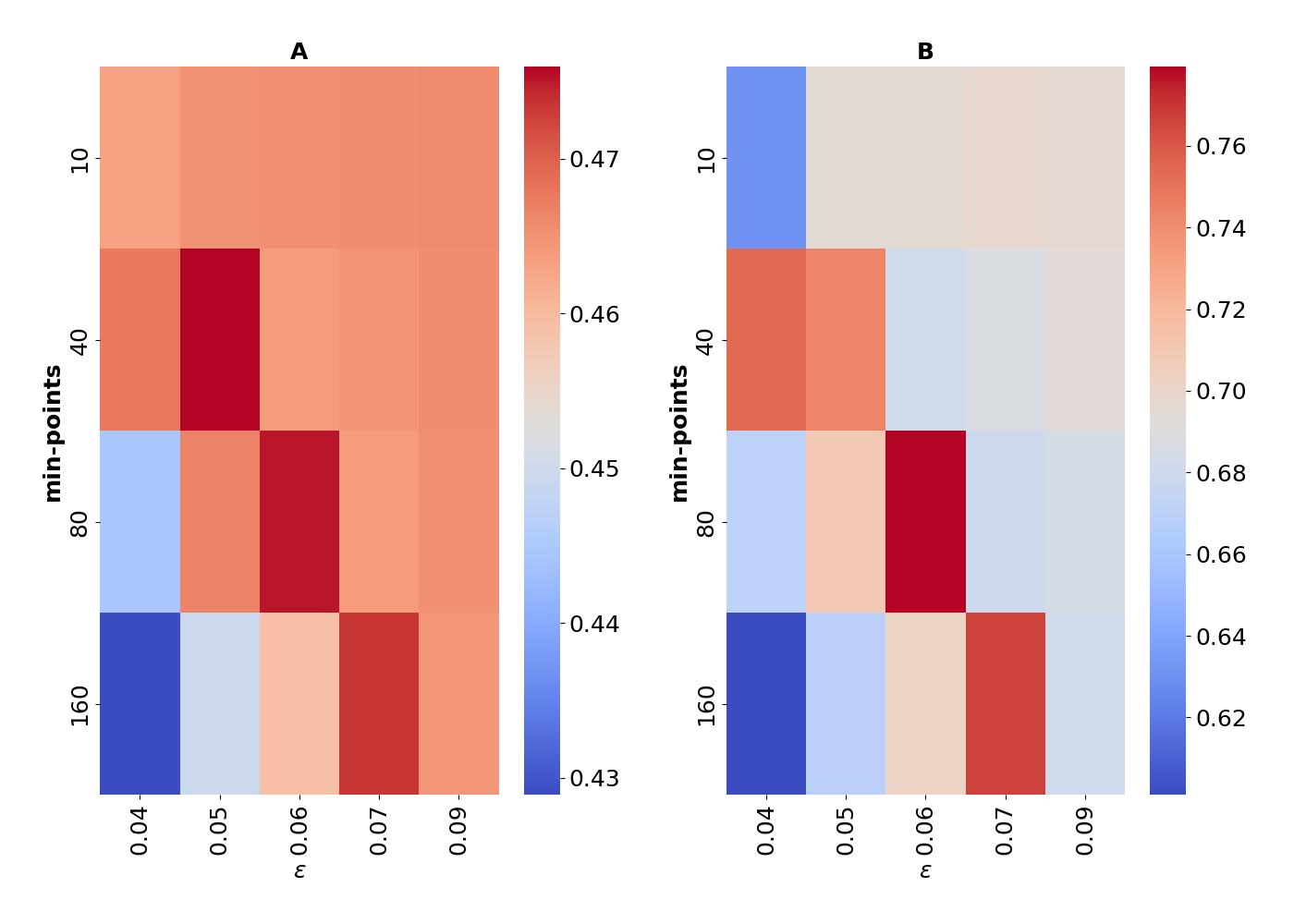

**Figure S36:** A two dimensional heatmap showing the scores obtained by our scoring metric-1(A) and scoring metric-2(B) for DBSCAN algorithm obtained by setting a weights of W_rmsd-sugar_=0.35, W_distances_=0.35, W_presence-of-h-bonds_=0.15, and W_presence-of-salt-bridges_=0.15 to our overall scoring function(Equation-1 in the main manuscript) for Smlt1473 bound to Mana.

**Analysis on a protein-ligand dissociated trajectory**

As a final system of interest, we considered a full protein-ligand dissociated trajectory, we considered a FKBP protein bound to DMSO (PDB ID-1D7H). This system is small in size making it an attractive system to study by many MD method development studies^1-3^. For our interest, this ligand bound to the protein has a very short residence time, making it an attractive system to analyze using our methods. We simulated the system in Gromacs after building the system using CHARMM-GUI. Further details of the system preparation are provided in Reference 3^3^. After simulating the system, we see that the ligand dissociates from the protein after around 20 ns of simulation.

To convert the raw simulated trajectory into an usable ML form, we collected the distances between all the residues of the protein and the center of mass of the ligand to form a one-dimensional 106 length array representing the protein-ligand state at a particular time step. Next we combine the data across all the time steps to form a 16000*106 dimensional array. Next we run our clustering algorithms on top of this matrix.

The various properties that we chose again for our clustering the RMSD of the protein, RMSD of the sugar, and the distances between the various important residues on the protein to the sugar. We followed the same methods as we used earlier to compute the various properties in Ref 4^4^. We again show the various properties in figure S37, namely the RMSD of the protein, the RMSD of the ligand, and the distances of the most important residues on the protein to the ligand. In figures S38-S40, we present the cluster labels obtained for the various properties respectively. In figures S42-S44, we present the ground truth labels obtained for the various properties. In figure S41, we show the various scores obtained by our two metrics on the various properties. Finally, in figure S45, we show the overall scores obtained by setting a weight of 0.5 equally on the rmsd of the sugar as well as the distance property. This is because based on our physical intuitions, RMSD of the ligand might be equally as important as the distance of the protein from the ligand. We show that our overall scoring algorithm is able to get two clusters mainly for the associated and the dissociated state as shown by the RMSD of the sugar in figure S39 and S40 by two clusters for KMeans/Agglomerative clustering.

Finally, we also show how we can find out the optimal min-points and ε for DBSCAN algorithm using our scoring metrics in figures S46-S49. We show the various clusters obtained by DBSCAN algorithms for our various properties in figures S46-S48 respectively. In figure S49, we show the heatmap that shows the various scores obtained by our scoring algorithm. We show that two clusters is obtained for min-points=40 and ε=20.0(by setting equal weights W_rmsd-sugar_=0.50 and W_distances_=0.50 to our overall scoring function) which is also what we get when we physically look at figure S47 and S48 for the optimal clustering using DBSCAN. Again, we would like to point out here that it would be unfair to compare DBSCAN against the other algorithms as our scoring algorithms are different for DBSCAN against the other algorithms..

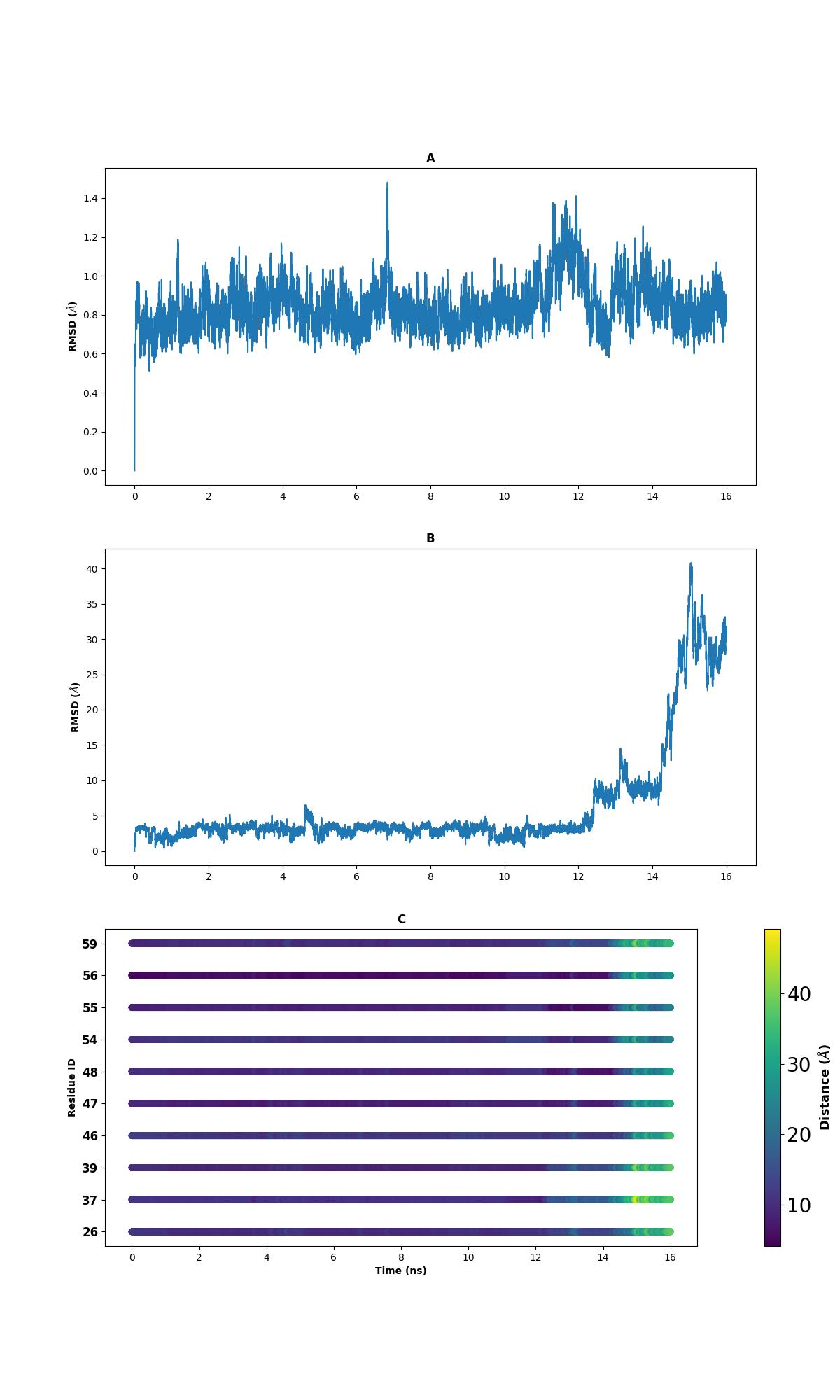

**Figure S37:** The various properties of the protein ligand simulation trajectories for FKBP protein bound/unbound to DMSO that we have. (A) denotes the RMSD of the protein plotted as a function of time. (B) denotes the RMSD of the ligand as a function of time. (C) denotes the distances between the various important residues of the protein to the center of mass of the ligand.

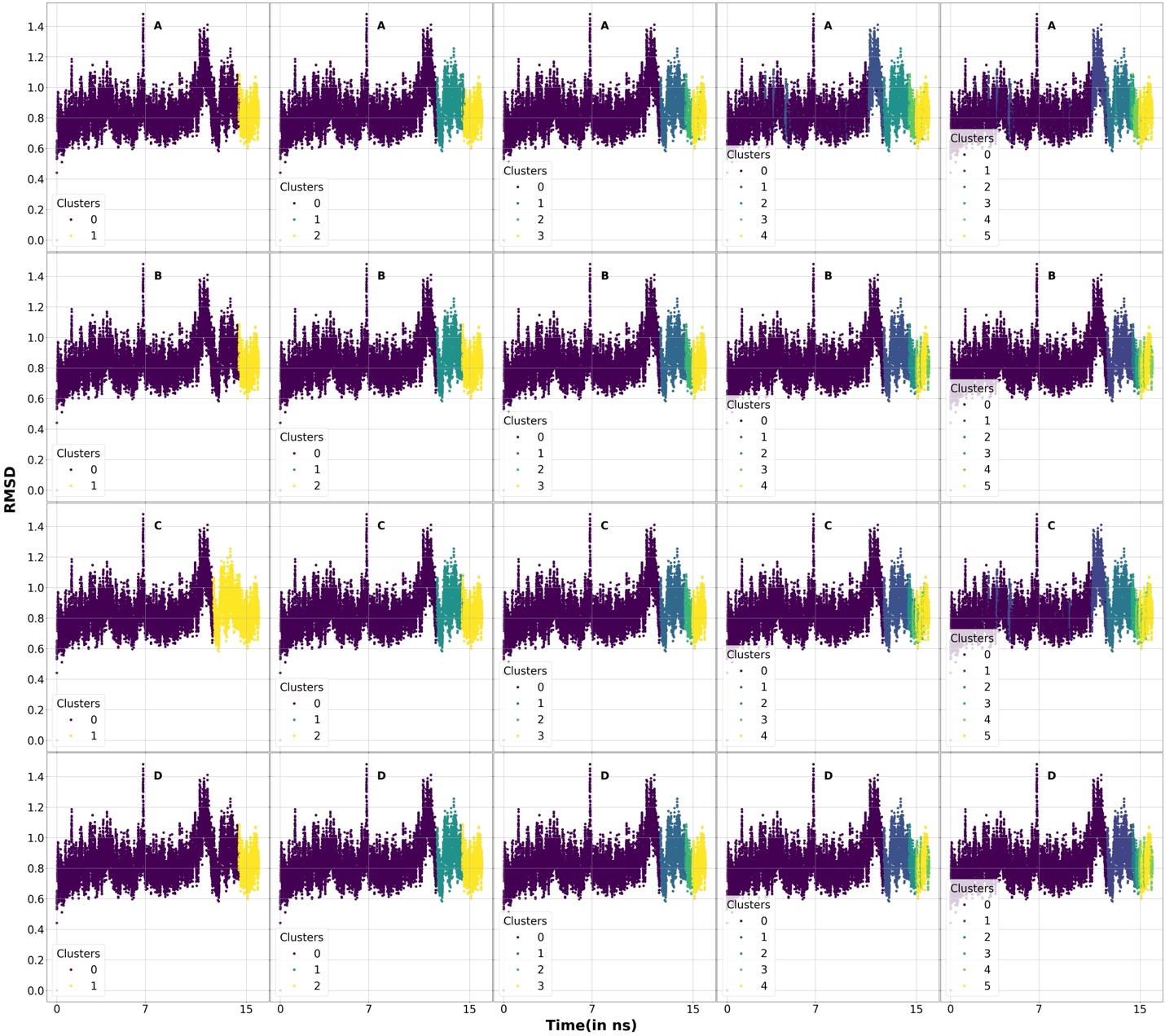

**Figure S38:** RMSD for the protein properties of the protein-ligand simulation trajectories used for clustering for FKBP protein bound/unbound to DMSO. Different clusters are marked using different colors. A, B, C, and D sets denotes our 4 clustering algorithms K-Means, Agglomerative, BIRCH, and Agglomerative. Each column respectively denotes 2,3,4,5, and 6 clusters.

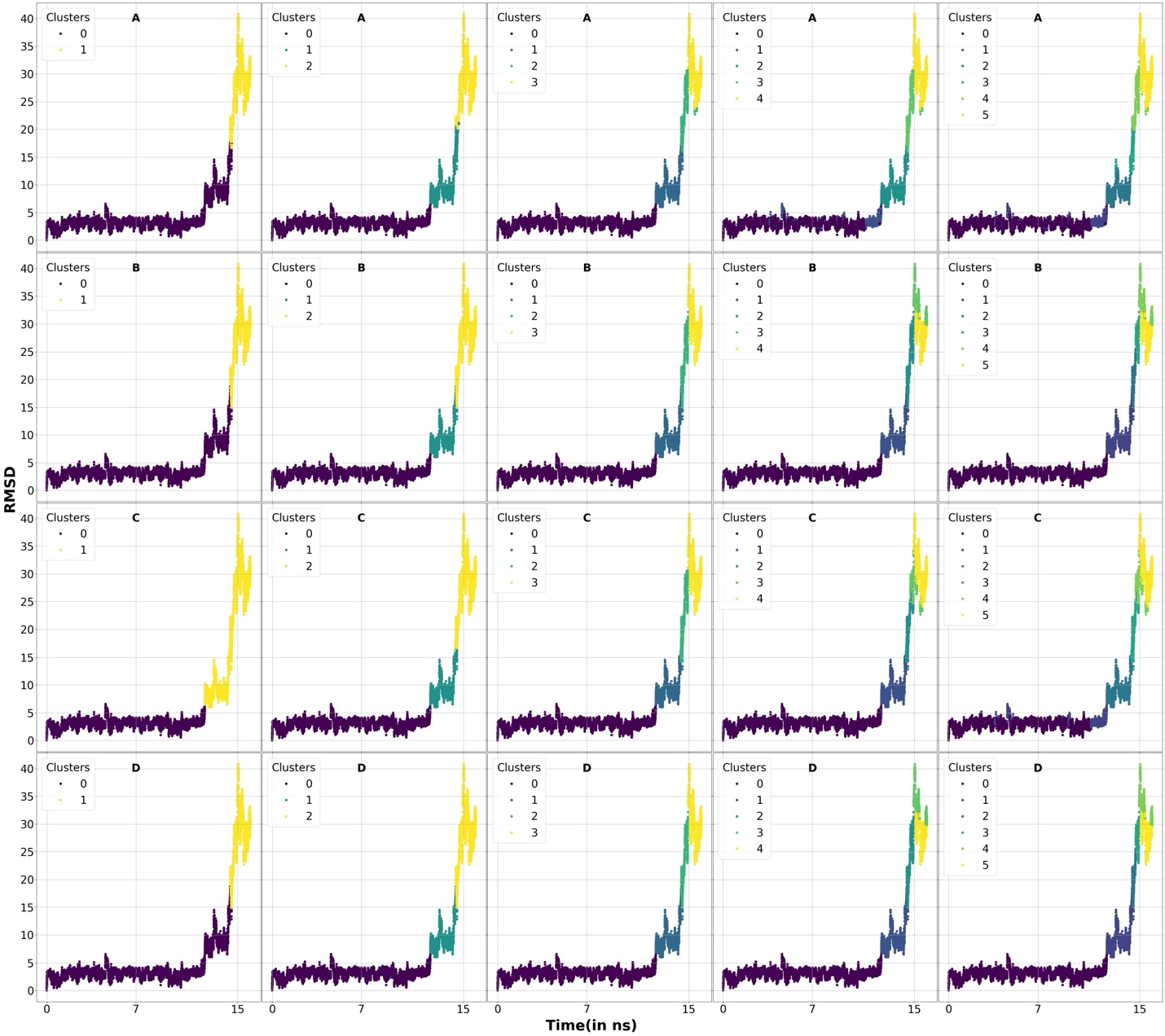

**Figure S39:** RMSD for the ligand properties of the protein-ligand simulation trajectories used for clustering for FKBP protein bound/unbound to DMSO. Different clusters are marked using different colors. A, B, C, and D sets denotes our 4 clustering algorithms K-Means, Agglomerative, BIRCH, and Agglomerative. Each column respectively denotes 2,3,4,5, and 6 clusters.

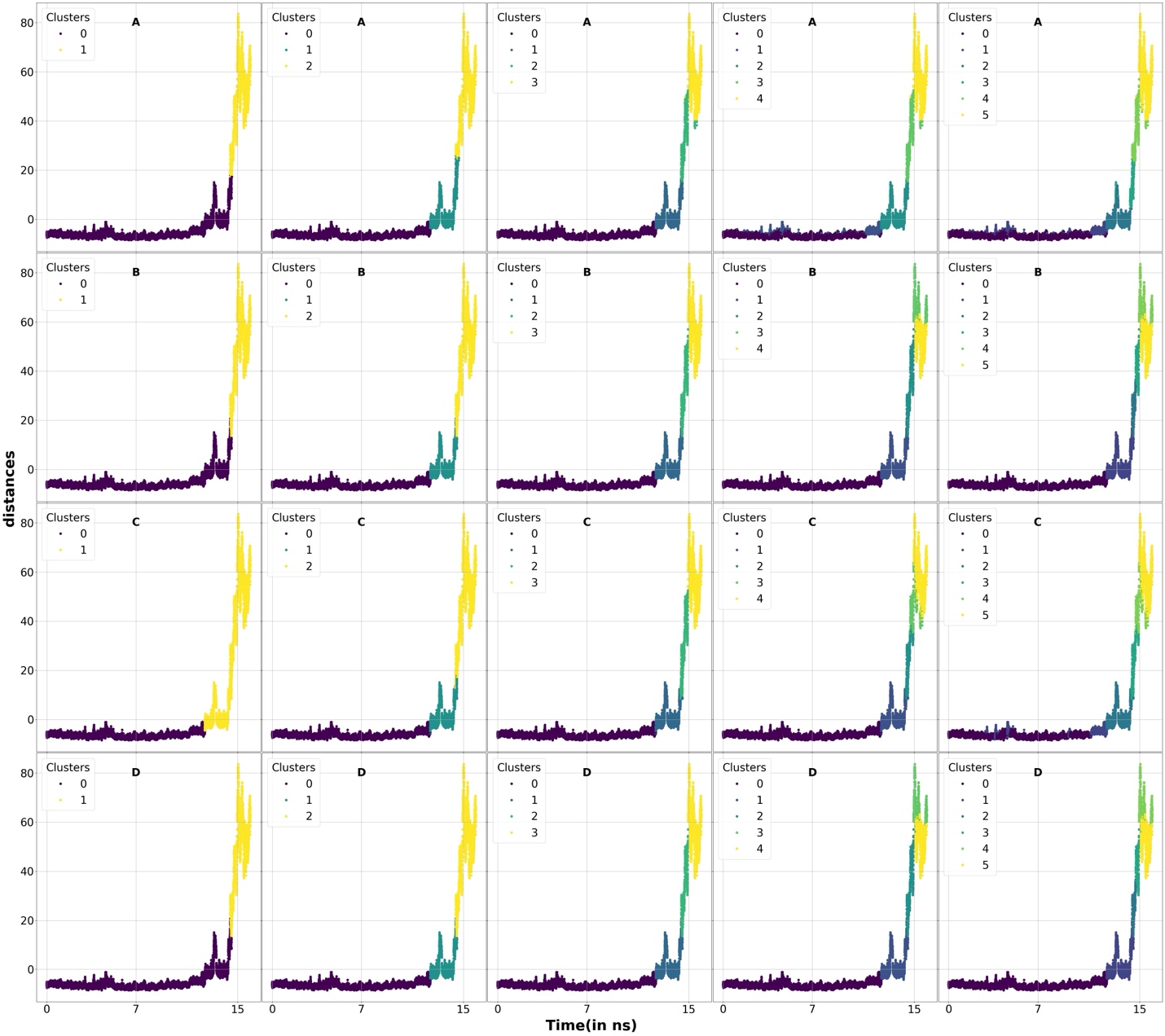

**Figure S40:** Reduced one-dimensional PCA component of protein-ligand distance properties of the protein-ligand simulation trajectories used for clustering for FKBP protein bound/unbound to DMSO. Different clusters are marked using different colors. A, B, C, and D sets denotes our 4 clustering algorithms K-Means, Agglomerative, BIRCH, and Agglomerative. Each column respectively denotes 2,3,4,5, and 6 clusters.

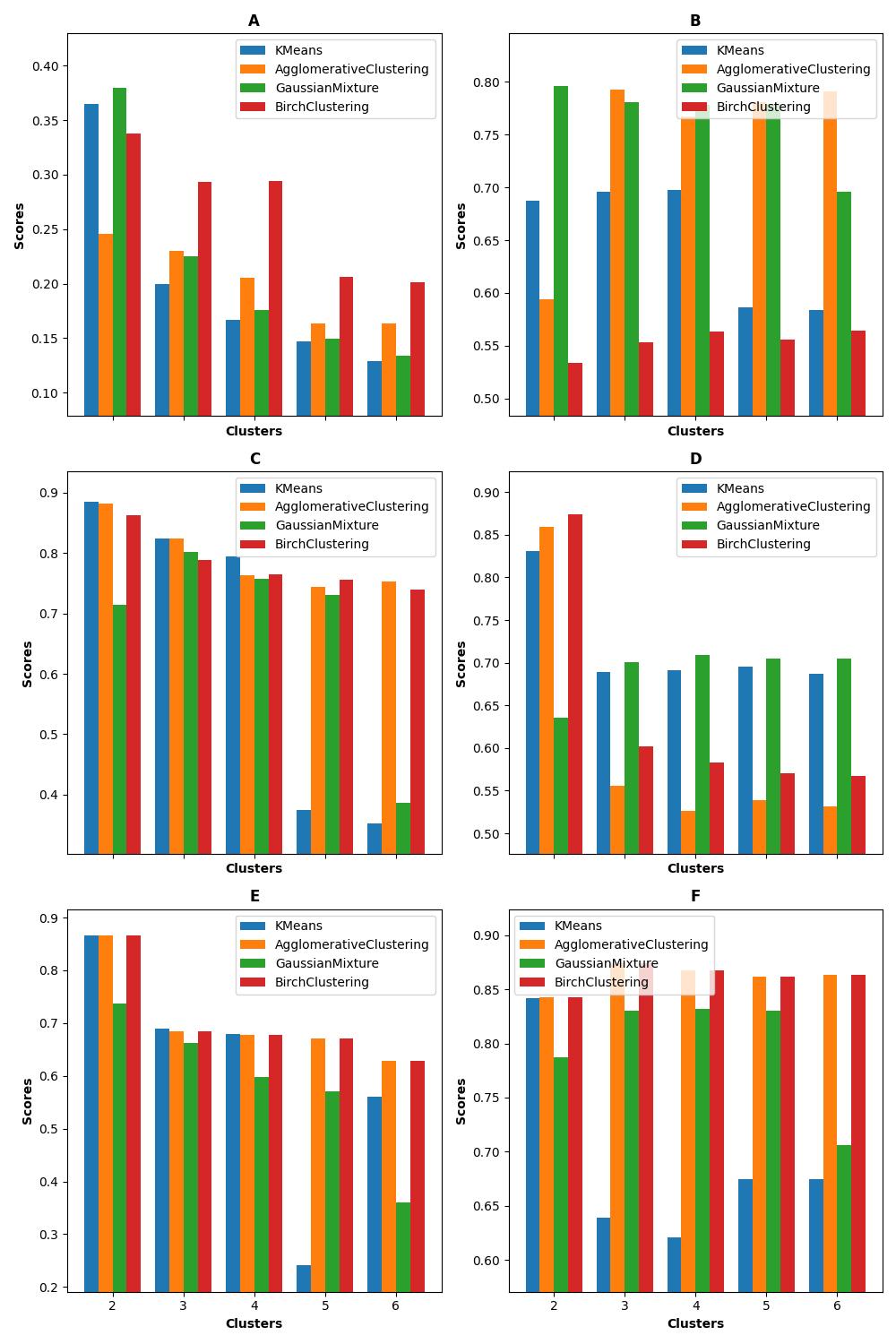

**Figure S41:** Scoring functions based on protein-ligand simulation trajectories used for clustering for FKBP protein bound/unbound to DMSO. (A) and (B) denotes the scoring functions-1 and 2 used on RMSD property for the backbone, (C) and (D) denotes the scoring functions-1 and 2 used on RMSD property of the sugar, (E) and (F) denotes the scoring functions-1 and 2 on the distances between the protein and the sugar.

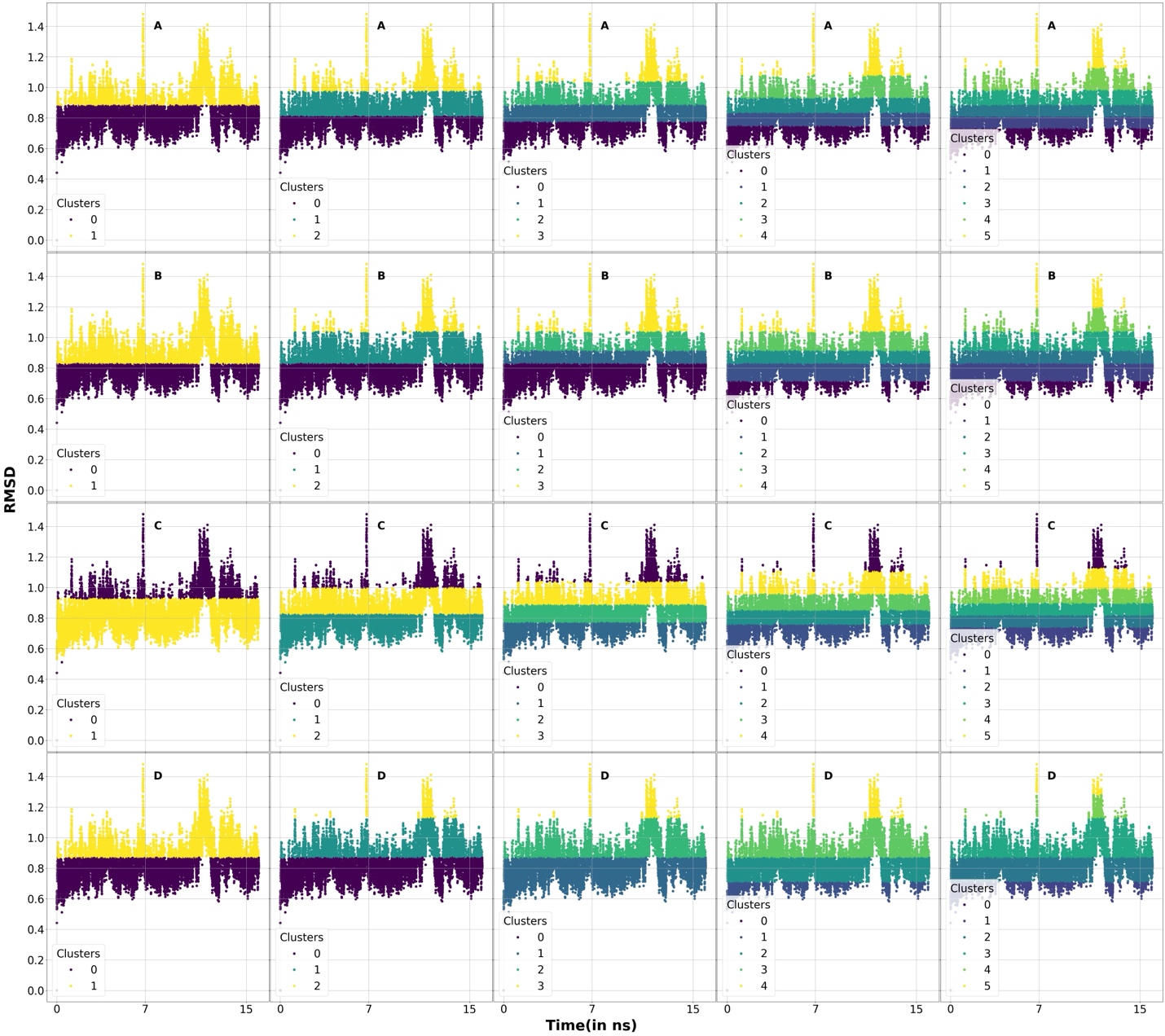

**Figure S42:** A graphical representation of the cluster states for the ground truth label for the RMSD of the protein backbone (for scoring metric-1) as a function of time for FKBP protein bound/unbound to DMSO. A, B, C, and D sets denotes our 4 clustering algorithms K-Means, Agglomerative, BIRCH, and Agglomerative. Each column respectively denotes 2,3,4,5, and 6 clusters.

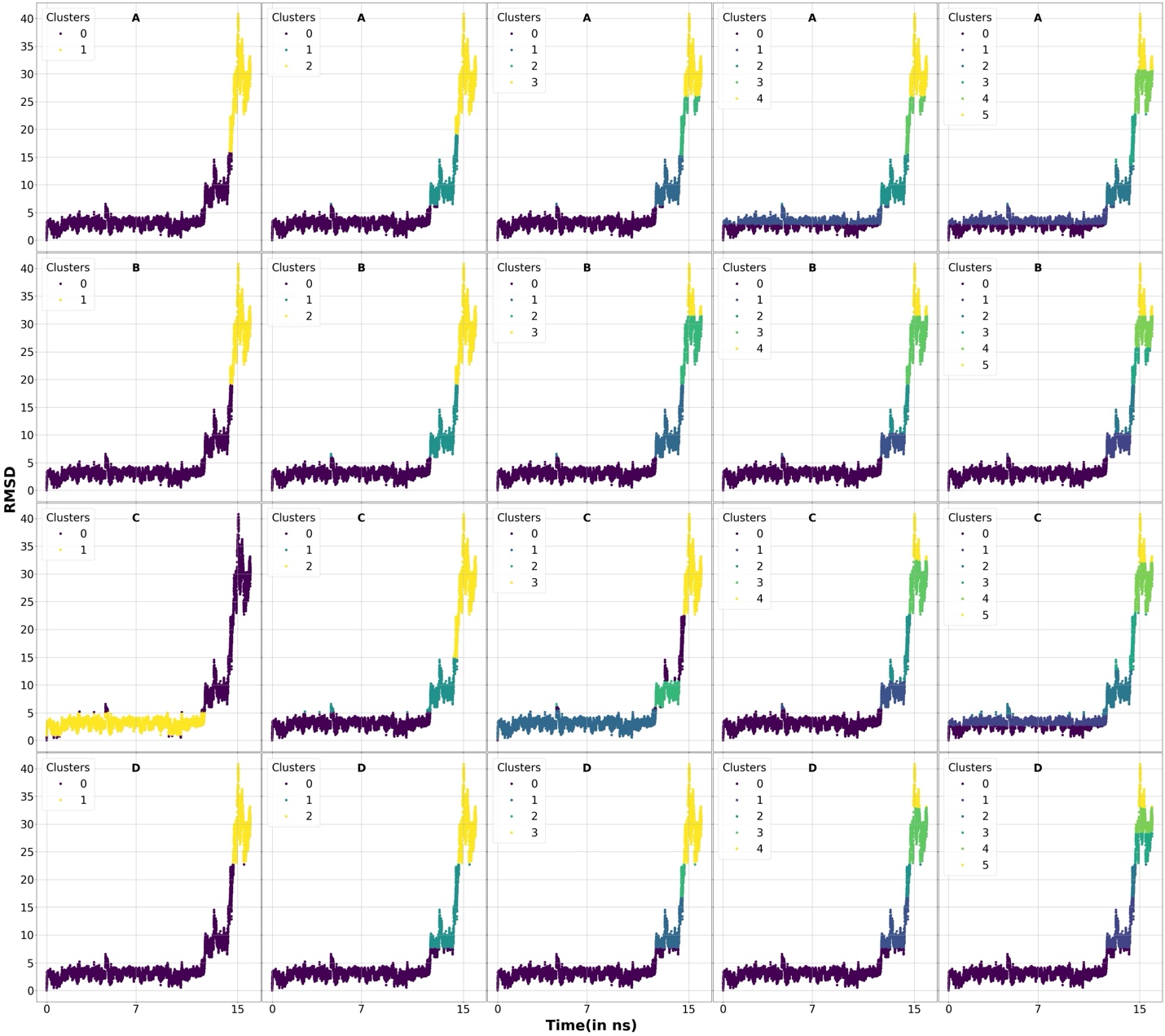

**Figure S43:** A graphical representation of the cluster states for the ground truth label for the RMSD of the sugar backbone (for scoring metric-1) as a function of time for FKBP protein bound/unbound to DMSO. A, B, C, and D sets denotes our 4 clustering algorithms K-Means, Agglomerative, BIRCH, and Agglomerative. Each column respectively denotes 2,3,4,5, and 6 clusters.

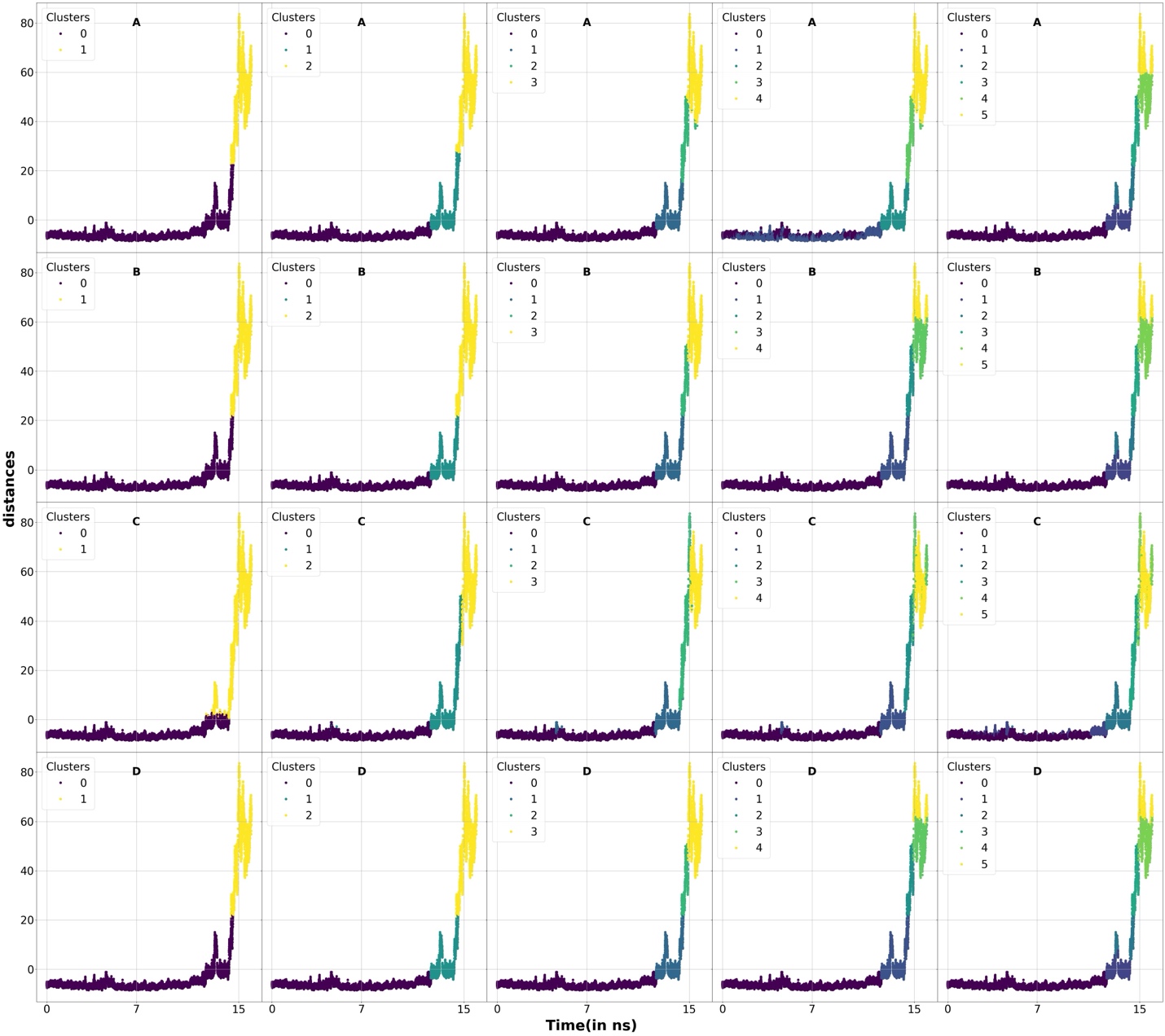

**Figure S44:** A graphical representation of the cluster states for the ground truth label for the distances of the protein to the sugar backbone (for scoring metric-1) as a function of time for FKBP protein bound/unbound to DMSO. A, B, C, and D sets denotes our 4 clustering algorithms K-Means, Agglomerative, BIRCH, and Agglomerative. Each column respectively denotes 2,3,4,5, and 6 clusters.

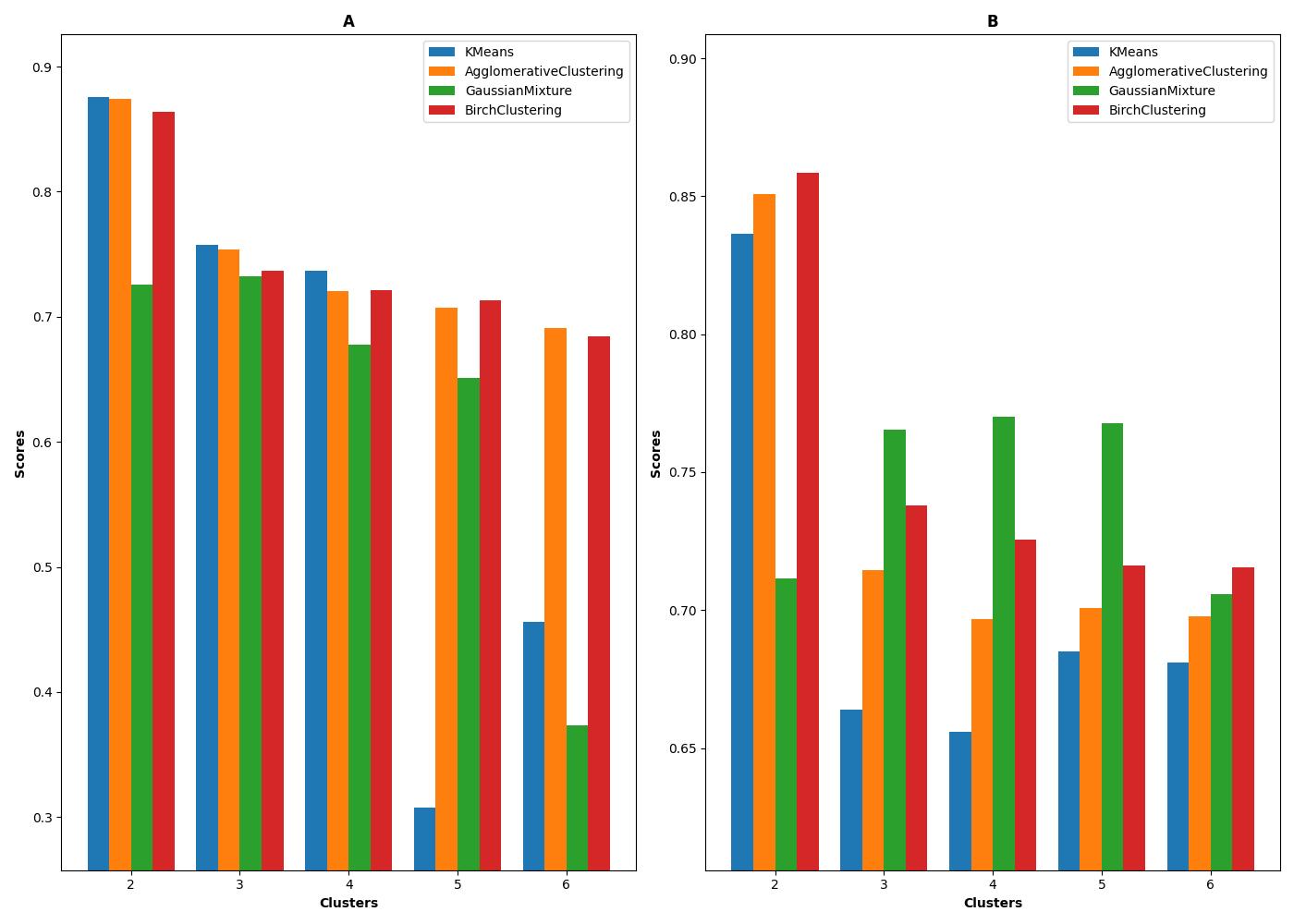

**Figure S45:** Overall scores based on our scoring function-1 (A) and scoring function-2 (B) based on the scores on RMSD of the sugar, distance properties of the sugar from the protein, as well as the presence and absence of hydrogen bonds/salt-bridges shown in figure 15 for the protein-ligand simulations for FKBP protein bound/unbound to DMSO. The weights W_rmsd-sugar_ and W_distances_, are set to 0.50 and 0.50 respectively.

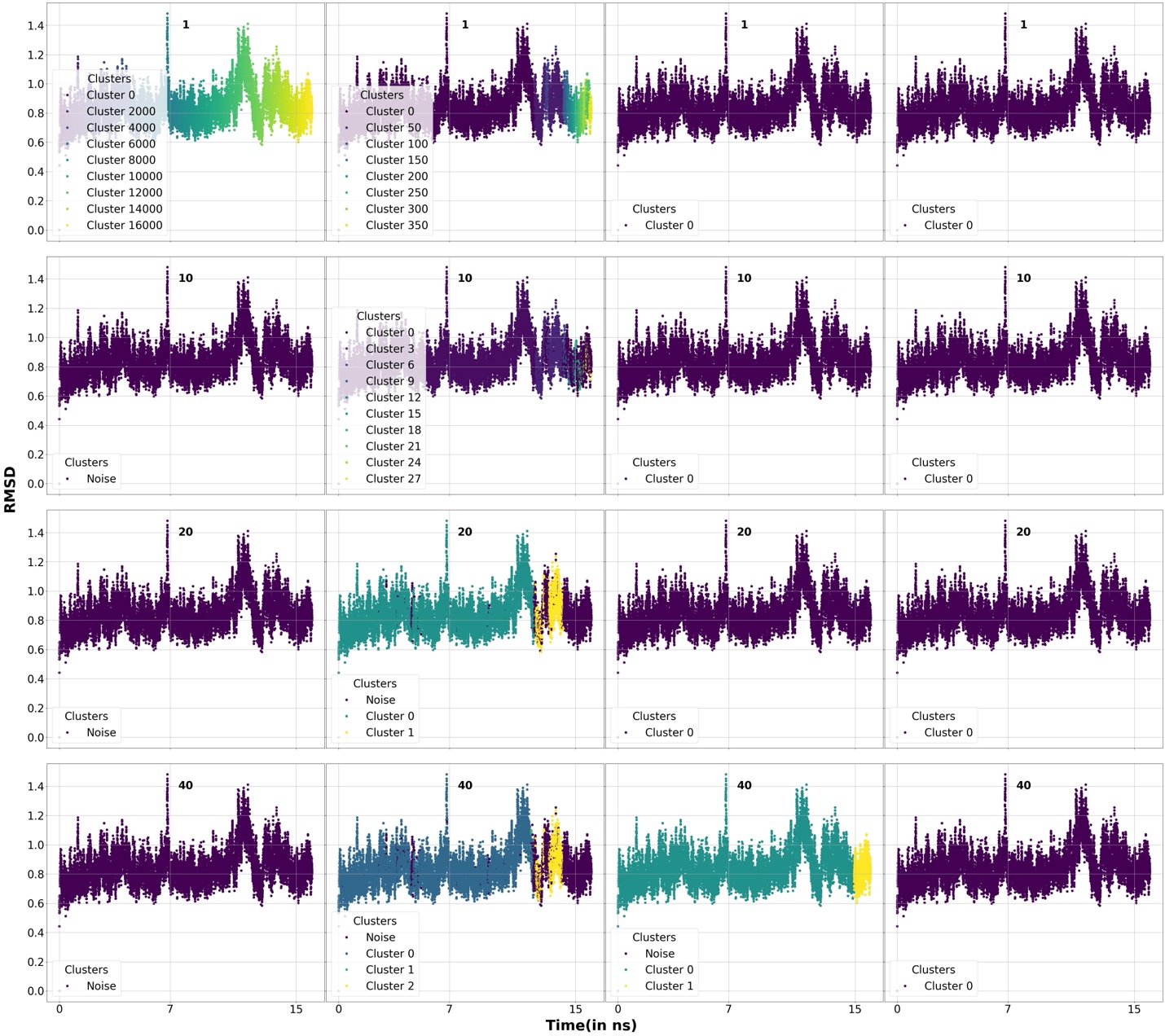

**Figure S46:** RMSD of the ligand properties of the different protein-ligand samples used for clustering by DBSCAN algorithm for the FKBP protein bound/unbound to DMSO. Different clusters as well as the unclustered points(noise) are marked using different colors. The min-points used for the clustering are shown in labels. Each row represents the min-points used in labels with increasing ε values=1.0, 5.0, 20.0, and 40.0.

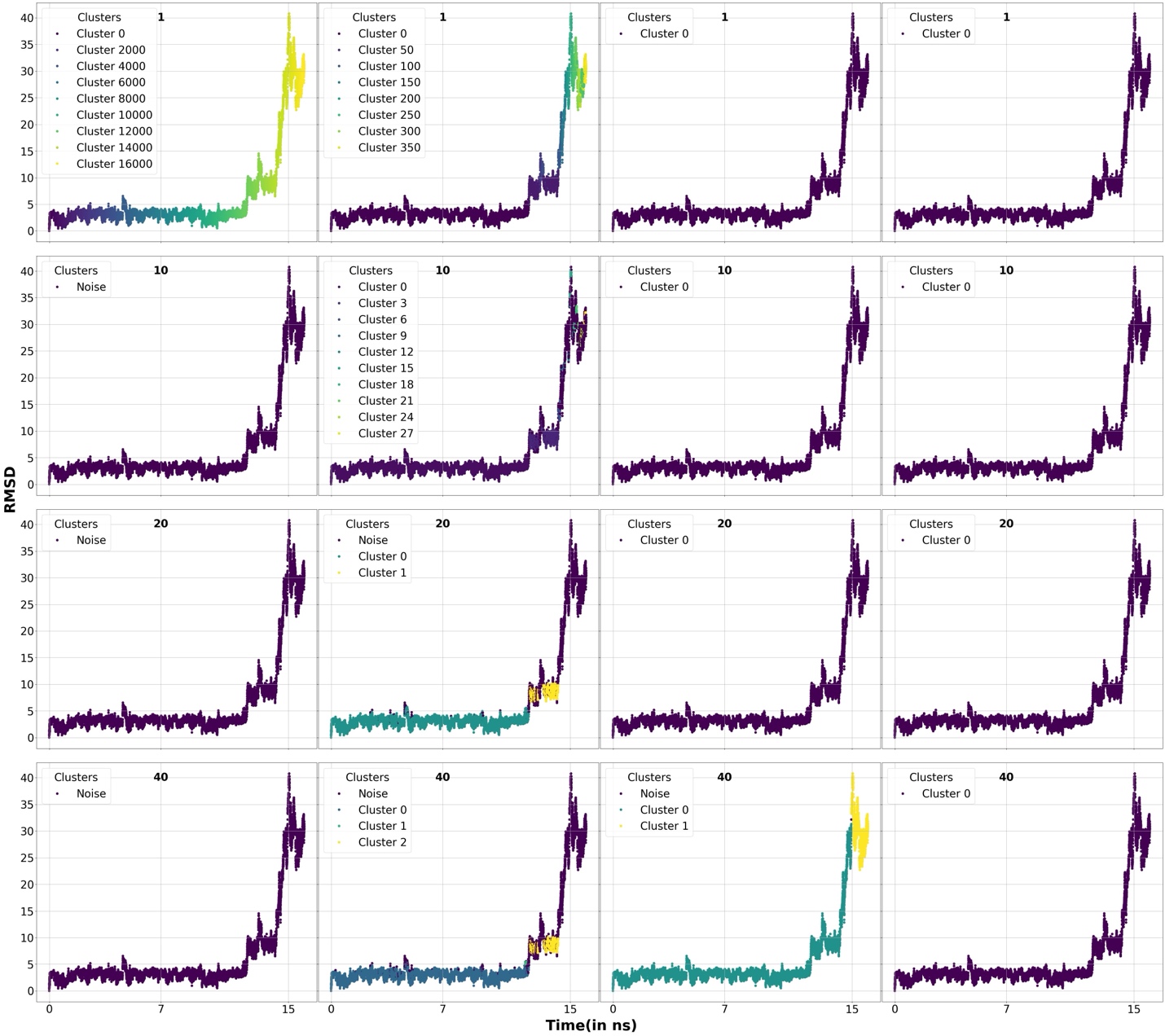

**Figure S47:** RMSD of the ligand properties of the different protein-ligand samples used for clustering by DBSCAN algorithm for the FKBP protein bound/unbound to DMSO. Different clusters as well as the unclustered points(noise) are marked using different colors. The min-points used for the clustering are shown in labels. Each row represents the min-points used in labels with increasing ε values=1.0, 5.0, 20.0, and 40.0.

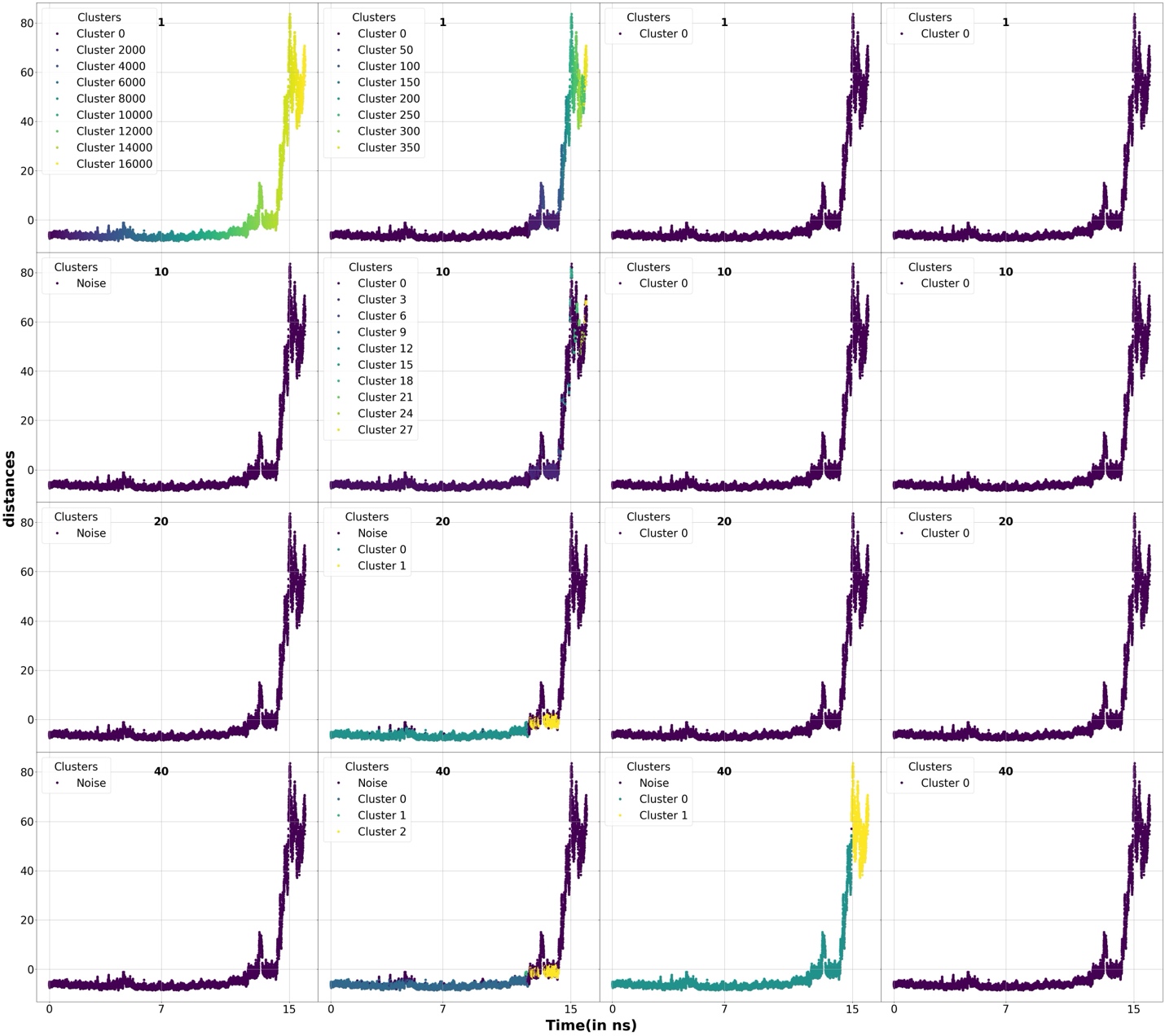

**Figure S48:** A dimensional PCA component of the distance properties of the different protein-ligand samples used for clustering by DBSCAN algorithm for the FKBP protein bound/unbound to DMSO. Different clusters as well as the unclustered points(noise) are marked using different colors. The min-points used for the clustering are shown in labels. Each row represents the min-points used in labels with increasing ε values=1.0, 5.0, 20.0, and 40.0.

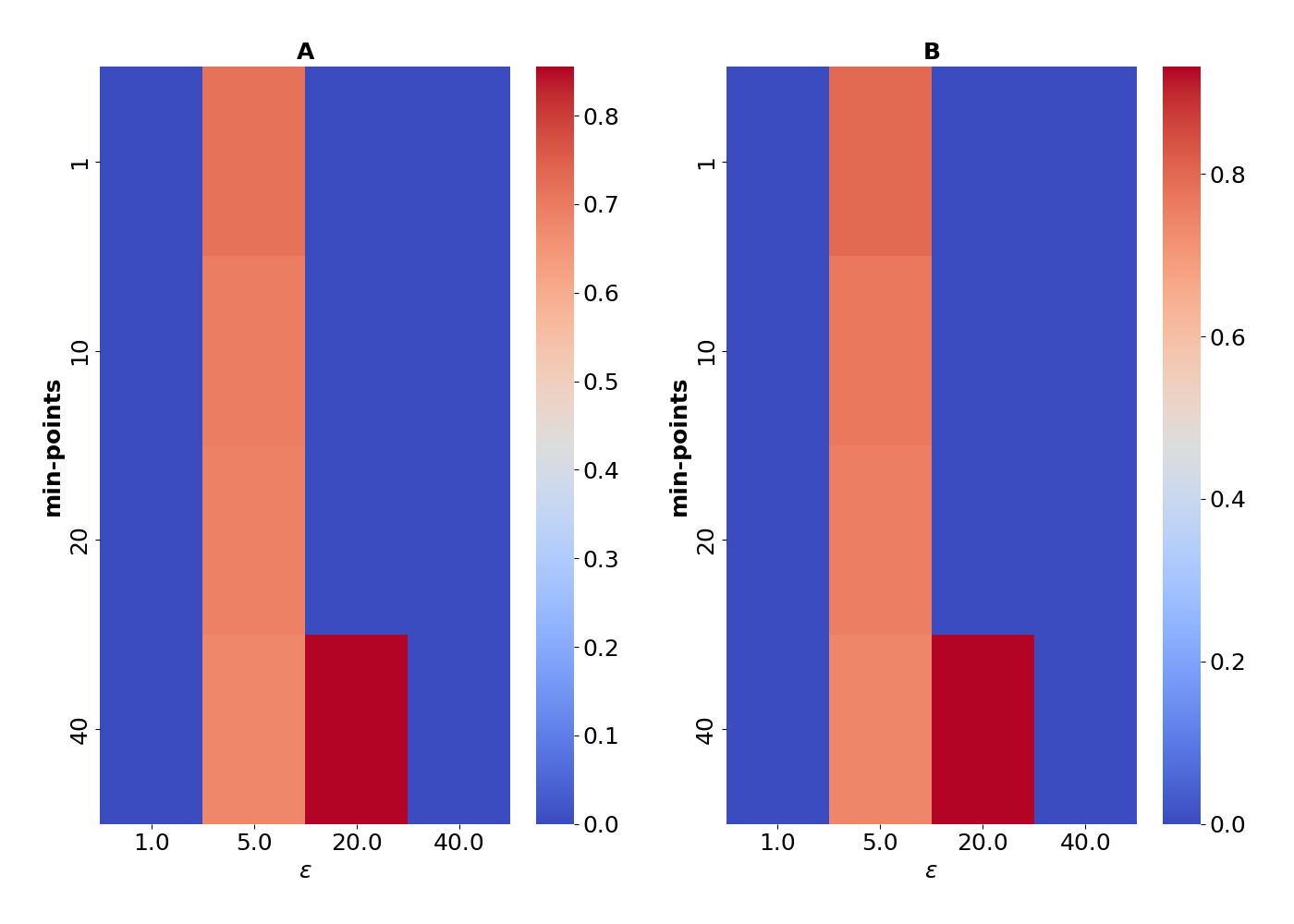

**Figure S49:** A two dimensional heatmap showing the scores obtained by our scoring metric-1(A) and scoring metric-2(B) for DBSCAN algorithm obtained by setting a weights of W_rmsd-sugar_=0.50 and W_distances_=0.50 to our overall scoring function(Equation-1 in the main manuscript) for FKBP protein bound/unbound to DMSO.

**References**

^1^ A. C. Pan *et al.*, Journal of Chemical Theory and Computation **13** (2017) 3372.

^2^ D. Pramanik *et al.*, The Journal of Physical Chemistry B **123** (2019) 3672.

^3^ S. Lee *et al.*, Journal of Chemical Theory and Computation **20** (2024) 6341.

^4^ K. Mondal *et al.*, bioRxiv (2024) 2024.09.24.614745.
